# Mycothiol Conjugation Drives Biotransformation and Biodefluorination of Fluorotelomer Carboxylic Acids by *Actinomycetota* and Sludge Microbiomes

**DOI:** 10.1101/2025.09.24.678248

**Authors:** Boyuan Su, Hao Chen, Mengyan Li

**Affiliations:** Department of Chemistry and Environmental Science, New Jersey Institute of Technology, Newark, New Jersey, United States

**Keywords:** PFAS, FTCA, Biotransformation, Biodefluorination, Mycothiol, *Actinomycetota*, Non-target Analysis, Thiol Conjugation

## Abstract

Microbial breakdown of poly- and perfluoroalkyl substances (PFASs) remains poorly understood, often hindered by mass balance discrepancies and unexplained fluoride release. Here we identify a previously unrecognized PFAS biodefluorination mechanism mediated by mycothiol (MSH), a ubiquitous thiol antioxidant produced by *Actinomycetota*. Using high-resolution mass spectrometry and non-target analysis, MSH-mediated conjugation is unraveled as a dominant biotransformation pathway for two representative fluorotelomer carboxylic acids (6:2 and 5:3 FTCAs) in *Rhodococcus jostii* RHA1 and other MSH-producing species. Conjugation proceeds through nucleophilic substitution and β-elimination in 6:2 FTCA, releasing two fluoride ions, and through Michael addition in the unsaturated analogue of 5:3 FTCA. These reactions are catalyzed by the conserved enzyme mycothiol S-transferase (MST), which contains a conserved His47–Asp150–His154 tripod motif essential to activate MSH for nucleophilic attack. Subsequent hydrolysis by mycothiol S-conjugate amidase (Mca) yields mercapturic acid derivatives, with more than 20 novel downstream thiol metabolites detected. Time-series experiments confirmed that MSH conjugation accounted for up to half of FTCA removal and fluoride release, resolving major fluorine mass balance gaps. The conjugation process also interlinks with chain-shortening “one-carbon removal” routes, forming a cascading metabolic network. Importantly, conjugated metabolites were consistently detected in activated sludge microbiomes, and metagenomic surveys revealed more than 100,000 MST homologs worldwide. Collectively, these findings establish MSH-mediated conjugation as a widespread and mechanistically distinct route of PFAS biodefluorination, highlighting new opportunities for microbial PFAS removal in natural and engineered systems.

## 1. Introduction

Per- and polyfluoroalkyl substances (PFASs) have attracted escalating concern due to their widespread use, bioaccumulation potential, environmental persistence, and adverse health effects [1–4]. These synthetic chemicals are used extensively in industrial applications and consumer products, including aqueous film-forming foams (AFFFs), food packaging, and textiles, attributed to their hydrophobic and lipophobic properties [5–7]. Despite thousands of PFASs being identified [8], only a limited number are strictly regulated [9, 10]. PFAS exposure has been linked to endocrine disruption, liver damage, elevated cholesterol, and developmental effects in humans, alongside ecological risks to wildlife [11, 12]. Consequently, understanding the fate and transformation of PFASs is of paramount importance to promote the development of effective strategies for PFAS contamination control and mitigation.

Microbial biodegradation of heavily fluorinated PFASs remains challenging, primarily due to the exceptional stability of C–F bonds, among the strongest in organic chemistry with a bond dissociation energy of 544 kJ/mol (Table S1) [13]. However, recent studies have demonstrated microbial-mediated biodefluorination in both pure cultures and mixed consortia [14, 15]. Under anaerobic conditions, caffeoyl-CoA reductase has been implicated for reductive defluorination of α, β-unsaturated perfluorinated carboxylic acids by *Acetobacteria* [16]. Aerobic biotransformation of fluorotelomer-based PFASs has been proposed to proceed via sequential “one-carbon removal pathways”, in which CF2 units are cleaved from the fluorinated chain [17]. The prevalence of “one-carbon removal pathways” is evident from the consistent detection of key metabolites in activated sludge-based investigations [17–20] . For instance, 5:3 fluorotelomer carboxylic acid (5:3 FTCA) can undergo transformation into 4:3 FTCA and shorter chain perfluorocarboxylic acids (PFCAs), such as PFPeA and PFBA, via α-hydroxyanionoxylation and sequential decarboxylation. However, substantial mass balance discrepancies observed in these studies suggest the involvement of additional, yet unidentified, pathways.

Emerging evidence highlights the pivotal role of low-molecular-weight (LMW) thiols in PFAS biotransformation. Glutathione (GSH) has been reported for its ability to conjugate with fluorotelomer alcohols (FTOHs), fluorotelomer carboxylic acids (FTCAs), and fluorotelomer unsaturated carboxylic acids (FTUCAs) at β-carbons or terminal moieties (Table S2) [16, 21–27]. An alternative conjugative pathway that is analogous to fatty acid β-oxidation has recently been proposed based on the formation of Coenzyme A (CoA) adducts for short-chain FTCAs with non-fluorinated moieties [28]. In *Gordonia* sp. NB4-1Y, a medium-chain acyl-CoA synthetase was found capable of conjugating 2:3 FTCA, 2:3 FTUCA, and 1:4 FTCA, but not other longer-chain FTCAs [28]. These studies revealed the involvement of LMW thiols in PFAS biotransformation (Table S2). However, the underlying biochemical thiol-mediated mechanisms remain poorly understood, and their environmental relevance has yet to be fully established.

This study investigates FTCA biotransformation by four archetypical *Actinomycetota*species and activated sludge microbiomes to elucidate dominant pathways and the associated enzymatic processes. 6:2 and 5:3 FTCAs were selected as model substrates (Table S3), given their roles as key intermediates during PFAS biotransformation [29, 30] and their widespread detection in landfill leachates and other environmental matrixes [31–34]. These compounds also exhibit significantly greater toxicity and bioaccumulation potential than PFOA, their perfluorinated counterpart with identical chain length (C8) [35, 36]. As depicted in Figure 1a, a nano-electrospray ionization high-resolution mass spectrometry (Nano-ESI HRMS, Figure S3) method previously developed in our laboratory [34] was employed for both target and non-target analyses to monitor FTCA degradation and identify major transformation products (TPs).

**Figure 1.**
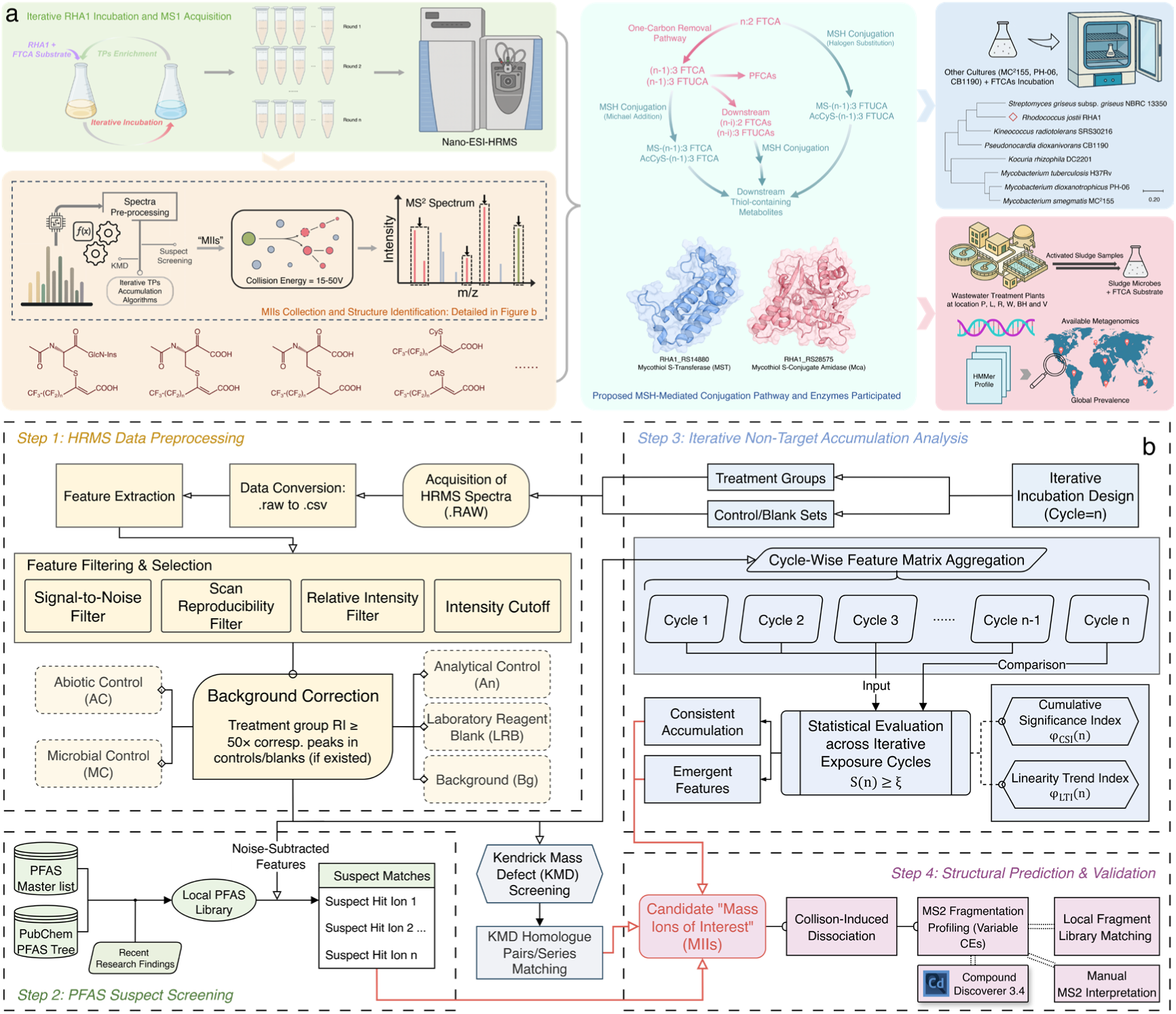
Flowcharts illustrating (a) the overall research workflow and (b) the non-target analysis algorithms for the screening and identification of putative TPs.

High-throughput screening algorithms (Figure 1b) facilitated the identification of novel TPs, and collision-induced dissociation (CID) provided structural insights based on their fragmentation patterns. Fluorine-based mass balance calculations enabled quantitative assessment of the divergent biotransformation pathways. Enzymes initiating the newly discovered pathways were identified and their catalytic functions validated through AlphaFold2-based structural homology analysis with model enzymes. Furthermore, metagenomic database mining revealed their environmental prevalence at the global scale. These findings unveil previously unrecognized PFAS biotransformation mechanisms and their widespread distribution in wastewater and aquatic environments.

## 2. Results & Discussion

### 2.1 Detection of Metabolites Conjugated with Mycothiol and Acetyl-L-cysteine in Rhodococcus jostii *RHA1*

For the enrichment of TPs of FTCAs, *Rhodococcus jostii* RHA1 was selected as a model *Actinomycetota* species given its superior ability to biotransform FT-based PFASs, including 6:2 FTS [37] and 6:2 FTOH [38], for which 6:2 and 5:3 FTCAs were identified as important intermediates. RHA1 is also known as a powerhouse of catabolic genes, enabling its degradation of a wide spectrum of xenobiotics [39]. Consistent with previous studies, RHA1 exhibited conceivably high biodefluorination of 6:2 FTCA in resting cell assays compared to 15 other Gram-positive and Gram-negative bacteria tested (Figure S1), including *Pseudomonas putida* F1 [40] and *Rhodococcus opacus* 1cp [41], which were reported to defluorinate fluorinated ethers and phenols, respectively. In this study, iterative 48-h FTCA exposures of RHA1 resting cells were conducted, with the spent media from each cycle were reused for the following exposure cycle to enrich TPs (Figure 2; Figure S2 and S4).

**Figure 2.**
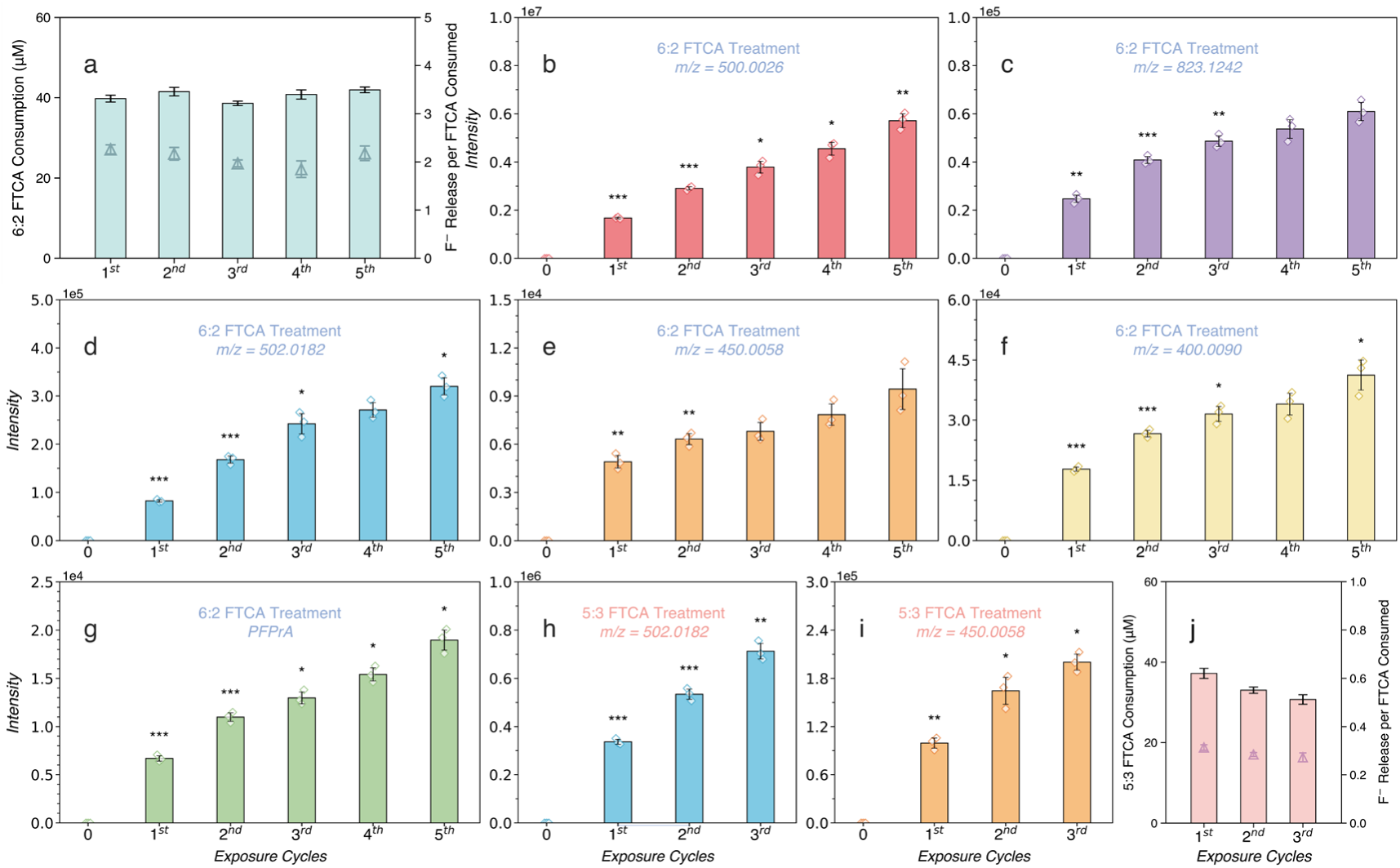
Iterative exposure enriches TPs from FTCA biotransformation and biodefluorination by *Rhodococcus jostii* RHA1. Disappearance of 6:2 FTCA (a) and 5:3 FTCA (j) with corresponding release of fluoride for each exposure cycle. Accumulation of non-target TPs over 5 cycles of exposure to 6:2 FTCA (b-g) and 3 cycles of exposure to 5:3 FTCA (h-i). Error bars represent the standard deviation of triplicate experiments. Asterisks indicate statistically significant increase in intensity compared to the previous cycle (\**p* < 0.05; \*\**p* < 0.01; \*\*\**p* < 0.001).

As the disappearance of 6:2 FTCA and the liberation of free fluoride (Figure 2a), “mass ions of interest” (MIIs) were screened as candidate TPs given their continuous enrichment through the iterative exposure cycles based on an indicator-based statistics (Figure 1b; SI Section S1.4). In addition to the detection of less fluorinated FTCAs/FTUCAs (e.g., 5:3 FTCA,5:3 FTUCA, OH-5:3 FTCA, 4:2 FTUCA) and shorter-chain PFCAs (e.g., PFHxA, PFPeA) (Figure S4d-i), three MIIs at *m/z* 500.0026, 823.1242, and 502.0182 (Figure 2b-d) were observed as major non-target metabolites, which exhibited significantly greater intensities when compared to prior cycles. Absent in corresponding controls and blanks, these three MIIs are larger than the parent 6:2 FTCA ion (*m/z* 376.9847, Table S3), suggesting possible formation of conjugated metabolites.

The MII at *m/z* 500.0026 was consistently detected at high intensities, accumulating to >5e6 by the 5^th^ enrichment cycle (Figure 2b). Its MS2 spectrum generated via CID confirmed it as a thiol conjugated PFAS (Figure 3a; Figure S5). A series of fully or partially fluorinated alkyl fragments (marked in orange and green, respectively), such as C2F5^-^ (*m/z* 118.9915), C4F7^-^ (*m/z* 180.9887), and C F ^-^ (*m/z* 292.9831), were detected. Sulfur incorporation was revealed from several C8-thiolated fragments (highlighted in red): *C8H2-xF11-xO2S^-^ (x=0,1,2, m/z* 370.9606, 350.9544, 330.9482). The latter two could arise from HF eliminations from the parent *C8H2F11O2S^-^* fragment. Additional fluorinated alkyl thiolate fragments, *CxFyS^-^* (*x=6,7; y=6-9*) at *m/z* 217.9627, 236.9613, and 286.9583, indicate the thiol conjugation initiates at the β-carbon rather than the terminal carboxylate of 6:2 FTCA. A diagnostic fragment at *m/z* 162.0222 (*C5H8NO3S^-^*) corresponds to an N-acetyl-L-cysteine (AcCyS) moiety (Figure 3e; Figure S8). Together, these structural evidence reveals this dominant metabolite as AcCyS-5:3 FTUCA, a cysteinyl thiol conjugate with a 5:3 FTUCA backbone.

**Figure 3.**
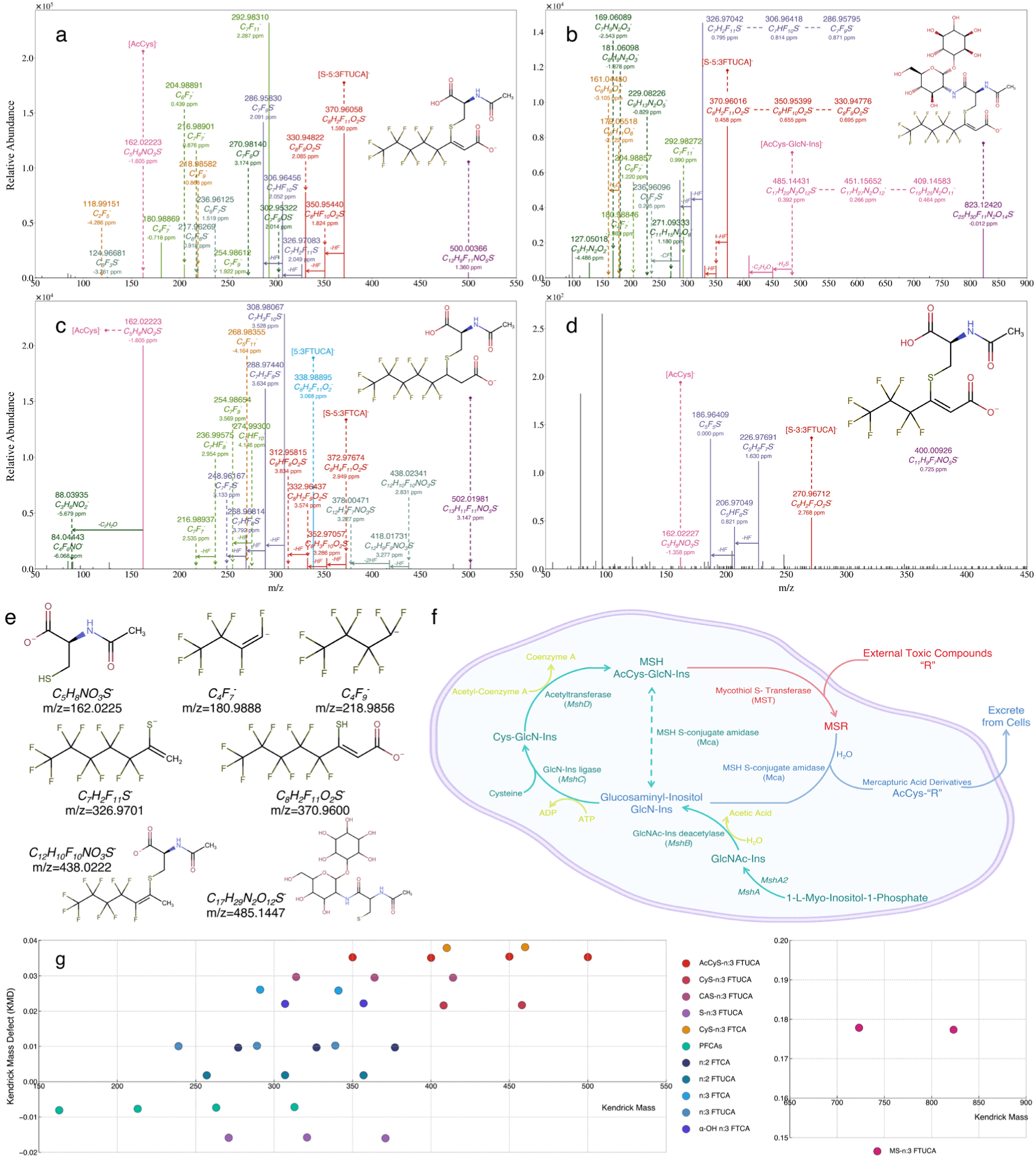
CID fragmentation spectra of four dominant TPs detected in iterative enrichment by RHA1, including (a) AcCyS-5:3 FTUCA (*m/z* 500.0026), (b) MS-5:3 FTUCA (*m/z* 823.1242), (c) AcCyS-5:3 FTCA (*m/z* 502.0182), and (d) AcCyS-3:3 FTUCA (*m/z* 400.0090). (e) Structures and *m/z* values of key fragments identified among conjugated TPs. (f) Proposed MSH-mediated conjugation mechanisms when FTCAs or other external toxic compounds “R” are introduced to an *Actinomycetota* cell. (g) KMD (CF2) map of TPs from iterative 6:2 FTCA exposure, revealing homolog series of conjugated TPs and TPs from “one-carbon removal pathways”.

The second MII at *m/z* 823.1242 also accumulated over enrichment cycles, though at intensities ∼2 orders of magnitude lower than AcCyS-5:3 FTUCA (Figure 2c). The 323.1216 Da mass difference between two MIIs matches the dehydrated mass of 1D-*myo*-inositol 2-ammonio-2-deoxy-alpha-D-glucopyranoside (GlcN-Ins). AcCyS and GlcN-Ins couple via an amide bond to form mycothiol (MSH, AcCyS-GlcN-Ins, Figure S8), a major LMW thiol antioxidant synthesized at millimolar concentrations by *R. jostii* RHA1 and other *Actinomycetota* [42, 43]. CID of this second MII (Figure 3b; Figure S6) yielded fragments at *m/z* 485.1443 and *m/z* 451.1565, corresponding to the mycothiolate ion (MS^-^; Figure 3e) and a secondary fragment from the neutral loss of hydrogen sulfide, respectively. C8 and C7 fluorinated thiolate tail fragments, C8H2F11O2S^-^ (*m/z* 370.9602) and C7H2F11S^-^ (*m/z* 326.9704), further confirmed the structure. Collectively, these fragmentation features corroborate this MII as the direct MSH conjugate, MS-5:3 FTUCA. As shown in Figure 3f, MS-5:3 FTUCA can be further converted to AcCyS-5:3 FTUCA, since the amide linking AcCys and GlcN-Ins in MS-conjugates can be hydrolyzed by the enzyme mycothiol S-conjugate amidase (Mca), liberating mercapturic acid derivatives (AcCyS-R) [42, 44]. This explains the coupling production of MS-5:3 FTUCA and AcCyS-5:3 FTUCA.

Under identical exposure conditions, RHA1 did not fully consume equimolar dose (40 µM) of 5:3 FTCA, leaving ∼6 % residual substrate (Figure 2j). Though the majority of the 5:3 FTCA was biotransformed, the average fluoride release was at 9.80 ± 0.43 µM per cycle, equivalent to ∼0.30 mol of F^-^ release per mole of 5:3 FTCA removal. Non-target screening revealed a major MII at *m/z* 502.0182 arose from the 5:3 FTCA treatment (Figure 2h). Its MS2 fragmentation profile (Figure 3c; Figure S7) contains a fragment set similar to that of AcCyS-5:3 FTUCA, including AcCyS moiety, thiolate fluorotelomer, and fluorinated alkyl fragments. A +2.0156 Da mass shift matches the addition of two hydrogens as the saturation at the α, β-double bond of the 5:3 FTUCA moiety, indicating this MII as the reduced analogue AcCyS-5:3 FTCA. We propose this saturated mercapturic derivative arises from the Mca-mediated hydrolysis of its corresponding MSH conjugate, MS-5:3 FTCA [45]. The detection of AcCyS-5:3 FTUCA and AcCyS-5:3 FTCA supports an MSH-mediated conjugation pathway for 6:2 and 5:3 FTCA biotransformation by RHA1. AcCyS-5:3 FTCA also accumulated during the 6:2 FTCA-exposed assays, albeit at 3.0 to 4.1-fold lower intensities (Figure 2d), suggesting its secondary formation once 5:3 FTCA emerges during 6:2 FTCA biotransformation (see further explanations below).

### 2.2 MSH-mediated FTCA Biotransformation Pathways

Recent studies have reported the generation of GSH conjugates (Table S2) with fluorotelomer alcohols (e.g., 8:2 and 6:2 FTOHs) in bacteria [23], fungi [21], plants [22], and animals [25–27]. Functionally analogous to GSH, RHA1 and other *Actinomycetota* are known to synthesize glucosamine-inositol thiol (i.e., MSH) to perform similar antioxidant roles. Its reduced thiol (-SH) quenches reactive species, maintains cellular redox balance, and mitigates oxidative stress [46].

Here we reported MSH as a significant mediator in FTCA biotransformation and biodefluorination. Non-target analysis revealed the accumulation of major MSH-derived conjugates, MS-5:3 FTUCA and its mercapturic acid derivative, AcCyS-5:3 FTUCA, during exposure to 6:2 FTCA, with AcCyS-5:3 FTCA appeared in both 6:2 and 5:3 FTCA treatments. Formation of these conjugates can be explained by an MSH-mediated antioxidant framework, where MSH first conjugates with thiol-reactive xenobiotics, and the resulting MS-conjugates are further hydrolyzed to form AcCyS-conjugates (i.e., mercapturic acids) [42]. Once 6:2 FTCA enters the cells as a xenobiotic, the nucleophilic thiolate of MSH attacks the activated β-CF2 unit. This triggers a concurrent SN2 nucleophilic substitution, displacing one fluorine at the β-carbon, and an E2 elimination, expelling HF, to generate MS-5:3 FTUCA (Figure 4). These reactions lead to the liberation of two fluoride ions per 6:2 FTCA molecule conjugated.

**Figure 4.**
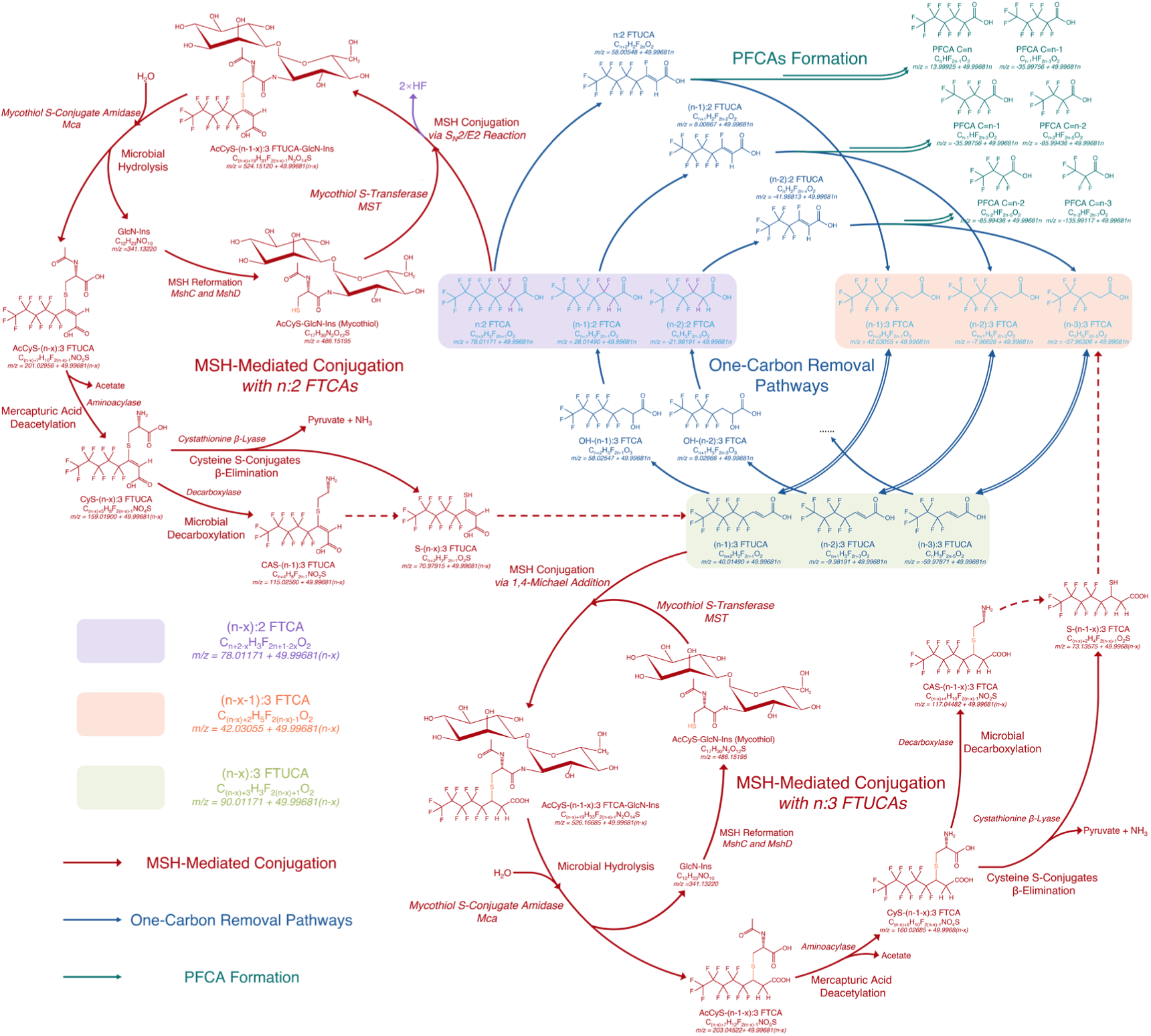
Proposed cascading biotransformation network that links two MSH-mediated conjugation pathways (red) with the “one-carbon removal pathways” (blue) that results in PFCA formation (green) when n:2 and n:3 FTCAs (in purple and orange shadows, respectively) are present in the microbial system.

As depicted in Figure 3f, this conjugation step can be catalyzed by a mycothiol S-transferase (MST). The amide bond linking GlcN-Ins and cysteine part within the MS-5:3 FTUCA conjugate is then hydrolyzed by Mca. This cleavage removes the toxic moieties and results in a rapid liberation of the mercapturic acid derivative AcCyS-5:3 FTUCA from cells. The non-toxic metabolite GlcN-Ins remains within the cell and recycles for the biosynthesis of new MSH, catalyzed by L-cysteine:1D-*myo*-inositol 2-amino-2-deoxy-alpha-D-glucopyranoside ligase (MshC) and MSH acetyltransferase (MshD) [42]. The FTCA detoxification mechanism, consisting of MSH conjugation, exportation of toxic conjugates, and MSH re-synthesis are depicted in Figure 3f and SI (Section S2.1).

5:3 FTCA lacks a fluorinated β-carbon and is therefore less likely to undergo similar nucleophilic substitution that initiates 6:2 FTCA conjugation. Instead, 5:3 FTCA could be reversibly transformed to its α,β-unsaturated intermediate, 5:3 FTUCA, via microbial dehydrogenase [17]. As an α,β-unsaturated carboxylate, 5:3 FTUCA acts as a Michael acceptor that could conjugate with MSH or other cysteine thiolates [47]. Resonance with the carbonyl pulls electron density from the β-carbon, making it electrophilic. The thiolate nucleophile of MSH likely attacks this center, while a proton adds to the α-carbon. Therefore, MSH can perform 1,4-Michael addition [45, 48] to 5:3 FTUCA, leading to the formation of the saturated MS-5:3 FTCA conjugate (Figure 4). As this step does not involve C-F bond cleavage, fluoride is not released. Further Mca-mediated hydrolysis cleaves the amide linkage, releasing its corresponding mercapturic acid, AcCyS-5:3 FTCA.

### 2.3 Downstream Thiol-containing Metabolites Derived from Mercapturic Acid Conjugates

Iterative enrichments of TPs during FTCA biotransformation by RHA1 revealed three additional families of thiol-containing metabolites, including cysteine-(CyS-), cysteamine-(CAS-), and thiolate-(HS-) conjugates, as downstream derivatives of primary mercapturic acid conjugates AcCyS-5:3 FTUCA and AcCyS-5:3 FTCA (Table S5). The formation of CyS-conjugates (e.g., CyS-5:3 FTUCA and CyS-5:3 FTCA) can be initiated by the deacetylation that hydrolyzes the amide bond in mercapturic acid conjugates and removes the *N*-acetyl group, catalyzed by deacetylase [49] or aminoacylase [50]. Subsequently, decarboxylation of the cysteine α-carboxyl group catalyzed by decarboxylases may occur and generate aminothiol cysteamine (CAS) derivatives (e.g., CAS-5:3 FTUCA and CAS-5:3 FTCA) [51–53]. Potential further cleavage of the carbon-amino tail in CAS-conjugates could release thiolate-conjugates (e.g., S-5:3 FTUCA) [54]. Alternatively, CyS-conjugates may be transformed to thiolate-conjugates through the enzymatic action of cystathionine β-lyase [55–57] and release pyruvate and ammonium.

Although the exact order of steps and enzyme specificities of these biotransformation processes remain to be confirmed, we propose a putative pathway for these downstream thiol-containing metabolites over 6:2 and 5:3 FTCA biotransformation (Figure 4). Their accumulation across multiple enrichment cycles (Figure S4a-c, j, k) coupled with their CID MS2 fragmentation profiles shown in Figure S19-S25. Importantly, these observed downstream thiol-containing metabolites are unlikely to be artifacts of in-source fragmentation from the parent MS- and mercapturic acid conjugates. This interpretation is supported by three lines of evidence: (1) tandem mass spectrometry of the parent conjugates did not produce fragment ions corresponding to either CyS- or CAS-conjugates (Figure 3a-c; Figure S5-S7), (2) the parent-fragment ion intensity ratios varied across the enrichment process rather than remaining constant as expected for an in-source fragmentation yield, and (3) these metabolites exhibited distinct temporal dynamics in their biotransformation profiles (Figure 5; Figure S10 and S35) from their parent conjugates, as discussed in Section 2.5.

**Figure 5.**
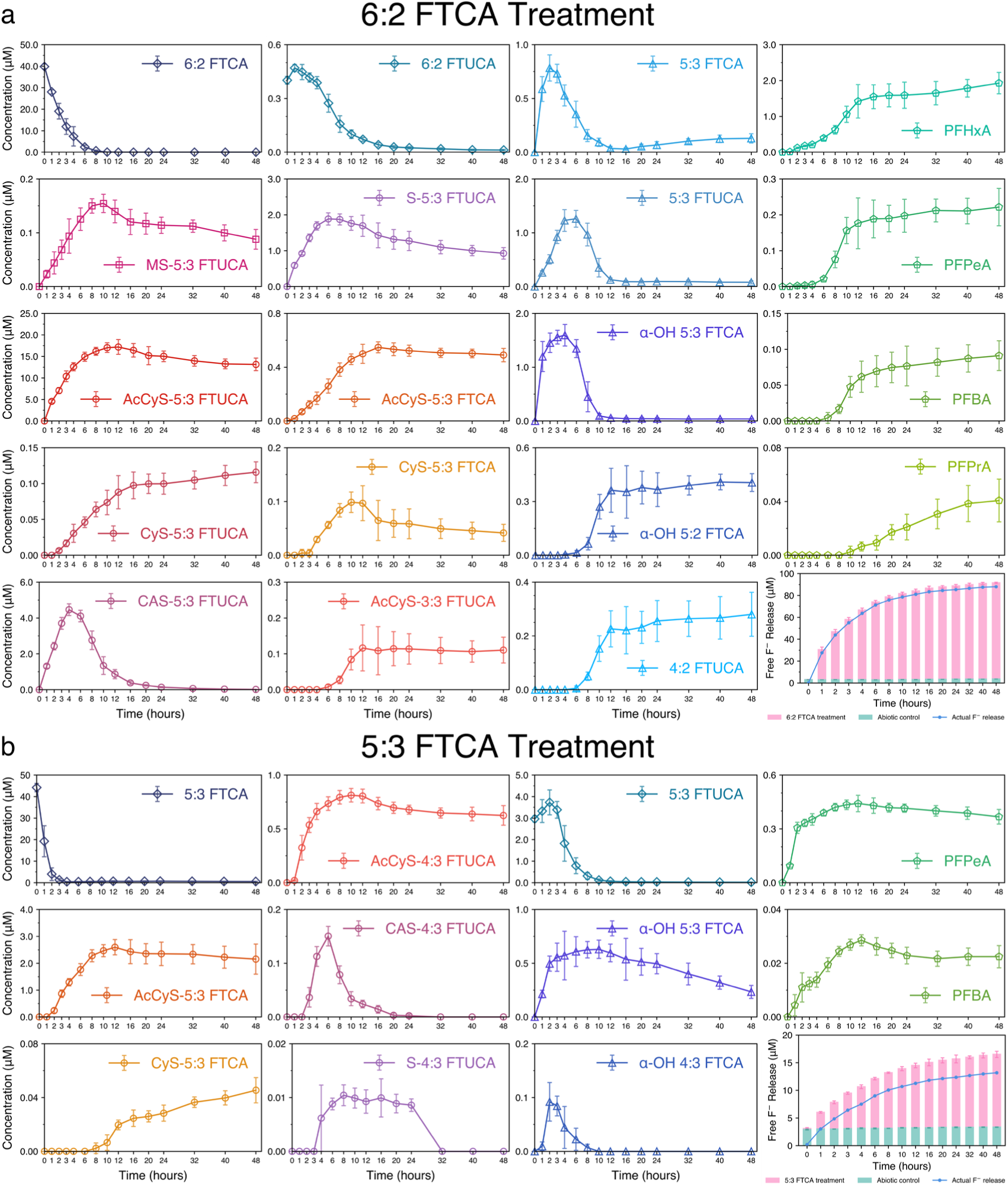
Target and non-target TP profiles during 48-h 6:2 FTCA (a) and 5:3 FTCA (a) (b) biotransformation by *Rhodococcus jostii* RHA1, respectively, along with FTCA depletion and F^-^ release. Error bars represent the standard deviation of three replicates.

### 2.4 Interlinked MSH-Mediated Conjugation and One-Carbon Removal Pathways during Cascading FTCA Biotransformation

Enabled by Kendrick mass defect (KMD) mapping (Figure 3g), homologous series of mercapturic acid conjugates and their downstream thiol-containing derivatives were identified. Each series differs systematically by a -CF2-unit, forming clear homologous series bearing progressively shorter lengths of fluorinated alkyl tails (Table S5). Notably, the detection of these conjugates coincided with the appearance of characteristic chain-shortened TPs, such as n:2 FTCA/FTUCAs, n:3 FTCA/FTUCAs, and hydroxy-n:3 FTCAs. Sequential biotransformation from 6:2 FTCA to 5:3 FTCA/5:3 FTUCA, then to 5:2 FTCA, 4:3 FTCA, and later 4:2 FTUCA was observed, accompanied by remarkable accumulation of C3-C6 PFCAs (Figure 2g; Figure S4d-f). A striking pattern is noted that the intensities of these intermediates declined substantially as the chain length decreased, supporting the similar characteristic progression of “one-carbon removal pathways” [58]. This observation is in good agreement with prior reports of sequential fluorinated chain shortening during FTCA biotransformation [18, 19, 23, 58].

Crucially, the newly discovered MSH-mediated conjugation reactions appear to be interlinked with “one-carbon removal pathways”, together forming a cascading network as depicted in Figure 4. Regardless of its chain length, each partially degraded FTCA intermediate can serve as a viable substrate for further MSH-mediated conjugation (e.g., AcCyS-5/4/3:3 FTUCA as shown in Figure 2b, e, f; Figure S5, S19, and S20; Table S5). Thus, the biotransformation proceeds through a dynamic interplay: chain-shortening reactions can generate fluorotelomer intermediates, which in turn become available for subsequent MSH conjugation, giving rise to a spectrum of structurally related conjugates. This interconnection suggests that rather than operating as isolated processes, “one-carbon removal pathways” and MSH-mediated conjugation pathways are mechanistically coupled, influencing each other across sequential metabolic steps. While the exact enzymatic mechanisms and regulatory controls remain to be fully elucidated, our results support a model of cascading, reiterative biotransformation in which MSH-mediated conjugation occurs in parallel with stepwise chain-shortening.

### 2.5 Dominant Generation of Mercapturic Acid Conjugates Coupled with Fluoride Release

Following the observation of conjugated and other TPs in the iterative enrichment assays, we employed a semi-quantification approach to profile the TP dynamics over a 48-h exposure of *R. jostii* RHA1 resting cells to individual FTCAs (Figure 5; Table S6 and S7). A calibrated series of the glutathione–monobromobimane (GS-mBBr) conjugate was prepared as a reference standard for the semi-quantification of thiol-conjugated TPs (SI Section SI1.6; Figure S36 and S37), and analyte concentrations were determined using calibration curves based on relative intensity (RI) (SI section S1.12; Figure S38; Table S8).

Exponential removal of 6:2 FTCA from an initial 39.83 ± 0.98 µM to < MDL occurred within the first 12 h of incubation (Figure 5a; Table S6), coupled with a rapid fluoride release to 81.04 ± 1.33 µM (≈ 2.03 mol F^-^ release per mole of 6:2 FTCA removed). Gradual liberation of fluoride continued at a slower pace thereafter, rising to 87.96 ± 0.57 µM at 48 h. The direct MSH conjugate, MS-5:3 FTUCA, culminated to 10 h, consistent with the depletion of the parent FTCA. Its hydrolysis product, AcCyS-5:3 FTUCA, accumulated to 17.19 ± 1.41 µM later by 12 h. The temporal offset between the MS- and AcCyS-conjugates supports the biotransformation sequence proposed in Section 2.2. The concentration of AcCyS-5:3 FTUCA decreased slightly afterwards and remained stable at 13.14 ± 1.20 µM by the end of the incubation.

As shown in the time-series data, the chain-shortened intermediate 5:3 FTCA was also formed during 6:2 FTCA biodegradation and reached its peak at 0.78 ± 0.10 µM as early as 2 h. Its dehydrogenated product, 5:3 FTUCA, accumulated and built up later to 1.26 ± 0.13 µM at 6 h. Both of these TPs declined rapidly thereafter, while the corresponding mercapturic acid AcCyS-5:3 FTCA rose to 0.55 ± 0.03 µM by 16 h (Figure 5a). Similarly, when RHA1 was exposed directly to 5:3 FTCA, rapid removal of 5:3 FTCA occurred within 4 h from 44.19 ± 1.58 µM to below 1 µM, exhibiting a faster removal rate than that of 6:2 FTCA (Figure 5b). AcCyS-5:3 FTCA was the major conjugated TP, culminating steadily to a maximum at 12 h and remaining dominant till the end of the incubation.

Coupling with the formation of these C8 mercapturic acid conjugates, their downstream thiol-containing metabolites (e.g., CyS-5:3 FTCA) followed a similar accumulation trend but declined more steeply after their peaks (Figure 5). CyS-5:3 FTUCA increased consistently throughout the incubation, while cysteamine conjugates CAS-5:3 FTUCA and CAS-5:3 FTCA (Figure S10) were transient, suggesting potential instability or rapid metabolic conversion. Concurrently or shortly after, the accumulation of mercapturic acid conjugates with shorter fluorotelomer tails (e.g., AcCyS-4:3/3:3 FTUCA) were observed, along with the emergence of their downstream thiol-containing derivatives. This strongly implies a stepwise -CF2-removal sequence that forms 5:2 and 4:2 FTCAs from the initial C8 precursors. The synchronous detection of C8 and shorter-chain FTCA derivatives (e.g., α-OH 5:3/4:3 FTCA and 5:2/4:2 FTUCA, Figure S30, S31, and S33), and C3-C6 PFCAs further support the occurrence of sequential chain-shortening process (Figure 5a). Taken together, the time-resolved data corroborate a cascading biotransformation network wherein MSH-mediated conjugation process and “one-carbon removal pathways” are functionally interlinked rather than parallel tracks. The consistent appearance and temporal order of these metabolites reinforce the hypothesis that chain shortened FTCA intermediates can serve as substrates for further MSH conjugation, thereby sustaining an iterative and overlapping metabolic sequence (Figure 4).

The 48-hour time-series TP profiling enabled a quantitative evaluation of the fluorine-based mass balance and the relative contributions of different metabolic pathways during FTCA biotransformation and biodefluorination (Table 1; Table S6 and S7). Total thiol conjugates derived from MSH-mediated conjugation accounted for a dominant portion of the overall transformation, representing 37.49 ∼ 51.97 % of the removed 6:2 FTCA and 6.47 ∼ 7.92 % of the removed 5:3 FTCA by RHA1. The conjugate pool contributed 4.7 ∼ 8.4-fold (6:2 FTCA) and 3.2 ∼ 4.5-fold (5:3 FTCA) greater than the pool of PFCAs and other FTCA intermediates formed via “one-carbon removal pathways”. Incorporating newly identified conjugates significantly improved the fluorine balance, increasing the fluorine mass recovery (FMR, SI Section S1.7) to ∼64.57 % and ∼12.40 % for 6:2 and 5:3 FTCA at 12 h, respectively. These findings indicate the substantial role of MSH-mediated processes, which have previously been underappreciated in PFAS biotransformation studies.

**Table 1.** Quantitative contribution of MSH-mediated conjugation and “one-carbon removal pathways” and fluorine-based mass balance for FTCA biotransformation and biodefluorination by *R. jostii* RHA1.

| Time Point | 6:2 FTCA Treatment |  |  | 5:3 FTCA Treatment |  |  | 6:2 FTCA Killed Control |  | 5:3 FTCA Killed Control |  |
| --- | --- | --- | --- | --- | --- | --- | --- | --- | --- | --- |
|  | 12h | 24h | 48h | 12h | 24h | 48h | 24h | 48h | 24h | 48h |
| ΔFTCAs (μM) | 39.81 | 39.83 | 39.82 | 43.54 | 43.49 | 43.65 | 1.60 | 2.16 | 3.85 | 2.91 |
| ΔConjugation TPs (μM) | 20.70 | 17.35 | 14.93 | 3.46 | 3.07 | 2.82 | 0.00 | 0.00 | 0.00 | 0.00 |
| ΔConjugation TPs/FTCA Consumed (μM/μM) | 51.97% | 43.56% | 37.49% | 7.92% | 7.03% | 6.47% | - | - | - | - |
| ΔPFCA TPs (μmol/L) | 1.67 | 1.89 | 2.28 | 0.47 | 0.44 | 0.39 | 0.00 | 0.00 | 0.00 | 0.00 |
| ΔPFCA TPs/FTCA Consumed (μM/μM) | 4.18% | 4.74% | 5.73% | 1.08% | 1.00% | 0.89% | - | - | - | - |
| $\Delta$"One-Carbon Removal" TPs (μM) | 0.81 | 0.82 | 0.93 | 0.60 | 0.50 | 0.23 | 0.00 | 0.00 | 0.00 | 0.00 |
| $\Delta$"One-Carbon Removal" Intermediate /FTCA Consumed (μM/μM) | 2.04% | 2.07% | 2.34% | 1.37% | 1.14% | 0.54% | - | - | - | - |
| F<sup>-</sup> Release (μM) | 81.05 | 85.34 | 87.96 | 11.27 | 12.36 | 13.17 | 0.49 | 0.61 | 0.40 | 0.43 |
| Avg. F<sup>-</sup> Release per FTCA Molecule Consumed | 2.04 | 2.14 | 2.21 | 0.26 | 0.28 | 0.30 | - | - | - | - |
| Theor. F<sup>-</sup> Release from Conjugation TPs /Total F<sup>-</sup> Release (μM/μM) | 47.59% | 39.96% | 34.45% | 12.73% | 10.46% | 9.49% | - | - | - | - |
| Theor. F<sup>-</sup> Release from PFCA TPs /Total F<sup>-</sup> Release (μM/μM) | 4.51% | 5.23% | 6.38% | 7.57% | 7.00% | 6.26% | - | - | - | - |
| Theor. F<sup>-</sup> Release from "One-Carbon Removal" Intermediate /Total F<sup>-</sup> Release (μM/μM) | 2.62% | 2.75% | 3.08% | 0.00% | 0.00% | 0.00% | - | - | - | - |
| Fluorine Mass Recovery (FMR, %) | 64.57% | 58.70% | 55.09% | 12.40% | 11.63% | 10.21% | 96.08% | 94.69% | 91.75% | 93.79% |

Notably, considerable biodefluorination was observed during 6:2 FTCA biotransformation, with an average of ∼2.21 fluoride ions released per molecule after 48 h (Figure 5a). Among two biotransformation pathways, metabolites formed through MSH-mediated conjugation accounted for 34.45 ∼ 47.59 % of this fluoride release, compared to only 7.13 ∼ 9.46 % can be explained by the “one-carbon removal pathways”. Although fluoride release during 5:3 FTCA biotransformation was limited (up to 13.17 ± 0.34 µM), the contribution from thiolate conjugates generated after chain-shortening to β-fluorinated n:2 homologs and subsequent MSH conjugation still exceeded that from PFCAs and other chain-shortening products combined (Table 1; Figure 5b). Taken together, these lines of evidence reveal that MSH-mediated conjugation operates with high flux and thereby supplies most of the observed defluorination, making it a quantitatively dominant route not only in FTCA biotransformation but also in promoting biodefluorination. Neglecting this pathway would therefore leave the fluoride profile unexplained (i.e., low FMR) and underestimate the degradative capacity of *Actinomycetota*.

### 2.6 Thiolate Conjugated Metabolite Formation in Other *Actinomycetota*

Batch assays with *R. jostii* RHA1 revealed MSH-mediated conjugation as a dominant pathway responsible for FTCA biotransformation and defluorination. Since this MSH conjugation process is well-recognized across the phylum *Actinomycetota* as a detoxification response against electrophilic xenobiotics, its interaction with FTCAs was validated using three additional MSH-producing, phylogenetically distinct *Actinomycetota* species, including *Mycobacterium smegmatis* MC^2^155, *Mycobacterium dioxanotrophicus* PH-06, and *Pseudonocardia dioxanivorans* CB1190.

As the 6:2 FTCA-derived TPs profiled (Figure 6), characteristic MSH-derived metabolites, including the primary conjugates and their downstream derivatives, were consistently observed in resting cell assays across all three additional strains, which accounted for 31.71 ∼ 55.41 mol% of the total identifiable TPs recovered from the biotransformation of 6:2 FTCA (Figure 6a). The formation of conjugates co-occurred with 6:2 FTCA removal, fluoride release, and the formation of PFCAs and other intermediates associated with “one-carbon removal pathways”, which parallel the patterns observed in the RHA1 assays. These findings therefore support the wide occurrence of MSH-mediated conjugation across diverse *Actinomycetota* species, albeit with strain-specific metabolic capabilities.

**Figure 6.**
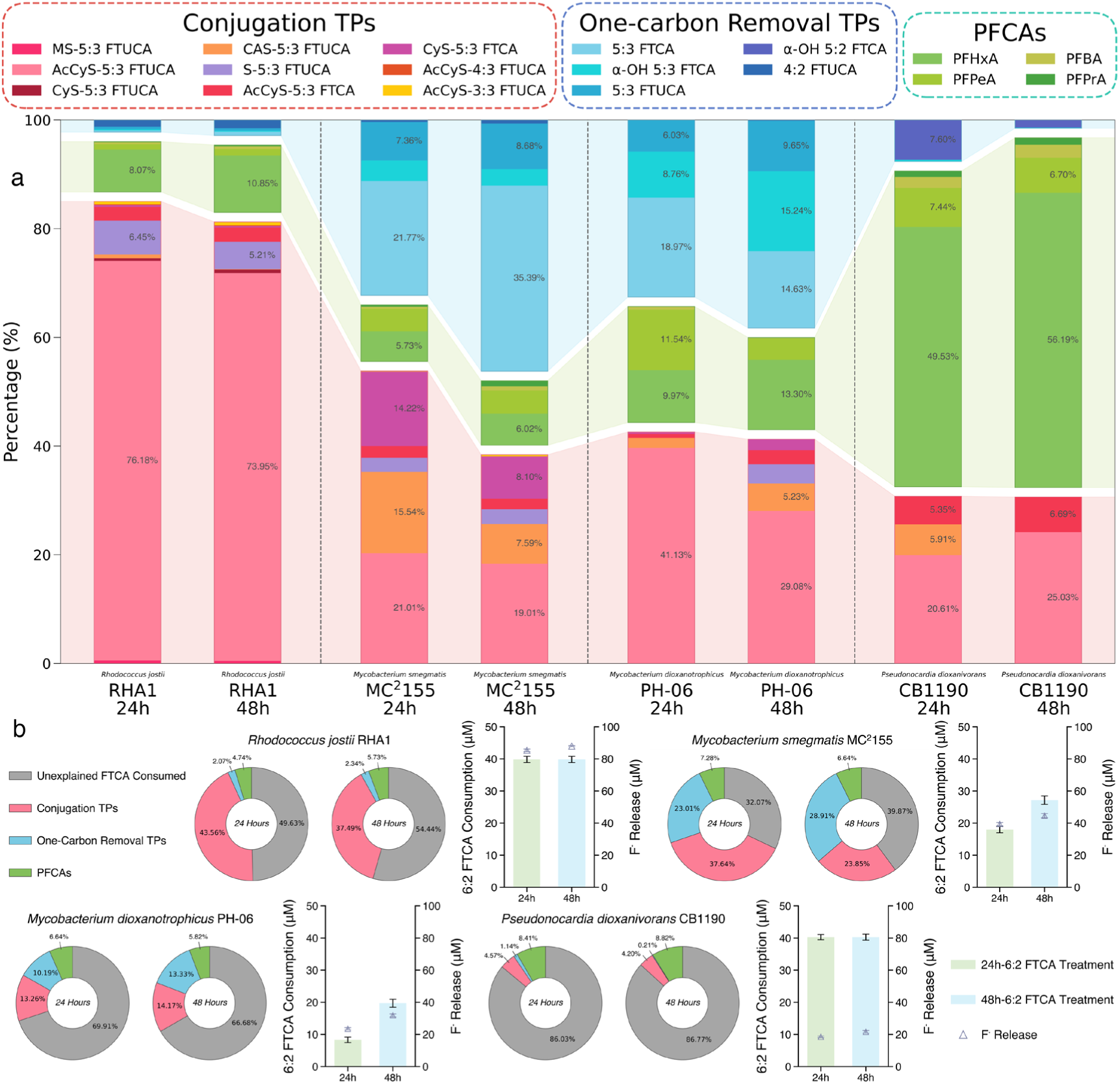
MSH-mediated conjugation with 6:2 FTCA across four *Actinomycetota* strains, including *Rhodococcus jostii* RHA1, *Mycobacterium smegmatis* MC^2^155, *Mycobacterium dioxanotrophicus* PH-06, and *Pseudonocardia dioxanivorans* CB1190. (a) TP profiles after 24-h and 48-h exposure to 6:2 FTCA. (b) 6:2 FTCA removal, F^-^ release, and the mass discrepancies unaccountable by MSH-mediated conjugation and “one-carbon removal pathways”-derived intermediates or PFCAs.

Among the strains tested, MC^2^155 and PH-06 showed slower and incomplete removal of 6:2 FTCA after 48 h (69.03 % and 47.49 %, respectively), and the TP contributions from MSH-derived conjugates were comparable to those generated from “one-carbon removal pathways” (Figure 6). The conjugation intermediate CAS-5:3 FTUCA, which was transient in RHA1, exhibited greater persistence in both *Mycobacterium* strains. Chain-shortened conjugates, including AcCyS-4:3 FTUCA and AcCyS-3:3 FTUCA, were observed in RHA1 and MC^2^155 during biotransformation of 6:2 FTCA (Figure 6a). This confirms that intermediates from sequential one-carbon shortening (i.e., 5:2 FTCA, 4:2 FTCA) can be conjugated with MSH to extend the cascade. In contrast, complete removal of 6:2 FTCA was achieved by CB1190 within 24 h. However, the combined concentration of quantifiable conjugated and chain-shortened TPs accounted for < 14 mol% of the consumed 6:2 FTCA, suggesting the potential involvement of currently uncharacterized TPs and alternative metabolic routes.

Strains MC^2^155 and PH-06 showed moderate 5:3 FTCA removal, generating TP profiles involving both MSH-mediated conjugation and the “one-carbon removal pathways” (Figure S11). Specifically, ∼18% and ∼56% of the initially dosed 5:3 FTCA was biotransformed by MC^2^155 and PH-06, respectively, while minimal by CB1190. AcCyS-5:3 FTCA and its downstream cysteine derivative, CyS-5:3 FTCA were consistently detected in both strains. Furthermore, conjugated derivatives of 5:2 FTCA, including AcCyS-4:3 FTUCA and its downstream metabolites, CAS-4:3 FTUCA and S-4:3 FTUCA, were also identified, supporting that 5:3 FTCA first underwent “one-carbon removal pathways” to 5:2 FTCA and validating the functional interlink between these two metabolic routes (Figure S11a). Conjugated TPs constituted ∼40% and ∼22% of the total recovered TPs in MC^2^155 and PH-06 at 24 h, respectively. Chain shortening TPs, including PFCAs (e.g., PFPeA, PFBA) and hydroxy-FTCAs (e.g., α-OH 5:3 FTCA, α-OH 4:3 FTCA), were also detected, albeit at lower concentrations. Fluoride release remained limited at ∼7 µM (Figure S11b), which is in good agreement with the dominance of the non-defluorinating conjugation.

Collectively, the findings confirm that MSH-mediated conjugation is widely conserved among *Actinomycetota* when exposed to FTCAs, though its extent varies by strain. The observed variation is possibly due to a combination of factors, including the expression, catalytic efficiency, and substrate specificity of MST and Mca, cell-surface properties and uptake kinetics, intracellular MSH availability, and strain-specific mechanisms controlling the interlink between the conjugation and “one-carbon removal pathways”. Nevertheless, a clear pattern was observed that mercapturic acid derivatives dominated the 6:2 FTCA TP profiles of RHA1, MC^2^155, and PH-06, exceeding the TPs formed through “one-carbon removal pathways”, while strain CB1190 produced a lower proportion of conjugation TPs (∼30%), and C3-C6 PFCAs dominate the pool of recovered TPs (∼60%). Overall, the capacities for FTCA biotransformation followed the order: RHA1 >> MC^2^155 > PH-06 > CB1190.

### 2.7 Structural and Evolutionary Characteristics of MST and Mca in RHA1 and Other

### Actinomycetota

As aforementioned, MST and Mca are two enzymes primarily involved in the initial MSH conjugation with FTCAs and the hydrolysis of these conjugates into mercapturic acids, respectively. The molecular basis for MSH biosynthesis and its mediated conjugation has been long centered on the model organism *Mycobacterium tuberculosis* H37Rv [59]. Here, sequences and structural similarities of these enzymes between *R. jostii* RHA1 and *M. tb* H37Rv were firstly compared. Sequence alignment revealed that the RHA1 MST and Mca share 52.5% and 69.6% amino acid identities with those in H37Rv homologs, respectively (Table S10 and S11). Three-dimensional enzyme structures predicted by AlphaFold2 (Figure S12) demonstrated structural conservation, with Cα root-mean-square deviations (RMSDs) as low as 0.516 Å for MST and 0.357 Å for Mca in RHA1 compared to the available crystal structures from H37Rv (Figure S13).

To assess if this similarity extends beyond RHA1 and H37Rv, we analyzed MST and Mca from six additional MSH-producing *Actinomycetota*, including *M. smegmatis* MC^2^155, *M. Dioxanotrophicus* PH-06, and *P. dioxanivorans* CB1190, whose capabilities to conjugate FTCAs with MSH were experimentally validated, as well as three *Actinomycetota* species previously reported to produce MSH from other genera [60] (i.e., *Kocuria rhizophila* DC2201, *Streptomyces griseus* subsp. *griseus* NBRC 13350, and *Kineococcus radiotolerans* SRS30216) as representatives. Eight *Actinomycetota* representatives (Table S9) shared amino acid identities of 45.12% ∼ 76.83% and 52.72% ∼ 78.75% for MST and Mca homologs with those in H37Rv, respectively (Table S10 and S11). Pair-wise RMSDs relative to the crystal structure of H37Rv were 0.255 Å ∼ 1.256 Å for MST and 0.212 Å ∼ 0.697 Å for Mca (Figure 13). Furthermore, sequence and structural alignments of the four MSH-biosynthetic enzymes MshA–D (Table S12-S15; Figure S14 and S15) also demonstrated consistently high similarity among the *Actinomycetota* representatives, indicating the conservation of the entire MSH-mediated processes.

Sequence logos (Figure 7a, d; Figure S42) generated from 150 MST and Mca homologs reveal that the residues responsible for substrate catalysis are highly conserved. The MSH binding and the subsequent catalysis of FTCA conjugation in RHA1 MST are mediated through a suite of conserved amnio acid residues including Asn39, His47, Tyr86, Gly87, Trp134, Asp150, His154, and Gln157 [59]. Asn39/Gly87 together form a clamp that locks the glucosamine (GlcN) ring and help orient the cysteinyl thiol. Asp150 forms a H-bond with the cysteine amide of MSH and organizes for S-transfer. Gln157 forms a H-bond with the inositol (Ins) ring (Figure 7b). A conserved His47–Asp150–His154 tripod structure immobilizes a divalent metal ion (typically Zn^2+^), activating the sulfur and adjusting MSH for nucleophilic attack on FT(U)CAs. Tyr86/Trp134 potentially undergo a coordinated structural shift upon MSH/metal binding, forming a hydrophobic gate that controls access of FTCAs to the active site (Table S16) [59].

**Figure 7.**
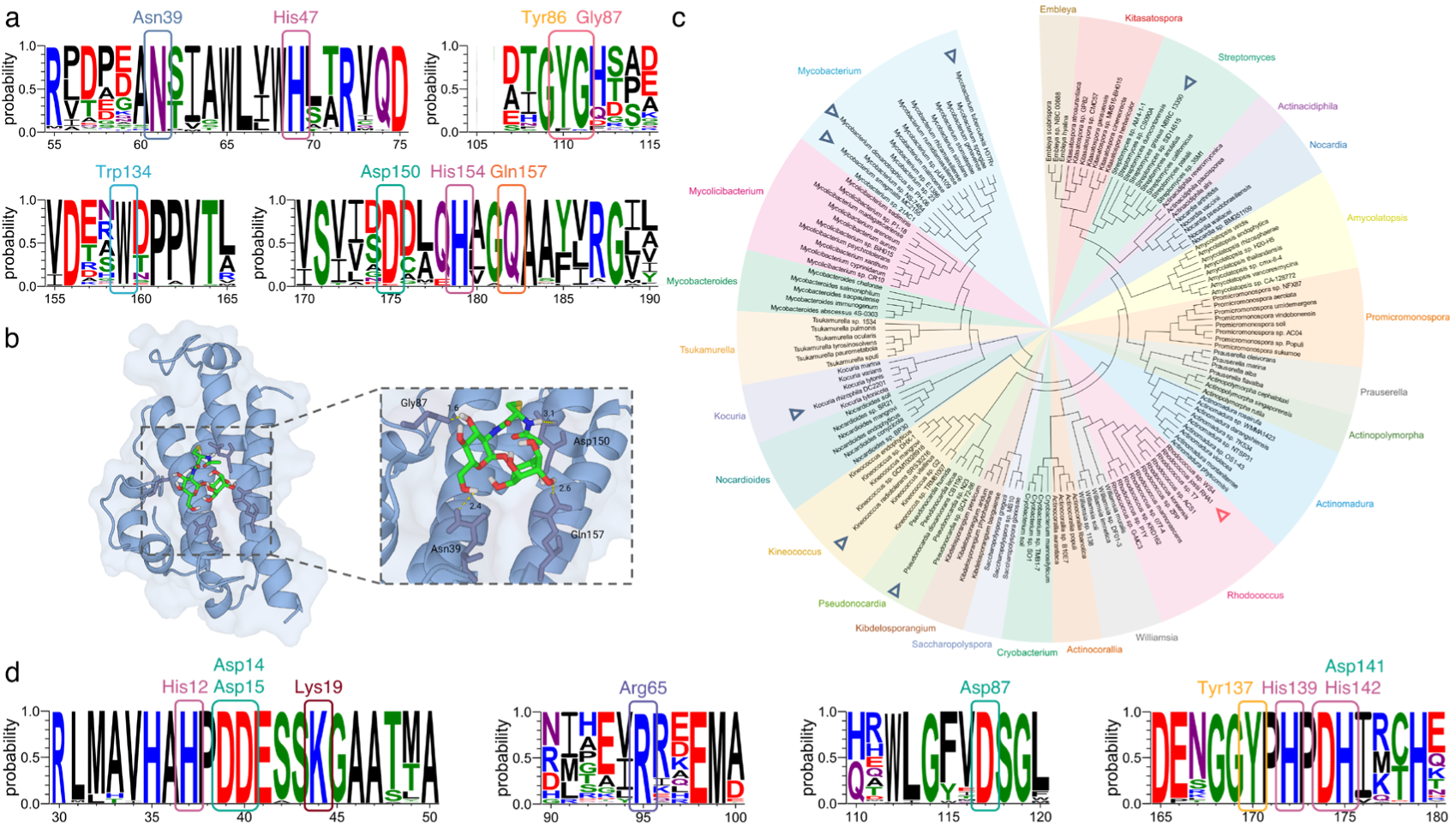
Highly conserved sites for MST (a) and Mca (d) that are responsible for MSH conjugation and subsequent hydrolysis of MS-conjugates across 150 *Actinomycetota* species, respectively. (b) AlphaFold2 structural model and key binding residues of *R. jostii* RHA1 MST with a predicted docked MSH molecule. (c) Phylogenetic tree of 150 MST homologs from 24 *Actinomycetota* genera. Triangles highlight eight *Actinomycetota* species for structural modeling and comparative analysis shown in Figure S13-S15.

Similar to MST, the His12–Asp15–His142 tripod structure in Mca positions the metal ion, polarizing the amide of the MS-R conjugates [42, 61, 62]. Asp14 initiates hydrolysis by deprotonating a metal-bound water molecule, generating a reactive hydroxide ion that attacks the amide carbonyl [42]. His139 acts as an electrophilic residue in the Mca catalytic center. Its Nε-imidazolium forms a H-bond with the carbonyl, stabilizes the oxyanion intermediate, and protonates the leaving GlcN-Ins amine to complete hydrolysis, while Asp141 forms a H-bond with His139 to stabilize its protonated state (Figure S16) [42, 61, 63, 64]. Lys19 is a unique conserved residue in Mca that distinguishes it from MshB, and its ε-ammonium side chain could prevent the entry of 1D-myo-inositol 2-acetamido-2-deoxy-alpha-D-glucopyranoside (GlcNAc-Ins) [64]. Other conserved residues, including Tyr137 [63], Arg65 [62, 64] and Asp87 [62], contribute to stabilizing the transition intermediate and positioning of the MS-R conjugates. Details of all conserved residues are presented in Table S17.

Catalytic amino-acid motifs are highly conserved across *Actinomycetota* for both enzymes (Table S18 and S19). In MST, the mentioned residues are strictly conserved among *Actinomycetota* representatives, except that Tyr86 is replaced by Phe in *K. rhizophila* DC2201 (Table S18; Figure S41). Analysis of a broader set of 150 MST homologs from 24 *Actinomycetota* genera further confirms this conservation (Figure 7a; Figure S42). All Mca’s catalytic residues are 100% conserved across eight representatives (Table S19; Figure S41).

The broader 150 Mca sequence set (Figure 7d; Figure S42) supports the observed strict conservation.

Maximum-likelihood phylogenetic trees built from 150 non-redundant MST (Figure 7c) and Mca (Figure S17) sequences resolve 24 *Actinomycetota* genera into monophyletic clades with internal branching. MST homologs are not restricted to *Rhodococcus* and *Mycobacterium* and found to be distributed across the *Actinomycetota* phylum, including *Kineococcus*, *Amycolatopsis*, *Streptomyces*, and *Nocardia*. All 150 genomes also encode the complete enzymatic machinery required for MSH-mediated processes, including MST, Mca, and MshA– D responsible for MSH biosynthesis. This reveals that MSH-mediated conjugation is a widespread and conserved strategy across the phylum, expanding the pool of microorganisms with PFAS biotransformation potentials. Although *Rhodococcus* and *Mycobacterium* species are evolutionarily divergent, they showed similar kinetic profiles for FTCA conjugation in the batch assays, revealing functional conservation across phylogenetic distance. Poorly characterized genera, such as *Nocardiodes* and *Amycolatopsis*, are MSH producers and thus represent promising but unexplored reservoirs of enzyme variants that are worth future PFAS biotransformation investigations.

### 2.8 Prevalence of Conjugated Metabolites in Sludge Microbiomes

In addition to pure culture experiments, we investigated the formation of thiol-conjugated metabolites in six different activated sludge microbiomes previously exposed to FTCAs. These samples originated from two studies: a short-term (7-day) incubation of activated sludge samples from four municipal WWTPs in the New York metropolitan area (i.e., P, L, R, and W; Table S20) with 6:2 or 5:3 FTCAs in synthetic wastewater (with glucose and acetate as carbon sources) [18], and a separate long-term enrichment of other sludge samples from two additional WWTPs (i.e., BH and V; Table S20) in FTCA-spiked ammonium mineral salts (AMS) medium [20] without extra carbon supplementation. Along with significant 6:2 FTCA biotransformation, consistent generation of AcCyS-5:3 FTUCA, CyS-5:3 FTUCA, and S-5:3 FTUCA were observed at high intensity levels in all six sludge microbiomes (Figure 8a). MS-5:3 FTUCA and AcCyS-3:3 FTUCA were both detected in three of the six sludges, particularly in samples where other conjugated metabolites co-occurred in high abundance. CAS-5:3 FTUCA was not positively detected in any of these sludge samples, aligning with its transient behavior observed in the RHA1 resting cell assays.

**Figure 8:**
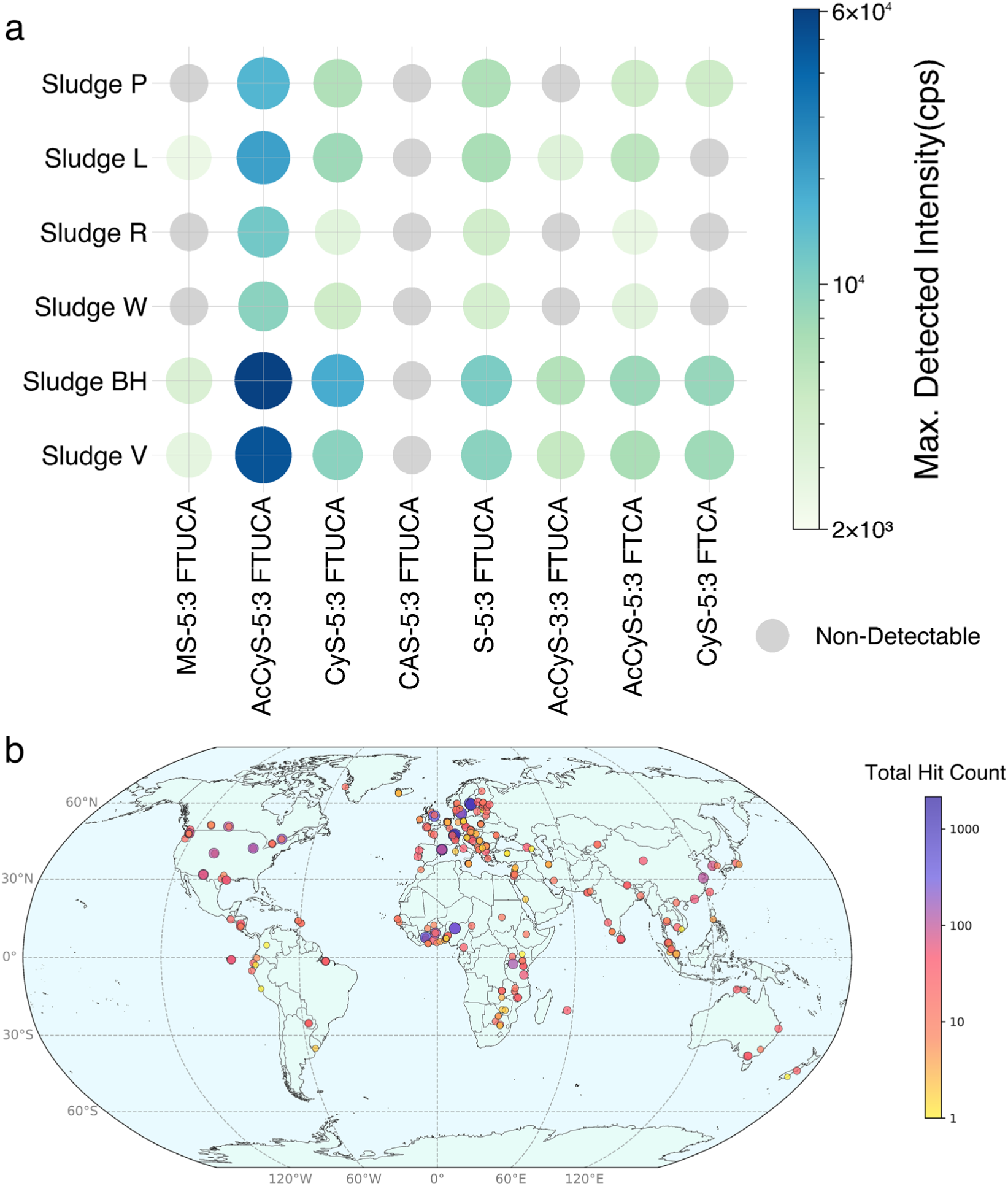
(a) Heatmap of the maximum detected intensities of key MSH-mediated conjugated TPs in six activated sludge microbiomes when incubated with FTCAs with (Sludge P, L, R, and W) or without (Sludge BH and V) the supplementation of external carbon sources. (b) Widespread occurrence of MST homologs across the global WWTP activated sludge metagenomes. Each circle represents the location where MST homologs were detected, and circle color and size correspond to the number of homolog positive hits identified.

Exposure to both 5:3 and 6:2 FTCA led to the generation of AcCyS-5:3 FTCA in all sludge microbiomes. CyS-5:3 FTCA appeared in only three sludges (P, BH, and V). The intensities of AcCyS-5:3 FTCA from 5:3 FTCA-exposed sludges were approximately one order of magnitude lower than those of AcCyS-5:3 FTUCA from 6:2 FTCA exposures. Collectively, these observations demonstrate MSH-mediated conjugation coupled with mercapturic acid and cysteine conjugate formation can be widespread in response to FTCA biotransformation in diverse activated sludge microbial communities.

Although RHA1 and the other three examined *Actinomycetota* strains were not identified in the six sludge microbial communities, 16S rRNA profiling showed these sludge microbiomes are rich in *Propionicimonas* and other members of *Actinomycetota* [18]. These taxa could actively contribute the enzymes involved in the MSH-mediated conjugation processes.

To further evaluate the global prevalence of the MSH-mediated pathway, we screened for MST homologs from WWTP activated sludge and aquatic environment metagenomes available in the MGnify repository. Hidden-Markov models (HMMs) built from *R. jostii* RHA1 and *M. tb* H37Rv MST homolog sequences recovered 14,141 and 88,213 non-redundant hits (E-value < 1e-5) across WWTP activated sludge and aquatic metagenomes worldwide, respectively (Figure 8b; Figure S18). Matches were observed on every inhabited continent, with dense clustering in Europe and Africa, probably due to current bias in publicly available metagenomes. The global presence of MST homologs indicates that diverse microbial communities possess the genetic potential for MSH-mediated conjugation. Overall, the direct detection of MSH conjugate derivatives in six FTCA-exposed activated sludges, along with the worldwide prevalence of MST homologs, supports MSH-mediated conjugation as a significant, yet previously under-recognized mechanism for PFAS biotransformation in both engineered and natural ecosystems.

## 3. Materials and Methods

### 3.1 Chemicals and Reagents

6:2 FTCA (3,3,4,4,5,5,6,6,7,7,8,8,8-tridecafluorooctanoic acid, ≥ 98%) and 5:3 FTCA (2H,2H,3H,3H-perfluorooctanoic acid, ≥ 97%) (Table S3) were purchased from Synquest Laboratories (Alachua, FL, USA). Stock solutions of both FTCAs (40 mM) were prepared in MeOH/Millipore water (2:1, v/v) mixture, passed through 0.22 µm polyethersulfone (PES) filters, and stored at -20 °C. Isotope-labeled internal standards, such as M8-PFOA and M2-6:2 FTCA, as well as FTCA and PFCA calibration standards, were purchased from Wellington Laboratories (Guelph, ON, Canada), each with a purity of ≥ 98%. LC/MS-grade ammonium acetate (≥ 99%) and molecular biology grade D-(+)-glucose (≥ 99.5%) were purchased from Sigma-Aldrich (St. Louis, MO, USA). Optima^®^ LC/MS-grade acetonitrile and methanol (≥ 99.9%) were purchased from Thermo Fisher Scientific (Waltham, MA, USA). For the conjugated metabolite semi-quantification, L-glutathione (GSH, ≥ 98%), HEPES buffer (pH 8.0), N-ethylmaleimide (≥ 99%), and methanesulfonic acid (≥ 98%) were all obtained from Thermo Fisher Scientific. Bromobimane (mBBr, ≥ 99%) was purchased from Arctom Scientific (Westlake Village, CA, USA). Sodium fluoride powder (≥ 99%) was obtained from Thermo Scientific Chemicals. Total ionic strength adjustment buffer (TISAB II) was purchased from Millipore Sigma (Burlington, MA, USA). Ultrapure water (18.2 MΩ·cm) was produced on a Milli-Q RC Synthesis water purification system (Millipore Sigma). All chemicals and reagents were stored and managed in accordance with manufacturers’ recommendation.

### 3.2 Cultivation and Cell Preparation of *Rhodococcus jostii* RHA1

*Rhodococcus jostii* RHA1 was obtained from the Leibniz Institute (DSM44719, DSMZ, Braunschweig, Germany) and cultured according to supplier guidelines. Laboratory cell stocks were prepared by cultivating cells in liquid Luria-Bertani (LB) medium (30 °C, 130 rpm), and the mid-exponential phase culture was cryopreserved in 30% (v/v) glycerol at -80°C.

For resting cell preparations, single colonies were picked from LB agar plates and inoculated into 100 mL AMS medium [65], supplemented with 20 mM glucose as the sole carbon and energy source. Cultures were cultivated (30 °C, 130 rpm) to a mid-exponential growth phase (OD600 ≈ 1.0, ∼40 h). Cells were harvested by centrifugation (12,000 rpm, 15 min, 4°C), washed three times with 20× diluted AMS medium to remove residual carbon and salts, and sequentially resuspended in 5 mL of 20× diluted AMS medium to an OD600 ≈ 2.5. Cell assays were carried out in the diluted AMS medium to minimize interference from inorganic ions during TP screening by HRMS, while providing ionic strength to prevent osmotic stress and cell bursting.

### 3.3 Iterative Accumulation of FTCA Biotransformation Metabolites by RHA1

An iterative enrichment strategy was employed to accumulate TPs (Figure S2). Biotransformation assays were conducted using 5 mL *R. jostii* RHA1 resting cell suspension in 60 mL sterile amber serum bottles. Cells were diluted with 0.75 mL of AMS medium and 14.25 mL of Millipore water to achieve a final volume of 20 mL. FTCA stock solution (20 µL) was added to an initial concentration of 40 µM at the beginning of each cycle. At the end of each exposure cycle, cells were removed by centrifugation (12,000 rpm, 15 min). The supernatant containing the accumulated TPs was transferred to a new sterile serum bottle. Fresh RHA1 resting cell suspensions and diluted AMS medium were added to replenish the volume to 20 mL. A new dose of FTCA stock was introduced to initiate the subsequent exposure cycle. Iterative exposure cycles were conducted in an identical fashion. A set of controls and blanks including abiotic controls using autoclaved cells, microbial blanks with live cells without FTCAs, analytical controls with FTCA in media without cells, and laboratory reagent blanks, were prepared (Table S4; see details in SI section S1.1, S1.2) and processed in parallel. All treatments and controls/blanks were prepared in triplicate and incubated at 30 °C with shaking at 130 rpm for 48 h. Aliquot samples (0.2 mL) were collected at the start and end of each cycle for TP accumulation profiling by Nano-ESI-HRMS. Additionally, 2.0 mL samples were collected at the end of each cycle for fluoride analysis.

### 3.4 Time-series FTCA Biotransformation Assays by *R. jostii* RHA1 and Other

### Actinomycetota Model Species

Time-resolved biotransformation assays (0–48 h) were conducted for RHA1 following the same procedure described in cyclic enrichment assays. Briefly, prepared cell suspensions (20 mL) were spiked with 40 µM FTCA stock, with full set of triplicated controls and blanks prepared (Table S4). Each treatment or control assay was sampled at select intervals over 48 h (0, 1, 2, 3, 4, 6, 8, 10, 12, 16, 20, 24, 32, 40 and 48 h; 0.2 mL each) for TP profiling.

In addition to RHA1, we further investigated three MSH-producing *Actinomycetota* strains, including *Mycolicibacterium smegmatis* MC^2^155, *Mycobacterium dioxanotrophicus* PH-06, and *Pseudonocardia dioxanivorans* CB1190. Genomes of these three strains contain the complete MSH biosynthetic genes *mshA–D* and the *mst/mca* cassette for MSH-mediated conjugation (Table S10-S15). Notably, MC^2^155 is a model strain known for its high endogenous MSH production [66]. PH-06 and CB1190 available in our lab served as additional *Actinomycetota* representatives. Biotransformation assays for these three cultures were conducted under identical conditions to those for RHA1, except only sampled at 4, 16, 24, and 48 h. Detailed incubation method for these strains is provided in SI Section S1.10.

### 3.5 Free Fluoride Measurement

Aqueous samples were withdrawn, filtered through 0.22 µm PES filters, and stored at 4 °C for ≤ 24 h before analysis. 1.5 mL of the filtrate was mixed 1:1 (v/v) with TISAB II buffer. Free fluoride was measured at room temperature with an Orion™ Dual Star™ meter equipped with a fluoride-specific electrode (model 9609BNWP). Fresh sodium fluoride standards were prepared in an identical manner to establish a calibration curve (Figure S44) before each measurement batch.

### 3.6 PFAS Analysis by Nano-ESI-HRMS

Non-target screening of FTCA TPs was conducted following our established protocols [18, 34] on a high-resolution Q-Exactive™ hybrid quadrupole−Orbitrap™ mass spectrometer (Thermo Fisher Scientific) coupling with a nano-electrospray ionization (Nano-ESI) emitter (Figure S3). Analysis was conducted in negative ionization mode at a resolving power of 140,000 (at *m/z* 200) for MS1 acquisition. Structural identification of putative TPs used data-dependent acquisition (DDA) at a resolution of 80,000 (at *m/z* 200) for MS2 profiling.

The Nano-ESI emitter tips were prepared using a P-2000 laser-based micropipette puller (Sutter Instrument). Samples were introduced via direct infusion at a constant flow rate of 1.0 µL/min by a syringe pump (NE-1000, New Era Pump Systems, Inc.). A –3.0 kV potential was applied to the emitter to maintain consistent ionization. The capillary temperature was set to 300 °C. For each analytical run, acquisition was initiated only after the automatic gain control (AGC) stabilized at a 100% target (intensity setting of 1e6). The total ion current (TIC) variation was monitored to remain below 8% throughout each run.

Full scan MS1 spectra were acquired over a range of *m/z* 75-1,100. Each MS1 acquisition incorporated 75 scans, with each scan aggregating from 5 microscans, culminating in a total duration of ∼3.5 minutes per analytical run. Candidate MIIs were subjected to collision-induced dissociation (CID) for fragmentation analysis under DDA mode. The quadrupole filter was configured to an isolation window of ±0.2 Da at *m/z* 200, achieving a 500 (=m/Δm) full width at half maximum (FWHM) resolution. Putative structural identification was confirmed by the occurrence patterns of precursor and fragment ions in the MS2 spectra and their relative abundance shifts induced by varying collision energies (CE) applied (15-50 V).

Following data acquisition, raw MS1 and MS2 spectra were exported from Xcalibur^TM^ software and processed using a computational workflow (Figure 1b) including HRMS raw data preprocessing for noise reduction (SI Section S1.3), iterative statistical analysis for identification of accumulating TPs (SI Section S1.4), and CID fragmentation analysis for MII structure elucidation. The initial preprocessing of acquired MS1 spectra were used to identify reliable features prior to non-target screening. A multi-step filtering function integrating indicator-based statistics was employed, including signal-to-noise ratio (SNR) filtering (𝐸𝑞. 𝑠3), scan reproducibility filtering (𝐸𝑞. 𝑠4 − 5), background correction against controls and blanks, and RI filtering (𝐸𝑞. 𝑠10 − 12). Mass ions meeting all criteria (𝐸𝑞. 𝑠14) were retained for further analysis.

### 3.7 Non-target Analysis and Structural Identification of Enriched TPs

A customized non-target screening workflow was used for TP identification as featured by continuous accumulation during the iterative exposure experiments. Briefly, this algorithm quantitatively evaluates TP accumulation dynamics using two statistical indices, including the Cumulative Significance Index (CSI, 𝐸𝑞. 𝑠16 ) that evaluates significant changes in TP abundance across iterative cycles, and the Linearity Trend Index (LTI, 𝐸𝑞. 𝑠17) that assesses the consistency of TP accumulation trend. These indices are integrated into an Accumulation Score 𝑆 (𝑛) (𝐸𝑞. 𝑠18) to identify MIIs that exhibited significant and iterative enrichment. Features exceeding the threshold (𝐸𝑞. 𝑠20) were selected as MIIs for further analysis. The complete framework, including the specific equations, weighing factors, and score thresholds, is provide in the SI Section S1.4.

Molecular structures of MIIs were elucidated using their MS2 spectra. Fragments were interpreted using three approaches, including matching to a local fragment library developed for PFAS, manual interpretation based on exact mass, and predictions from Compound Discover 3.4, as showed in Figure 1b. The integrated evidence from MS1 and MS2 enabled the identification of FTCA-derived TPs and associated biotransformation pathways. A custom Python script was developed for mentioned analyses, available at https://github.com/Daitoueqaq/HRMS-modification. This script assists the workflow, from initial HRMS data preprocessing, to the statistical identification of putative accumulating TPs. Additionally, this script incorporates modules for suspect screening (SI Section S1.5) and the construction of Kendrick mass difference networks (SI Section S1.9). KMD maps were generated from processed MS1 spectra, with homolog series (ΔKMD ≤ 1 mDa; ≥ 2 members) were collected as MIIs [67].

### 3.8 Phylogenetic Analysis of MST and Mca in *Actinomycetota*

Evolutionary relationships of MST and Mca within *Actinomycetota* were constructed based on the alignment of full-length amnio acid sequences. MST and Mca sequences of *M. tb* H37Rv (Rv0043; Rv1082) and *R. jostii* RHA1 (RHA1_RS14880; RHA1_RS28575) were used as BLAST-P references against the NCBI database (Table S10-S11). Homolog sequences sharing ≥ 55% amino acid identity were obtained, as this threshold generally corresponds to < 1.0 Å Cα RMSD within the catalytic core regions [68]. We used CD-HIT (v4.8.1) to cluster sequences and reduce sequence redundancy at a 90% identity threshold. A collection of 150 representative *Actinomycetota* taxa were selected to comprise 24 genera, consisting of paired MST and Mca sequences. Genomes were then screened with HMMER v3.3.2 to confirm the presence of the MSH biosynthetic cassette *mshA-D*.

Multiple sequence alignments (MSAs) were generated in MEGA 11 using the ClustalW [69] and poorly aligned termini (< 10 aa) were trimmed. Maximum-likelihood trees were inferred based on the JTT substitution matrix [70], with initial branch arrangements by neighbor-joining. The phylogenetic trees with the best log-likelihood score (log L = -1.81 × 10⁴ for MST, Figure 7c; log L = -1.94 × 10⁴ for Mca, Figure S17) were retained.

Sequence logos were produced using WebLogo3 (v3.7.9) for MST and Mca sequences [71]. Logos were generated from sequence sets including (i) the 150 MST and Mca homologs, and (ii) sequences from the eight *Actinomycetota* representatives. The resulting logos were used to identify the conservation of catalytic residues and to support functional interpretations.

### 3.9 Protein Structure Prediction and Molecular Interaction Analysis

Protein sequences for MST, Mca, and the MSH-biosynthetic enzymes MshA-D of the eight *Actinomycetota* representatives were retrieved from the NCBI GenBank database (Table

S10-S15). Full-length sequences were modelled using ColabFold-AlphaFold2 (v2.3.2) for 3-D structure prediction, with parameters listed in the SI Section S1.8. The highest-ranked prediction for each protein was retained (pTM > 0.82 and mean pLDDT > 88). Low-confidence terminal segments were removed (e.g., a contiguous stretch of ≥ 3 residues at either terminus with pLDDT < 50). PyMOL (Schrödinger, LLC) was used to assess structural similarity through pairwise alignments among the predicted models and previously published structures (Figure S13-S15).

AutoDock 4.2.6 (The Scripps Research Institute, San Diego, CA, USA) was used to conduct molecular docking simulations with the binding of MSH to MST. Coordinates for RHA1 MST were taken from AlphaFold2 predictions, and the MSH ligand structure was converted to PDBQT file with Open Babel (v3.1). The docking grid box was centered on the metal ion-coordinating tripod, the putative active site of MST. The docking was run using the Lamarckian genetic algorithm (LGA), with detailed docking parameters provided in the SI (Table S21). Multiple independent docking runs were carried out and the lowest-energy and most populated cluster was considered as the preferred binding mode. The predicted binding free energies (ΔG), hydrogen-bond networks, metal contacts, and hydrophobic interactions were extracted from the conformation.

### 3.10 Analysis of Conjugated TPs in WWTP Samples

Filtered aqueous phase samples from our previous FTCA biotransformation studies [18, 20] were re-analyzed to examine the generation of MSH-conjugated TPs. All archived samples were stored at -20 °C. In the first study, sludge samples from WWTP P, L, R and W (1 g wet weight, Table S20) were incubated aerobically for 7 d in 50 mL synthetic wastewater (COD = 415 ± 21 mg/L, NH4^+^*-N* = 7.5 ± 0.4 mg/L, and pH = 7.3 ± 0.1) with glucose and acetate as external carbon sources. In the other study, same weight of sludge samples from WWTP BH and V (Table S20) was seeded to enrich consortia through four cycles of successive (bi)monthly enrichment in AMS medium without external carbon source. In both studies, FTCAs were dosed to achieve an initial concentration of ∼80 μM. For Nano-ESI-HRMS analysis, 75 µL of each stored sample was diluted with methanol to 1 mL. Screening of MSH-derived conjugates including MS-, AcCyS-, CyS-, CAS- and S-conjugates (Table S5) were carried out using the aforementioned method by Nano-ESI-HRMS.

### 3.11 Screening of MST Homologs in Global Activated Sludge and Aquatic Metagenomics

The MGnify portal (https://www.ebi.ac.uk/metagenomics/search/analyses, accessed January 2025) was accessed for available metagenomes from WWTPs and aquatic environments (e.g., groundwater, freshwater and marine). Protein coding sequences (CDS) from 1,022 WWTP and 6,621 aquatic metagenomic assemblies were downloaded. Relevant metadata, including study IDs, sample IDs, analysis IDs, geographic coordinates, biome descriptions and other features were collected using a custom Python script via the MGnify API (https://www.ebi.ac.uk/metagenomics/api/v1/).

Protein BLAST was run against the NCBI nr protein database using MST sequences from *R. jostii* RHA1 (Accession: WP_009475824.1) and *M. tb* H37Rv (Accession: CCP43174.1) as queries to build a reference sequence set of MST homologs. Hits with an amino acid identity larger than 55% were collected. CD-HIT was applied at a 90% identity clustering threshold to reduce redundancy. The resulting sequences were aligned using MAFFT (v7.310). An MST profile HMM was built using *hmmbuild* (HMMER v3.3.2) to screen the CDS from all downloaded metagenomes for MST homologs using *hmmsearch*. Positive hits with an E-value < 1e-5 were collected as output putative MST homologs. Geographic coordinates for each positive hit were assigned, with manual corrections applied to any missing or erroneous entries. The resulting counts of MST homologs detected in each sample were visualized by Python.

## 4 Acknowledgements

This work was funded by US Geological Survey (USGS, G24AP00026), National Science Foundation (NSF, CBET-1903597), and New Jersey Department of Environmental Protection (NJDEP, SR21-019). We thank Chen Wu and Jose Manual Diaz Antunes at New Jersey Institute of Technology for their assistance with the culture and sludge assays. Funders had no role in study design, data collection, and interpretation, or the decision to submit the work for publication.

We declare no competing financial interest.

## References

1. Evich, M.G., et al., Per- and polyfluoroalkyl substances eabg9065. in the environment. Science, 2022. 375(6580): p.

2. Rahman, M.F., S. Peldszus, and W.B. Anderson, Behaviour and fate of perfluoroalkyl and polyfluoroalkyl substances (PFASs) in drinking water treatment: a review. Water Res, 2014. 50: p. 318–40.

3. Xiao, F., Emerging poly- and perfluoroalkyl substances in the aquatic environment: A review of current literature. Water Research, 2017. 124: p. 482–495.

4. Ackerman Grunfeld, D., et al., Underestimated burden of per- and polyfluoroalkyl substances in global surface waters and groundwaters. Nature Geoscience, 2024. 17(4): p. 340–346.

5. Buck, R.C., et al., Perfluoroalkyl and polyfluoroalkyl substances in the environment: terminology, classification, and origins. Integr Environ Assess Manag, 2011. 7(4): p. 513–41.

6. Wang, Z., et al., A Never-Ending Story of Per- and Polyfluoroalkyl Substances (PFASs)? Environmental Science & Technology, 2017. 51(5): p. 2508–2518.

7. Glüge, J., et al., An overview of the uses of per- and polyfluoroalkyl substances (PFAS). Environmental Science: Processes & Impacts, 2020. 22(12): p. 2345–2373.

8. Schymanski, E.L., et al., Per- and Polyfluoroalkyl Substances (PFAS) in PubChem: 7 Million and Growing. Environmental Science & Technology, 2023. 57(44): p. 16918–16928.

9. Xiao, F., et al., Cross-national challenges and strategies for PFAS regulatory compliance in water infrastructure. Nature Water, 2023. 1(12): p. 1004–1015.

10. Hansen, S.F., et al., Late lessons from early warnings on PFAS. Nature Water, 2024. 2(12): p. 1157–1165.

11. Fenton, S.E., et al., Per- and Polyfluoroalkyl Substance Toxicity and Human Health Review: Current State of Knowledge and Strategies for Informing Future Research. Environ Toxicol Chem, 2021. 40(3): p. 606–630.

12. Hekster, F.M., R.W. Laane, and P. de Voogt, Environmental and toxicity effects of perfluoroalkylated substances. Rev Environ Contam Toxicol, 2003. 179: p. 99–121.

13. Wackett, L.P., Nothing lasts forever: understanding microbial biodegradation of polyfluorinated compounds and perfluorinated alkyl substances. Microbial biotechnology, 2022. 15(3): p. 773–792.

14. Wackett, L.P. and S.L. Robinson, A prescription for engineering PFAS biodegradation. Biochemical Journal, 2024. 481(23): p. 1757–1770.

15. Che, S., et al., Bird’s-Eye View: Current Understanding and Future Perspectives on the Biodefluorination of Per-and Polyfluoroalkyl Substances (PFAS). Water Research X, 2025: p. 100356.

16. Yu, Y., et al., Electron bifurcation and fluoride efflux systems implicated in defluorination of perfluorinated unsaturated carboxylic acids by Acetobacterium spp. Science Advances, 2024. 10(29): p. eado2957.

17. Wang, N., et al., 5:3 Polyfluorinated acid aerobic biotransformation in activated sludge via novel "one-carbon removal pathways". Chemosphere, 2012. 87(5): p. 527–34.

18. Wu, C., et al., Distinctive biotransformation and biodefluorination of 6:2 versus 5:3 fluorotelomer carboxylic acids by municipal activated sludge. Water Research, 2024. 254: p. 121431.

19. Geng, F. and D.E. Helbling, Cascading Pathways Regulate the Biotransformations of Eight Fluorotelomer Acids Performed by Wastewater Microbial Communities. Environmental Science & Technology, 2024. 58(52): p. 23201–23211.

20. Wu, C. and M. Li, Enriching fluorotelomer carboxylic acids-degrading consortia from sludges and soils. Science of The Total Environment, 2025. 958: p. 177823.

21. Tseng, N., et al., Biotransformation of 6:2 Fluorotelomer Alcohol (6:2 FTOH) by a Wood-Rotting Fungus. Environmental Science & Technology, 2014. 48(7): p. 4012–4020.

22. Zhang, H., et al., Biotransformation of 6:2 fluorotelomer alcohol by the whole soybean (Glycine max L. Merrill) seedlings. Environmental Pollution, 2020. 257: p. 113513.

23. Bhardwaj, S., et al., Biotransformation of 6:2/4:2 fluorotelomer alcohols by Dietzia aurantiaca J3: Enzymes and proteomics. Journal of Hazardous Materials, 2024. 478: p. 135510.

24. Fasano, W.J., et al., Absorption, Distribution, Metabolism, and Elimination of 8-2 Fluorotelomer Alcohol in the Rat. Toxicological Sciences, 2006. 91(2): p. 341–355.

25. Nabb, D.L., et al., In Vitro Metabolism of 8-2 Fluorotelomer Alcohol: Interspecies Comparisons and Metabolic Pathway Refinement. Toxicological Sciences, 2007. 100(2): p. 333–344.

26. Fasano, W.J., et al., Kinetics of 8-2 fluorotelomer alcohol and its metabolites, and liver glutathione status following daily oral dosing for 45 days in male and female rats. Chemico-Biological Interactions, 2009. 180(2): p. 281–295.

27. Martin, J.W., S.A. Mabury, and P.J. O’Brien, Metabolic products and pathways of fluorotelomer alcohols in isolated rat hepatocytes. Chemico-Biological Interactions, 2005. 155(3): p. 165–180.

28. Mothersole, R.G., et al., Formation of CoA Adducts of Short-Chain Fluorinated Carboxylates Catalyzed by Acyl-CoA Synthetase from Gordonia sp. Strain NB4-1Y. ACS Omega, 2023. 8(42): p. 39437–39446.

29. Zhang, S., et al., Biotransformation potential of 6: 2 fluorotelomer sulfonate (6: 2 FTSA) in aerobic and anaerobic sediment. Chemosphere, 2016. 154: p. 224–230.

30. Shaw, D.M., et al., Degradation and defluorination of 6: 2 fluorotelomer sulfonamidoalkyl betaine and 6: 2 fluorotelomer sulfonate by Gordonia sp. strain NB4- 1Y under sulfur-limiting conditions. Science of the Total Environment, 2019. 647: p. 690–698.

31. Allred, B.M., et al., Orthogonal zirconium diol/C18 liquid chromatography-tandem mass spectrometry analysis of poly and perfluoroalkyl substances in landfill leachate. J Chromatogr A, 2014. 1359: p. 202–11.

32. Fuertes, I., et al., Perfluorinated alkyl substances (PFASs) in northern Spain municipal solid waste landfill leachates. Chemosphere, 2017. 168: p. 399–407.

33. Lang, J.R., et al., National estimate of per-and polyfluoroalkyl substance (PFAS) release to US municipal landfill leachate. Environmental science & technology, 2017. 51(4): p. 2197–2205.

34. Wu, C., et al., Rapid quantitative analysis and suspect screening of per-and polyfluorinated alkyl substances (PFASs) in aqueous film-forming foams (AFFFs) and municipal wastewater samples by Nano-ESI-HRMS. Water Research, 2022. 219: p. 118542.

35. Shi, G., et al., 6:2 fluorotelomer carboxylic acid (6:2 FTCA) exposure induces developmental toxicity and inhibits the formation of erythrocytes during zebrafish embryogenesis. Aquat Toxicol, 2017. 190: p. 53–61.

36. Phillips, M.M., et al., Fluorotelomer acids are more toxic than perfluorinated acids. Environmental science & technology, 2007. 41(20): p. 7159–7163.

37. Yang, S.-H., et al., Desulfonation and defluorination of 6: 2 fluorotelomer sulfonic acid (6: 2 FTSA) by Rhodococcus jostii RHA1: Carbon and sulfur sources, enzymes, and pathways. Journal of Hazardous Materials, 2022. 423: p. 127052.

38. Yang, S.-H., L. Shan, and K.-H. Chu, Root exudates enhanced 6: 2 FTOH defluorination, altered metabolite profiles and shifted soil microbiome dynamics. Journal of Hazardous Materials, 2024. 466: p. 133651.

39. McLeod, M.P., et al., The complete genome of Rhodococcus sp. RHA1 provides insights into a catabolic powerhouse. Proceedings of the National Academy of Sciences, 2006. 103(42): p. 15582–15587.

40. Bygd, M.D., et al., Unexpected Mechanism of Biodegradation and Defluorination of 2,2-Difluoro-1,3-Benzodioxole by Pseudomonas putida F1. mBio, 2021. 12(6): p. e03001–21.

41. Finkelstein, Z.I., et al., Identification of Fluoropyrogallols as New Intermediates in Biotransformation of Monofluorophenols in Rhodococcus opacus 1cp. Applied and Environmental Microbiology, 2000. 66(5): p. 2148–2153.

42. Newton, G.L., N. Buchmeier, and R.C. Fahey, Biosynthesis and functions of mycothiol, the unique protective thiol of Actinobacteria. Microbiol Mol Biol Rev, 2008. 72(3): p. 471–94.

43. Anderberg, S.J., G.L. Newton, and R.C. Fahey, Mycothiol Biosynthesis and Metabolism: CELLULAR LEVELS OF POTENTIAL INTERMEDIATES IN THE BIOSYNTHESIS AND DEGRADATION OF MYCOTHIOL IN <EM>MYCOBACTERIUM SMEGMATIS</EM> *. Journal of Biological Chemistry, 1998. 273(46): p. 30391–30397.

44. Newton, G.L., Y. Av-Gay, and R.C. Fahey, A Novel Mycothiol-Dependent Detoxification Pathway in Mycobacteria Involving Mycothiol S-Conjugate Amidase. Biochemistry, 2000. 39(35): p. 10739–10746.

45. Dalleau, S., et al., Cell death and diseases related to oxidative stress:4-hydroxynonenal (HNE) in the balance. Cell Death & Differentiation, 2013. 20(12): p. 1615–1630.

46. Newton, G.L. and R.C. Fahey, Mycothiol biochemistry. Archives of microbiology, 2002. 178: p. 388–394.

47. Jackson, P.A., et al., Covalent Modifiers: A Chemical Perspective on the Reactivity of α,β-Unsaturated Carbonyls with Thiols via Hetero-Michael Addition Reactions. J Med Chem, 2017. 60(3): p. 839–885.

48. Ahangarpour, M., I. Kavianinia, and M.A. Brimble, Thia-Michael addition: the route to promising opportunities for fast and cysteine-specific modification. Org Biomol Chem, 2023. 21(15): p. 3057–3072.

49. Bürger, M. and J. Chory, Structural and chemical biology of deacetylases for carbohydrates, proteins, small molecules and histones. Communications Biology, 2018. 1(1): p. 217.

50. Pushkin, A., et al., Structural characterization, tissue distribution, and functional expression of murine aminoacylase III. American Journal of Physiology-Cell Physiology, 2004. 286(4): p. C848–C856.

51. Jiang, C. and B. Wu, Molecular cloning and functional characterization of a novel decarboxylase from uncultured microorganisms. Biochem Biophys Res Commun, 2007. 357(2): p. 421–6.

52. Strauss, E., et al., Mechanistic Studies on Phosphopantothenoylcysteine Decarboxylase: Trapping of an Enethiolate Intermediate with a Mechanism-Based Inactivating Agent. Biochemistry, 2004. 43(49): p. 15520–15533.

53. Paul, B.D. and S.H. Snyder, Therapeutic Applications of Cysteamine and Cystamine in Neurodegenerative and Neuropsychiatric Diseases. Frontiers in Neurology, 2019. 10.

54. Gallego-Villar, L., et al., Cysteamine revisited: repair of arginine to cysteine mutations. J Inherit Metab Dis, 2017. 40(4): p. 555–567.

55. Clausen, T., et al., Slow-Binding Inhibition of Escherichia coli Cystathionine β-Lyase by l-Aminoethoxyvinylglycine: A Kinetic and X-ray Study. Biochemistry, 1997. 36(41): p. 12633–12643.

56. Tateishi, M., S. Suzuki, and H. Shimizu, Cysteine conjugate beta-lyase in rat liver. A novel enzyme catalyzing formation of thiol-containing metabolites of drugs. J Biol Chem, 1978. 253(24): p. 8854–9.

57. Matoba, Y., et al., Catalytic specificity of the Lactobacillus plantarum cystathionine γ- lyase presumed by the crystallographic analysis. Scientific Reports, 2020. 10(1): p. 14886.

58. Wang, N., et al., 5: 3 Polyfluorinated acid aerobic biotransformation in activated sludge via novel “one-carbon removal pathways”. Chemosphere, 2012. 87(5): p. 527–534.

59. Jayasinghe, Y.P., et al., The Mycobacterium tuberculosis mycothiol S-transferase is divalent metal-dependent for mycothiol binding and transfer. RSC Medicinal Chemistry, 2023. 14(3): p. 491–500.

60. Newton, G.L., et al., The DinB Superfamily Includes Novel Mycothiol, Bacillithiol, and Glutathione S-Transferases. Biochemistry, 2011. 50(49): p. 10751–10760.

61. Fan, F., et al., Structures and mechanisms of the mycothiol biosynthetic enzymes. Curr Opin Chem Biol, 2009. 13(4): p. 451–9.

62. Metaferia, B.B., et al., Synthesis of Natural Product-Inspired Inhibitors of Mycobacterium tuberculosis Mycothiol-Associated Enzymes: The First Inhibitors of GlcNAc-Ins Deacetylase. Journal of Medicinal Chemistry, 2007. 50(25): p. 6326–6336.

63. Lamprecht, D.A., et al., An enzyme-initiated Smiles rearrangement enables the development of an assay of MshB, the GlcNAc-Ins deacetylase of mycothiol biosynthesis. Org Biomol Chem, 2012. 10(27): p. 5278–88.

64. Maynes, J.T., et al., The crystal structure of 1-D-myo-inosityl 2-acetamido-2-deoxy-alpha-D-glucopyranoside deacetylase (MshB) from Mycobacterium tuberculosis reveals a zinc hydrolase with a lactate dehydrogenase fold. J Biol Chem, 2003. 278(47): p. 47166–70.

65. Whittenbury, R., K.C. Phillips, and J.F. Wilkinson, Enrichment, Isolation and Some Properties of Methane-utilizing Bacteria. Microbiology, 1970. 61(2): p. 205–218.

66. Newton, G.L., P. Ta, and R.C. Fahey, A mycothiol synthase mutant of Mycobacterium smegmatis produces novel thiols and has an altered thiol redox status. Journal of bacteriology, 2005. 187(21): p. 7309–7316.

67. Young, R.B., et al., PFAS Analysis with Ultrahigh Resolution 21T FT-ICR MS: Suspect and Nontargeted Screening with Unrivaled Mass Resolving Power and Accuracy. Environmental Science & Technology, 2022. 56(4): p. 2455–2465.

68. Chothia, C. and A.M. Lesk, The relation between the divergence of sequence and structure in proteins. Embo j, 1986. 5(4): p. 823–6.

69. Tamura, K., G. Stecher, and S. Kumar, MEGA11: Molecular Evolutionary Genetics Analysis Version 11. Mol Biol Evol, 2021. 38(7): p. 3022–3027.

70. Jones, D.T., W.R. Taylor, and J.M. Thornton, The rapid generation of mutation data matrices from protein sequences. Bioinformatics, 1992. 8(3): p. 275–282.

71. Crooks, G.E., et al., WebLogo: a sequence logo generator. Genome Res, 2004. 14(6): p. 1188–90.

