## Supporting Information for "Mycothiol Conjugation Drives Biotransformation and Biodefluorination of Fluorotelomer Carboxylic Acids by *Actinomycetota* and Sludge Microbiomes"

for

### S1. Methods and Materials

#### S1.1 Analytical Control (An) and Laboratory Reagent Blank (LRB) Preparation

Analytical controls (Ans) consisted of 19 mL reagent water, 1 mL AMS solution, and 20 μL methanolic PFAS analyte solutions (i.e., 6:2 FTCA, 5:3 FTCA, and the mixture of 6:2 and 5:3 FTCA) to achieve a final analyte concentration of 40 μM, without live or killed cells added.

Laboratory reagent blanks (LRBs) were prepared with: (1) 19 mL reagent water, 1 mL AMS solution, and 13.3 μL of methanol (MeOH); (2) 19 mL reagent water, 1 mL AMS solution, and 13.3 μL of acetonitrile (MeCN); and (3) 19 mL of reagent water and 1 mL of AMS solution. Both Ans and LRBs were prepared in triplicate, with processing, preservation, and storage identical to that of the treatment groups.

#### S1.2 Abiotic Control (AC) and Microbial Blank (MB) Preparation

Abiotic (killed) controls (ACs) were prepared similarly to the treatment groups, using autoclaved bacterial cells at $OD_{600} \approx 1.0$ to assess abiotic loss, such as adsorption and evaporation. The culture stock was sterilized using a Hirayama HICLAVE HVE-50 Autoclave from Marshall Scientific (Hampton, NH, USA). Killed cells were washed and adjusted to a final volume of 20 mL with 20× diluted AMS medium. Four sets of ACs were prepared in triplicate, including two spiked with individual FTCA stocks, one spiked with the mixture of both FTCAs, and one blank without FTCAs added.

Microbial blanks (MBs) were set up using resting cells prepared identically to the treatment groups. Washed live cells were resuspended with diluted AMS to a final volume of 20 mL with 13.3 μL of pure methanol added.

Both ACs and MBs were centrifuged after each exposure cycle, and the supernatants were recycled for the subsequent batches. Cell pellets were discarded after each round.

**S1.3 HRMS Data Preprocessing**

HRMS data preprocessing was conducted to eliminate background noise from analyte signals and extract mass features before further analysis (e.g., suspect screening, KMD, and non-target analysis for the iterative TP accumulation assays). Raw MS1 spectra were extracted from Thermo Xcalibur™ software as Microsoft Excel files. Background correction was performed against five types of controls/blanks (Table S4), including LRBs, ACs, MBs, Ans, and backgrounds (Bgs). The following notation and procedures were employed.

$m_{E,i}$, $I_{E,i}$: The *m/z* value and corresponding intensity of the $i^{\text{th}}$ mass feature in the treatment spectrum;

$m_{C,j}^{(b)}$, $I_{C,j}^{(b)}$: The *m/z* value and corresponding intensity of the $j^{\text{th}}$ mass feature in control/blank set $b$, where $b \in \{LRB, AC, MB, An, Bg\}$;

$I_{IS,E,i}$, $I_{IS,C,j}^{(b)}$: Internal standard intensities for treatment and control features;

Indicator function: $\tau(x) = \begin{cases} 1, & if\ x\ is\ true, \\ 0, & otherwise, \end{cases} \quad (Eq. s1)$

$RI_{E,i}$, $RI_{C,j}^{(b)}$: Relative intensities for features in treatment and control/blank spectrum, where

$$RI_{E,i} = \frac{I_{E,i}}{I_{IS,E,i}}, RI_{C,j}^{(b)} = \frac{I_{C,j}^{(b)}}{I_{IS,C,j}^{(b)}} \quad (Eq. s2)$$

(a) Signal-to-Noise Ratio (SNR) Filtering

Peaks with SNR > 3 were retained in all spectra as positive detection; a value greater than 3 generally indicates the detected signal is unlikely a noise with 99.7% confidence.

$$\sigma_{E,i} = \tau(SNR_{E,i} > 3), \sigma_{C,j}^{(b)} = \tau\left(SNR_{C,j}^{(b)} > 3\right) \quad (Eq. s3)$$

(b) Scan Reproducibility Filtering

Features in treatment samples recorded as positive signals must appear in ≥ 50 scans out of 75 total scans across all triplicate runs. For control/blanks, the threshold was reduced to ≥ 25 scans out of 75 total scans in at least two of triplicate runs to increase the scrutiny of the screening.

$$\eta_{E,i}^{(r)} = \tau\left(\frac{scan_{E,i}^{(r)}}{75} \geq \frac{50}{75}\right), \quad \eta_{C,j}^{(b,r)} = \tau\left(\frac{scan_{C,j}^{(b,r)}}{75} \geq \frac{25}{75}\right), \quad (r = 1,2,3) \quad (Eq. s4)$$

$$\eta_{E,i} = \prod_{r=1}^{3} \eta_{E,i}^{(r)}, \quad \eta_{C,j}^{(b)} = \tau\left(\sum_{r=1}^{3} \eta_{C,j}^{(b,r)} \geq 2\right) \quad (Eq. s5)$$

(c) Control/Blank Feature Collection

Control/blank features were identified when meeting (a) and (b) filtration criteria and having RI > 0.01% and intensity < 2e3.

$$\Omega^{(b)} = \left\{j \,\middle|\, \sigma_{C,j}^{(b)} \cdot \eta_{C,j}^{(b)} \cdot \tau\left(RI_{C,j}^{(b)} > 0.01\%\right) \cdot \tau\left(I_{C,j}^{(b)} < 2000\right) = 1\right\} \quad (Eq. s6)$$

(d) Mass Matching Filtering

Mass features with anionic *m/z* values within ±5 ppm interval ($\epsilon_m = 5 \times 10^{-6}\ Da/Da$) were matched across measurements for subsequent analysis.

$$\Gamma_i = \left\{(b,j) \,\middle|\, \left|m_{E,i} - m_{C,j}^{(b)}\right| < \epsilon_m, j \in \Omega^{(b)}\right\} \; or \; \Gamma_i = \left\{\left|m_{E,i} - m_{E,ii}\right| < \epsilon_m\right\} \quad (Eq.s7)$$

And if a matched mass pair exists between treatment group and controls/blanks,

$$\mu_{E,i} = \tau\left(\min_{(b,j)\in\Gamma_i}\left\{\left|m_{E,i} - m_{C,j}^{(b)}\right|\right\} < \epsilon_m\right), \quad \Gamma_i \neq \emptyset \quad (Eq.s8)$$

otherwise,

$$\mu_{E,i} = 1, \quad \Gamma_i = \emptyset \quad (Eq.s9)$$

(e) RI Filtering

Mass features in treatment groups required a minimum RI threshold of 0.01% in at least two of triplicates. Matched mass features (d) were accepted when their mean RIs in treatments were at least 50-fold higher than the corresponding matched control features.

$$\rho_{E,i}^{(r)} = \tau\left(RI_{E,i}^{(r)} > 0.01\%\right) \quad \rho_{E,i} = \tau\left(\sum_{r=1}^{3} \rho_{E,i}^{(r)} \geq 2\right) \quad (Eq.s10)$$

If matched controls/blanks exist,

$$\theta_{E,i} = \tau\left(\overline{RI_{E,i}} > 50 \times \max_{(b,j)\in\Gamma_i}\left\{RI_{C,j}^{(b)}\right\}\right) \bigwedge \rho_{E,i}, \quad \Gamma_i \neq \emptyset \quad (Eq.s11)$$

If there are no matched controls/blanks,

$$\theta_{E,i} = \rho_{E,i}, \quad \Gamma_i = \emptyset \quad (Eq.s12)$$

(f) Intensity Cutoff

Any mass features with any matching control/blank features having intensity >2e3 were strictly removed regardless of the feature's intensity/RI in treatment triplicate.

$$\kappa_i = \prod_{(b,j)\in\Gamma_i} \tau\left(I_{C,j}^{(b)} < 2000\right) \quad (Eq.s13)$$

As a result, an experimental mass feature $i$ was retained only if it satisfied all criteria ($\Phi_{E,i} = 1$):

$$\Phi_{E,i} = \sigma_{E,i} \cdot \eta_{E,i} \cdot \mu_{E,i} \cdot \theta_{E,i} \cdot \kappa_i \quad (Eq.s14)$$

Features with an RI < 0.5% that passed these filters were retained as potential qualitative references but excluded from further MS2 analyses.

The data preprocessing algorithm was implemented through a custom Python script available at https://github.com/Daitoueqaq/HRMS-modification, and executed in JupyterLab on a local workstation.

**S1.4 Non-target Screening for TP Accumulation through Iterative Exposure Cycles**

Our iterative non-target screening workflow employed an index-based statistical approach to identify enrichment of mass ions across exposure cycles. After HRMS data preprocessing, candidate features were evaluated based on two indices: the Cumulative Significance Index (CSI) and the Linearity Trend Index (LTI), as quantitative indicators of statistically significant RI changes and linear accumulation trend over enrichment cycles, respectively. These indices were integrated into a final Accumulation Score, $S(n)$, for each feature. Candidate features with a score exceeding the threshold $\xi$ were collected as "mass ions of interest" (MIIs).

A matching mass error within 5 ppm ($Eq.s5$) was used to pair features across iterative exposure cycles.

(a) Cumulative Significance Index (CSI, $\varphi_{CSI}(n)$)

The CSI quantifies the statistical significance of RI changes of features at the cycle $n$ compared to each preceding cycle $s$ ($s = 1, 2, \ldots, n-2, n-1$). Pairwise $t$ tests of the triplicate RI measurements were performed for each feature and $p$-values ($p_{n,s}$) were obtained. The Benjamini–Hochberg (BH) correction procedure was employed at a significance threshold of $\alpha = 0.05$.

$$\underbrace{R_{n,s}}_{"pseudo-ranking"} = 1 + \sum_{t=1}^{n-1} H(p_{n,t} - p_{n,s}), \quad q_{n,s} = \frac{(n-1)p_{n,s}}{R_{n,s}} \qquad (Eq.s15)$$

where $q_{n,s}$ represent BH-adjusted $p$-value, and $H(z)$ is the Heaviside step function. The normalized CSI was calculated as:

$$\varphi_{CSI}(n) = \frac{1}{(n-1)[-ln(\alpha)]} \sum_{s=1}^{n-1} [-\ln(q_{n,s}) \cdot H(\alpha - q_{n,s})] \qquad (Eq.s16)$$

$\varphi_{CSI}(n)$ estimates the aggregate strength of how significantly the RI of each feature at cycle $n$ differs from previous cycles, irrespective of direction. The index can sensitively detect both accumulations and transient metabolic behaviors.

(b) Linearity Trend Index (LTI, $\varphi_{LTI}(n)$)

The LTI identifies features demonstrating a sustained accumulation trend over exposure cycles. A linear regression model is fitted to the feature's RIs across cycles. The slope ($m$) indicating the TP accumulation rate, and the coefficient of determination ($R^2$) reflecting the linearity of the accumulation trend, are extracted. The LTI combines two metrics to evaluate the linear trend of each feature, and the slope is normalized using the hyperbolic tangent function (tanh).

$$\varphi_{LTI}(n) = H(m) \cdot \tanh(m) \cdot R^2 \quad (Eq.s17)$$

$\varphi_{LTI}(n)$ is suitable for features demonstrating progressive enrichment across cycles but requires a minimum of three datapoints ($n \geq 3$) to yield a meaningful $R^2$.

(c) Accumulation score $S(n)$

The final accumulation score combined two indices:

$$S(n) = \delta_1 \varphi_{CSI}(n) + \delta_2 \varphi_{LTI}(n), \quad (\delta_1 + \delta_2) = 1 \quad (Eq.s18)$$

Weighting factors $\delta_1$, $\delta_2$ were used to prioritize the contribution of statistical significance, while incorporating linear trend. Features were classified as MIIs when $S(n)$ exceeded the threshold $\xi$:

$$\chi = H(S(n) - \xi) \quad (Eq.s19)$$

In this study, we used weighting factor set below to screen for MIIs in iterative TP enrichment experiments.

| Analysis | $\delta_1$ | $\delta_2$ | $\xi$ |
|---|---|---|---|

| | | | |
|---|---|---|---|
| Iterative TP Enrichment | 0.70 | 0.30 | 0.25 |

The overall equation for the MII identification:

$$\chi = H\left(\delta_1 \frac{1}{(n-1)[-ln(\alpha)]}\sum_{s=1}^{n-1}\left[-\ln\left(q_{n,s}\right)\cdot H\left(\alpha - q_{n,s}\right)\right] + \delta_2 H(m)\cdot \tanh(m)\cdot R^2 - \xi\right) \quad (Eq. s20)$$

For single-cycle (0-48 h) experiments, the workflow was also used to detect potential TPs, both accumulating and transient. Therefore, the LTI was less critical for detection of short-lived TPs than CSI. Accordingly, the weighting factors were adjusted to $\delta_1 = 0.85$, $\delta_2 = 0.15$ with the same threshold ($\xi = 0.25$).

**S1.5 Suspect Screening**

The suspect screening procedure in this study was developed following previous studies [1, 2], consisting of 4 stages: local database construction, background correction (spectra preprocessing), identification of potential matches and structural validation [2].

Our local database integrates the U.S. EPA's PFAS MASTER list and the PubChem PFAS Tree. The suspect list was routinely updated with newly identified PFAS from recent publications [1-7]. The PFAS MASTER list (available at https://comptox.epa.gov/dashboard/chemical-lists/pfasmaster) and PubChem PFAS Tree (available at https://pubchem.ncbi.nlm.nih.gov/classification/#hid=120) provide a comprehensive catalog for PFASs and other fluorinated compounds. The database consists of detailed information including molecular formulas, identifiers (DTX-SID, CAS RN, IUPAC name), structural codes (SMILES, InChI), average molecular weights, monoisotopic molecular weights, putative monoisotopic anionic weights, and data source.

Features after background correction were compared against the local PFAS database with a mass error threshold of 5 ppm ($5\times10^{-6}$). Features exhibiting a RI below 0.5% were disregarded since a low signal typically lack the reliability for obtaining MS2 spectra. All positive hits that met the criteria were collected and ranked by intensity.

Molecular structures of potential hits were subjected to CID to obtain their fragmentation patterns derived from MS2 spectra.

*Confidence level*

Confidence levels associated with the structure of newly identified PFASs were determined with a combination of MS1 spectra, MS2 fragmentation pattern, library MS2 profiles, and the availability of reference standards. The evaluation follows the guidelines by Schymanski et al. [8] and Charbonnet et al. [9].

**S1.6 Semi-Quantification Analysis**

A semi-quantification (SQ) approach following established methodologies from previous studies [2, 3, 10-12] was used to estimate the concentration of newly discovered conjugated metabolites. Due to the unavailability of commercial standards for these conjugated metabolites, we selected calibration standards that share structural and physicochemical properties with the analytes [13]. As reported in previous studies [14, 15], an effective calibration standard typically exhibits structural homology, similar functional groups influencing ionization response and molecular weights. Calibration standards were assigned to metabolites to enhance SQ accuracy:

(1) The glutathione conjugate of monobromobimane (GS-mBBr, SI Section S1.11) was selected as the calibration standard for MSH-derived conjugates, including MS-5:3 FTUCA, AcCyS-5:3 FTUCA, AcCyS-5:3 FTCA, CyS-5:3 FTUCA, and other downstream products. The choice is justified by several similarities, since (i) both GS-mBBr and newly identified MSH conjugates contain the thiol-derived sulfur linkage (i.e., the thioether bond) as the major structural feature; (ii) mercapturic acid metabolites and GS-mBBr both contain two terminal carboxyl groups (-COOH) serving as the primary sites of deprotonation under negative-ion electrospray conditions; (iii) all molecules are products via conjugation of a thiol compound, resulting in a similar backbone; (iv) GS-mBBr (498 g/mol) shares a close molecular weight to those of the major analytes (e.g., AcCyS-5:3 FTUCA *m/z* 500.0026; AcCyS-5:3 FTCA *m/z* 500.0182; CyS-5:3 FTUCA *m/z* 457.9920; AcCyS-4:3 FTUCA *m/z* 450.0058). Although some downstream thiol-containing TPs contain a single carboxyl group, the core thioether linkage remains.

(2) 5:3 FTCA was selected as the calibration standard of fluorotelomer intermediates, such as 5:3 FTUCA, OH-5:3 FTCA, OH-4:3 FTCA. Structural similarities, including ionizable carboxylate group and similar fluorotelomer backbone, provided basis for yielding a similar negative-ion ESI response.

(3) Perfluorobutyric acid (PFBA) standard was used as the calibration standard to semi-quantify PFPrA.

Two mass-labeled internal standards (IS) were employed during analysis. M8-PFOA

($^{13}C_8$-PFOA) was used as the IS for perfluorinated carboxylates (PFCAs) quantified in target analysis by LC/MS/MS. M2-6:2 FTCA ($^{13}C_2$-6:2 FTCA) was used as the IS for all fluorotelomer-based metabolites quantified by Nano-ESI-HRMS, including thiol-conjugates and fluorotelomer intermediates. All IS were spiked right before analysis. All calibrations were established using the peak area ratio of the analyte to their corresponding IS (Figure S38).

#### S1.7 Fluorine Mass Recovery (FMR)

The fluorine mass recovery (FMR) was calculated following previous studies [12, 16] for fluorine mass balance. The FMR represents the percentage of the sum of measurable fluorine species retained and transformed in the biological assays, which consist of all organofluoride (OF) compounds and released fluoride ions, relative to the theoretical total amount of fluorine (initial fluorine dose) in the system. An abiotic defect correction (AD) accounts for any potential fluorine loss due to abiotic processes. The AD is determined by subtracting the total fluorine measured in the AC from the An at the end of the experiment.

$$AD = \left[\sum\left(C_{OF} \times \frac{F\ atom}{molecule}\right) + C_{free\ F^-}\right]_{An} - \left[\sum\left(C_{OF} \times \frac{F\ atom}{molecule}\right) + C_{free\ F^-}\right]_{AC}$$

$$(Eq. s21)$$

The FMR for the treatment groups was calculated by the sum of all measured fluorine-containing species (all detected thiol conjugated metabolites [Conj.] and metabolites from the "one-carbon removal pathways" [OCRP]) and the AD factor, which was normalized to the initial fluorine dose.

$$FMR = \frac{\left[\sum\left(C_{Conj.} \times \frac{F\ atom}{molecule}\right) + \sum\left(C_{OCRP} \times \frac{F\ atom}{molecule}\right) + C_{free\ F^-}\right]_{Treatment} + AD}{\left[\sum\left(C_{OF} \times \frac{F\ atom}{molecule}\right) + C_{free\ F^-}\right]_{An}}$$

$$(Eq. s22)$$

**S1.8 Protein Structure Prediction by AlphaFold2**

AlphaFold2 algorithm was employed for the prediction of three-dimensional structures of the essential enzymes (i.e., MST, Mca, and MshA-D) for *Actinomycetota* representatives. The predictions were accessed via the AlphaFold2 ColabFold notebook (https://colab.research.google.com/github/sokrypton/ColabFold/blob/main/AlphaFold2.ipynb).

Amino acid sequences were obtained from the NCBI database from eight representative *Actinomycetota* species. Multiple-sequence alignments (MSAs) were generated with MMseqs2, and the running parameters are listed below.

| Key Parameter for Alphafold2 ColabFold | Setting |
|---|---|
| Number of relax iterations | 5 |
| Recycles | 24 (MST); 12 (Mca, MshA-D) |
| Pairing strategy | Complete |
| Seeds per target | 4 |

The model with the highest mean per-residue confidence score (pLDDT) was collected from the set of predictions for downstream analysis.

The predicted protein structures were compared with the known crystal structure of *Mycobacterium tuberculosis* H37Rv, and among *Rhodococcus jostii* RHA1, and other MSH-producing representative bacteria (Figure S13-S15). Pairwise structure alignments were performed with PyMOL (Schrödinger, Inc.) to assess the structural conservation. Structural similarities were quantified by the alpha-carbon (Cα) root-mean-square deviation (RMSD) [17]. A lower RMSD (< 2.0 Å) typically indicates a highly conserved protein topology.

#### S1.9 Kendrick Mass Defect (KMD) Analysis

The KMD workflow was applied to the deconvoluted MS1 feature list after data preprocessing ($Eq.\ s14$). The mass of each feature was converted to its Kendrick mass (KM) following the equation below that normalize the extracted mass $m_{E,i}$ by the ratio of the nominal mass $m_n$ and the exact mass $m_e$ relative to a repeating unit (e.g., $CF_2$).

$$KM = \frac{m_n}{m_e} m_{E,i} \quad (Eq. s23)$$

Then, the KMD is the difference between the feature's KM and its nominal KM:

$$KMD = KM - nint(KM) \quad (Eq. s24)$$

where $nint()$ is the nearest integer function.

The primarily KMD investigated $CF_2$ as the repeating unit ($m_e$= 49.99681 Da, $m_n$= 50 Da, Figure 3g), and additional KMD calculations were performed using other units detailed in the table below to capture chain variations, hetero-substitutions, and isotopologues.

| Repeating Unit | $m_e$ (Da) | $m_n$ (Da) |
|---|---|---|
| $CF_2$ | 49.99681 | 50 |
| $CH_2$ | 14.01565 | 14 |
| $F_2$ | 37.99681 | 38 |
| $H_2$ | 2.01565 | 2 |

| | | |
|---|---|---|
| +F-H | 17.99058 | 18 |
| $C_2H_4O$ | 44.02621 | 44 |
| $^{13}C/^{12}C$ | 1.00335 | 1 |
| $^{34}S/^{32}S$ | 1.99580 | 2 |

A KMD map was constructed for each unit, and homolog clusters with at least 2 members aligned at similar KMD differences ($\Delta < 1\ mDa$) were attributed to MIIs [18]. The MII with the highest intensity in each identified homolog series was prioritized for MS2 fragmentation, and all MIIs will be monitored across (i) iterative enrichment cycles and (ii) 48-h resting cell assays. The script of the complete KMD procedure is available at https://github.com/Daitoueqaq/RHA1/blob/main/KMD.

#### S1.10 Resting-cell FTCA biotransformation by *M. smegmatis* MC²155, *M. dioxanotrophicus* PH-06, and *P. dioxanivorans* CB1190

Three additional *Actinomycetota* strains were tested for their ability to biotransform FTCAs and generate thiol-conjugated metabolites following the same resting cell assay protocol as RHA1. A single colony of MC²155, PH-06, or CB1190 was transferred to 100 mL glucose-supplemented AMS medium (with PH-06 and CB1190 incubated with 1,4-dioxane-supplemented AMS medium) and incubated (30 °C, 130 rpm) until exponential phase ($OD_{600} \approx 1.0$, ~40 h). Cultures were harvested by centrifugation (12,000 rpm, 4 °C, 15 min), washed three times, and resuspended in 20× diluted AMS. 5 mL of cell suspension was transferred to 60-mL amber serum bottles and amended with 14.25 mL Milli-Q water and 0.75 mL AMS. 20 µL of 40 mM 6:2- or 5:3-FTCA stock (MeOH/$H_2O$ 2:1) were spiked to achieve an initial concentration of 40 µM. Similar triplicate bottles were prepared for ACs, Ans,

MBs, LRBs, and Bgs. Bottles were incubated at 30 °C, 130 rpm for 48 h. Aliquots (0.5 mL) for HRMS analysis were collected at 0, 4, 16, 24, and 48 h, and samples were vortexed and centrifuged, and the supernatants were stored at –20 °C until analysis. Samples for fluoride analysis (2 mL) were collected at 24 and 48 h, filtered (0.22 μm PES), and mixed 1:1 with TISAB II buffer before detection.

**S1.11 Synthesis of GS-mBBr Conjugate**

Monobromobimane (mBBr) can transfer thiols (R-SH) into stable R-S-bimane through an $S_N2$ displacement of bromine. This reaction proceeds rapidly at pH ≥ 7.5 when the thiolate anion is the reactive nucleophile [19]. Glutathione (GSH) conjugate with mBBr yields GS-mBBr (Figure S36) with chromatographic behavior and ionization efficiency similar to those of mercapturic acid conjugates, as detailed in SI Section S1.6. The GS-mBBr conjugate was synthesized following established protocols from prior studies [19, 20]. A 5-mL reaction solution was set up in the dark, consisting of 40 mM HEPES (pH = 8.0) and 2 mM mBBr in aqueous acetonitrile. Reduced-GSH standard solution was added to a final concentration of 1 mM. The mixture was gently vortexed and incubated at 50 °C water bath for 15 min. The supernatant was collected after centrifugation and diluted 4× with 10 mM HCl to quench residual reagent. The half-life of GSH conjugation with mBBr is ~20 s with > 99 % transformation within the reaction time [19]. The GS-mBBr conjugate solution (250 μM) was sealed in amber vials and stored at –20 °C until use. Stability tests have indicated < 2 % decomposition over three months. The sample was passed through an Oasis™ SPE cartridge and resuspended in MeCN/H2O (1:1, v/v) to remove salts and excess reagent.

### S1.12 Analytical Calibration

A two-segment calibration model was built based on the intensity ratio of the calibration standard to the corresponding IS to semi-quantify the responses of each analyte. Concentrations were calculated using either a low-concentration segment (0.1 – 10 μg/L) or a high-concentration segment (5 – 200/300 μg/L) to account for potential nonlinearity response at high concentrations due to MS saturation or ion suppression [21]. Concentrations that were above 10 μg/L were calculated using the high-concentration segment. A weighted least-squares (WLS) linear regression (1/x weighting) was applied to improve quantification accuracy at lower concentrations (< 1 μg/L) and mitigate influence from intercept.

The analytical sensitivity was determined by the limit of detection (LOD) and limit of quantification (LOQ) (Table S22). They were calculated from the standard deviation ($\sigma$) of seven parallel measurements (i.e., RIs) of the standard prepared at 0.1 μg/L:

$$LOD = 3.3 \times \frac{\sigma}{m} \quad LOQ = 10 \times \frac{\sigma}{m} \quad (Eq.s25)$$

where $m$ is the slope of the WLS calibration. The LOD and LOQ for the semi-quantified metabolites were conservatively assumed to equal those of their respective calibration standard.

### S1.13 Target PFAS Analysis by LC/MS/MS

A liquid chromatography-tandem mass spectrometry (LC/MS/MS) method was adopted, following the EPA Method 1633 (US EPA), for the validation of "one-carbon removal pathways" intermediates and the quantification of target PFCAs (i.e., PFHxA, PFPeA, and

PFBA). Concentrations were determined using IS-corrected calibration curves. The analysis employed a 1290 Infinity II high-performance liquid chromatography (HPLC) system, coupled with a 6470A triple quadrupole mass spectrometer from Agilent Technologies, Inc. (Santa Clara, CA, USA). Samples were diluted by 50 times with MeOH, spiked with mass-labelled IS, and filtered by 0.22 μm PES filters before injection. Chromatographic separation was performed on a Symmetry® C18 column (2.1 mm ID, 100 mm length, 3.5 μm particle size) from Waters Corporation (Milford, MA, USA). Injection volume was set to 10 μL with the flow rate adjusted within a range of 0.30 to 0.35 mL/min. The mobile phase began with an initial composition of 95% solvent A (water phase, 2 mM ammonium acetate in water with 5% MeCN), transitioning to 40% of solvent B (100% MeCN) over 2.5 minutes. The composition was then modulated to 10% solvent A over the next 5 min, maintained for 1 min before reverting to 95% solvent A in 0.5 min. The mobile phase was then held for 2 min for the post-run period. Parameters of liquid chromatographic conditions are shown in Table S23.

The triple quadrupole (QQQ) mass spectrometer was operated in negative electrospray ionization mode with ion source parameters set as follows: desolvation gas temperature at 350 °C with a flow rate of 2 L/min, nebulizer pressure at 25 psi, and capillary voltage at -3.5 kV. Multiple reaction monitoring (MRM) mode was used for quantifying target PFAS analytes. Detailed mass spectrometric conditions are provided in Table S24. MS data analysis was conducted using the Agilent MassHunter Quantitative Analysis software.

### S2. Results and Discussions

#### S2.1 Molecular Mechanisms for Mycothiol (MSH)-mediated Detoxification

Mycothiol (MSH; AcCys-GlcN-Ins) is a significant low molecular weight (LMW) thiol synthesized at millimolar concentrations by *Actinomycetota*, such as *Mycobacterium*, *Rhodococcus*, *Streptomyces*, *Micromonospora*, and *Nocardiopsis* [22]. Functionally analogous to glutathione (GSH), which is absent in these organisms, MSH serves as the primary thiol compound for maintaining intracellular redox homeostasis and mitigating oxidative and disulfide stress [22-24]. The molecular structure of MSH (Figure S8) features a functional acetylated cysteine linked through an amide bond to a glucosaminyl-inositol (GlcN-Ins) moiety. This structure stabilizes the reactive cysteine, reducing its rate of auto-oxidation by preserving it in a chemically less reactive form [22, 25]. The amide linkage is the specific site of enzymatic hydrolysis by mycothiol S-conjugate amidase (Mca) [22, 24, 26].

MSH performs its antioxidant function through the scavenging of reactive oxygen species and other electrophiles by the nucleophilic thiol group (-SH) of its cysteine moiety [22, 24]. It is known to detoxify a number of thiol-reactive xenobiotics, including alkylating agents (e.g., formaldehyde), antibiotics (e.g., erythromycin), reactive oxygen and nitrogen species (e.g. hydrogen peroxide, menadione), and other toxic molecules such as cumene hydroperoxide [23, 27, 28].

The detoxification process mediated by MSH involves a two-step enzymatic cascade (Figure 3f) [22-24, 26, 29]. First, the nucleophilic thiol group of MSH conjugates with a xenobiotic (R), forming mycothiol S-conjugates (MS-R) catalyzed by mycothiol S-transferase (MST). Then the enzyme Mca specifically hydrolyzes the amide bond of the conjugate. This cleavage releases the mercapturic acid derivative (AcCyS-R) and the non-toxic GlcN-Ins moiety. The AcCyS-R conjugate is exported from the cell, and GlcN-Ins recycles intracellularly back to the MSH biosynthesis via GlcN-Ins ligase (MshC) and mycothiol synthase (MshD).

The enzymatic synthesis of MSH follows a conserved enzymatic pathway [22, 29]. The process initiates with the formation of 1-O-(2-acetamido-2-deoxy-alpha-D-glucopyranosyl)-1D-myo-inositol 3-phosphate (GlcNAc-Ins-P) from 1D-myo-inositol 3-phosphate and UDP-N-acetyl-alphaD-glucosamine (UDP-GlcNAc), catalyzed by glycosyltransferases MshA (Table S12). A putative phosphatase MshA2 dephosphorylates GlcNAc-Ins-P to yield GlcNAc-Ins, which is further deacetylated by MshB (Table S13) to produce GlcN-Ins. MshC (Table S14) catalyzes the ATP-dependent ligation of cysteine to GlcN-Ins via an amide bond, producing 1D-myo-inositol 2-(L-cysteiniumylamino)-2-deoxy-alpha-D-glucopyranoside (Cys-GlcN-Ins). MshD (Table S15) acetylates the amino group of the cysteine within Cys-GlcN-Ins using acetyl-CoA to form the MSH molecule. The cyclic conjugation and MSH regeneration process, together with the enzymatic biosynthesis of MSH, are detailed in Figure 3f.

*Rhodococcus jostii* RHA1 is a Gram-positive *Actinomycetota*, originally isolated from γ-hexachlorocyclohexane-contaminated soil, recognized for its potent capabilities in degrading a wide array of xenobiotics, including polychlorinated biphenyls (PCBs) [30] and o-xylene [31] through co-metabolism. RHA1 is exemplified as a model organism for investigating *Actinomycetota*-mediated biodegradation due to its extensive repertoire of catabolic pathways [32]. The synthesis of MSH within RHA1 is crucial for mitigating the oxidative stress induced by the metabolism of toxic substances.

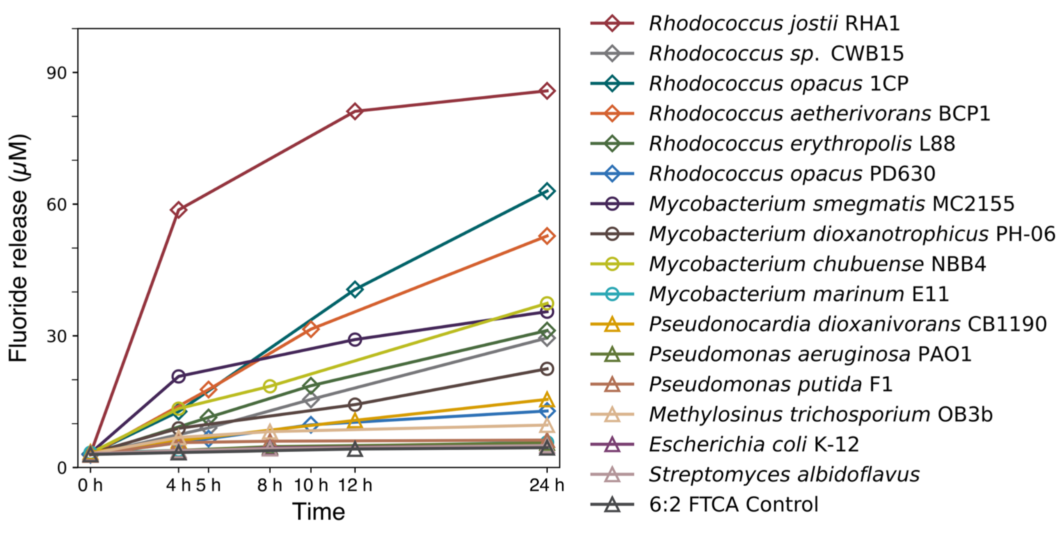


**Figure S1.** Biodefluorination comparisons among Gram-positive and Gram-negative cultures against 6:2 FTCA. Fluoride release after exposure to 6:2 FTCA (40 µM) for 24 h. All cultures were inoculated at an initial $OD_{600}$ ≈1.0.

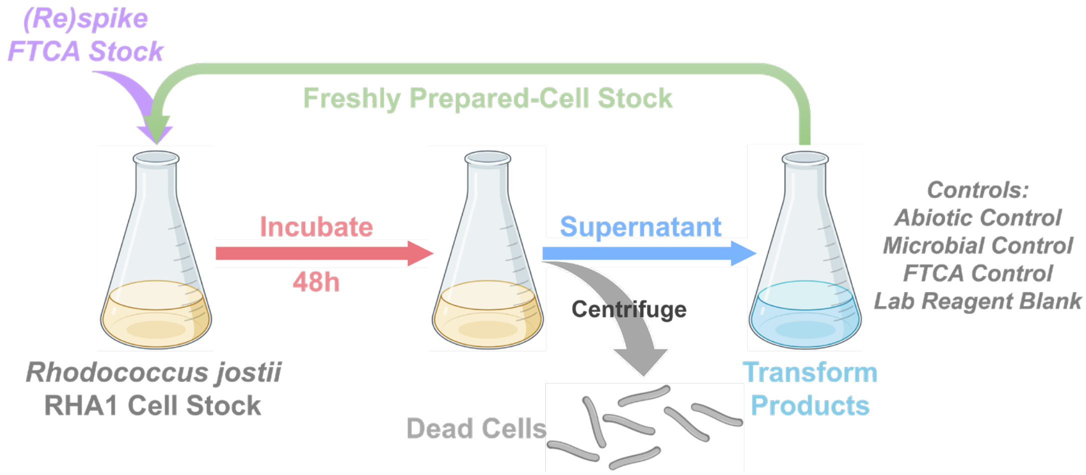


**Figure S2.** Experimental scheme depicting the iterative enrichment strategy for the accumulation of FTCA TPs by RHA1.

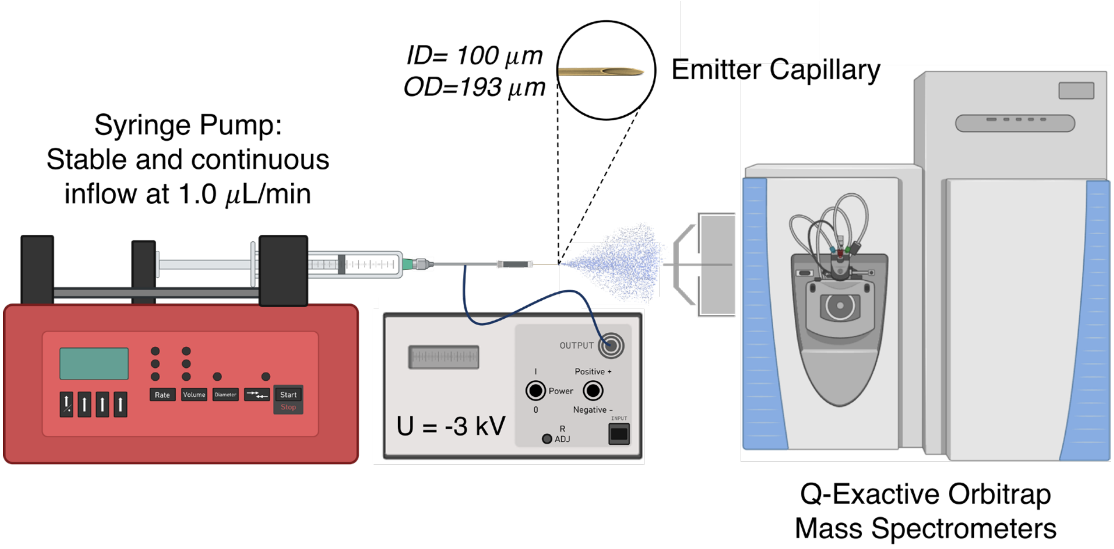


**Figure S3.** Apparatus setup for the Nano-ESI-HRMS analysis.

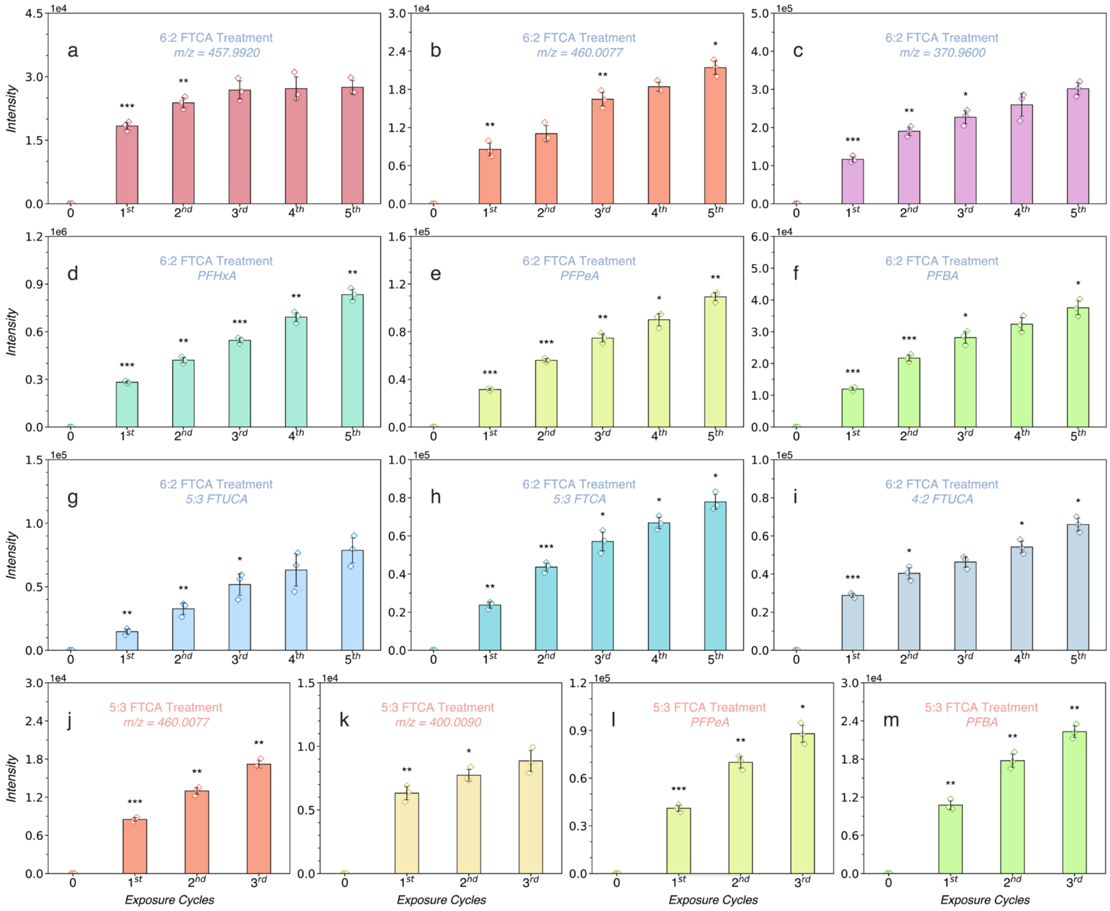


**Figure S4.** Accumulation profiles of additional TPs during iterative enrichment over 48-h exposure cycles to 6:2 and 5:3 FTCA by RHA1.

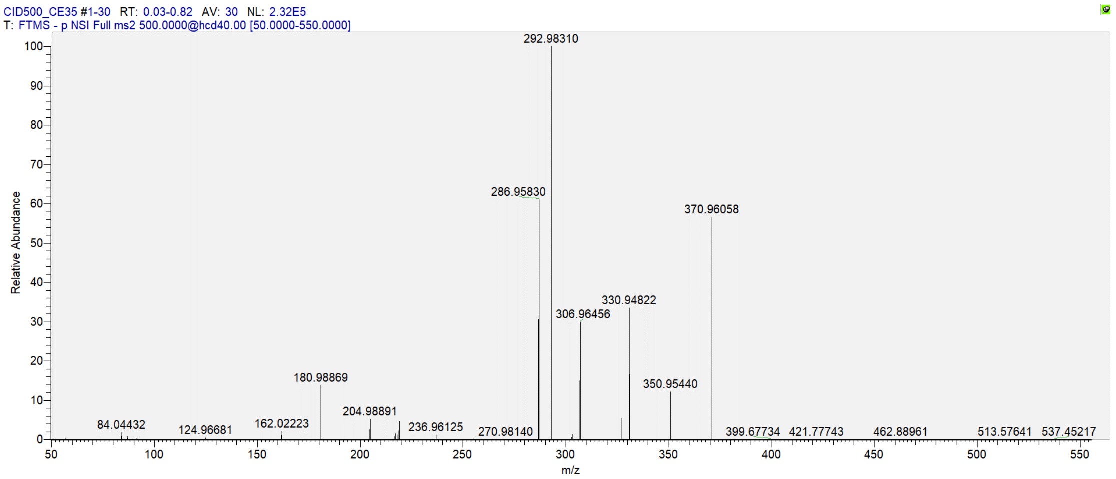


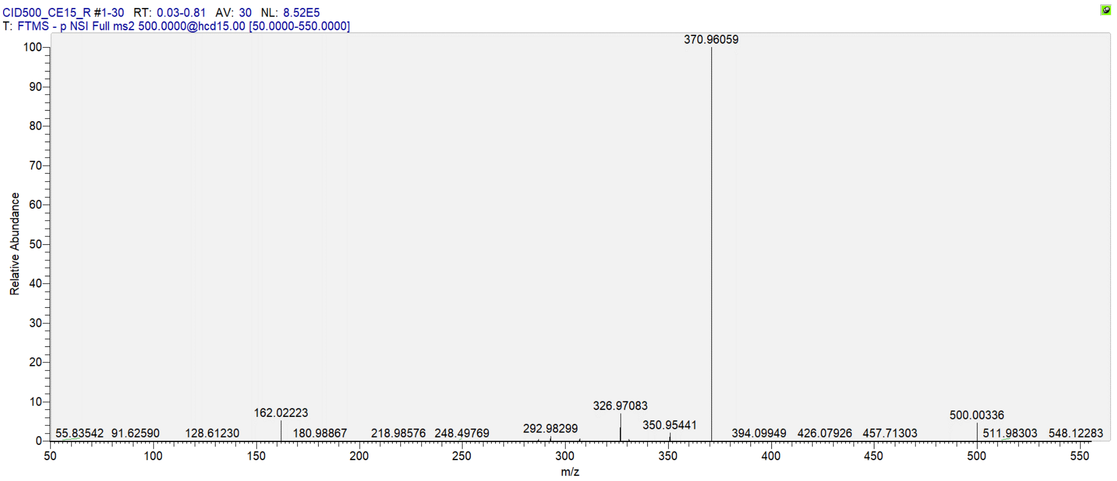


| Product Ions Formula | Theoretical m/z | Observed m/z | Absolute Mass Error (mDa) | Relative Mass Error (ppm) | MS2 Intensity |
|---|---|---|---|---|---|
| $C_2F_5^-$ | 118.99202 | 118.99151 | -0.510 | -4.286 | 349.9 |
| $C_3F_3S^-$ | 124.96728 | 124.96681 | -0.470 | -3.761 | 1153.2 |
| $C_5H_8NO_3S^-$ | 162.02249 | 162.02223 | -0.260 | -1.605 | 4983.2 |
| $C_4F_7^-$ | 180.98882 | 180.98869 | -0.130 | -0.718 | 33389.1 |
| $C_6F_7^-$ | 204.98882 | 204.98891 | 0.090 | 0.439 | 12183.4 |
| $C_7F_7^-$ | 216.98882 | 216.98901 | 0.190 | 0.876 | 3618.0 |
| $C_6F_6S^-$ | 217.96249 | 217.96269 | 0.200 | 0.918 | 3058.2 |
| $C_4F_9^-$ | 218.98563 | 218.98582 | 0.190 | 0.868 | 10859.1 |
| $C_6F_7S^-$ | 236.96089 | 236.96125 | 0.360 | 1.519 | 2962.7 |
| $C_7F_9^-$ | 254.98563 | 254.98612 | 0.490 | 1.922 | 455.4 |
| $C_7F_9O^-$ | 270.98054 | 270.98140 | 0.860 | 3.174 | 364.1 |
| $C_7F_9S^-$ | 286.95770 | 286.95830 | 0.600 | 2.091 | 142884.4 |
| $C_7F_{11}^-$ | 292.98243 | 292.98310 | 0.670 | 2.287 | 233662.9 |
| $C_7F_9OS^-$ | 302.95261 | 302.95322 | 0.610 | 2.014 | 3241.8 |
| $C_7HF_{10}S^-$ | 306.96393 | 306.96456 | 0.630 | 2.052 | 69563.2 |
| $C_7H_2F_{11}S^-$ | 326.97016 | 326.97083 | 0.670 | 2.049 | 12798.6 |

| | | | | | |
|---|---|---|---|---|---|
| $C_8F_9O_2S^-$ | 330.94753 | 330.94822 | 0.690 | 2.085 | 78096.6 |
| $C_8HF_{10}O_2S^-$ | 350.95376 | 350.95440 | 0.640 | 1.824 | 28381.8 |
| $C_8H_2F_{11}O_2S^-$ | 370.95999 | 370.96058 | 0.590 | 1.590 | 132769.8 |
| $C_{13}H_9F_{11}NO_5S^-$ | 500.00258 | 500.00366 | 1.080 | 2.160 | 40772.7 (CE=15V) |

**Figure S5.** MS2 spectrum of AcCyS-5:3 FTUCA and its major product ions generated by CID.

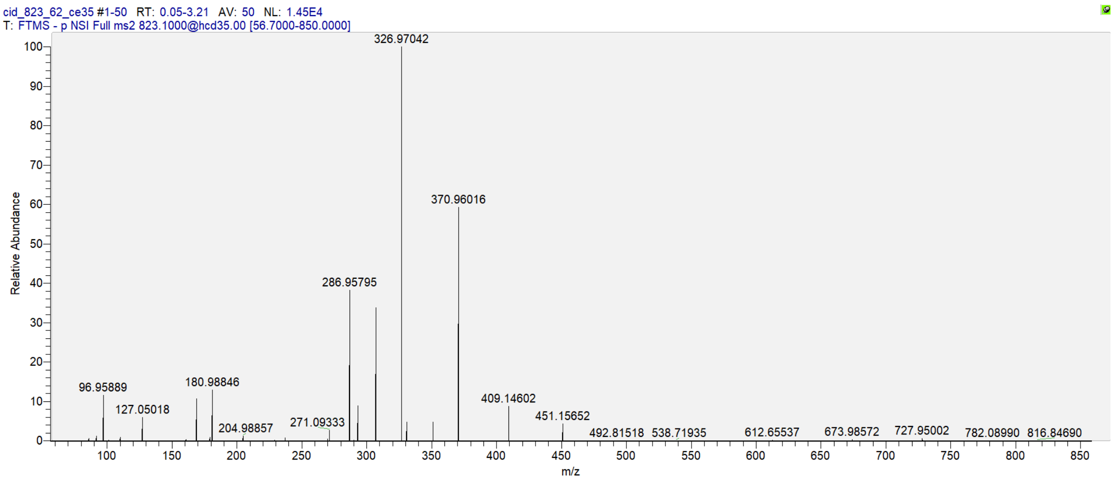


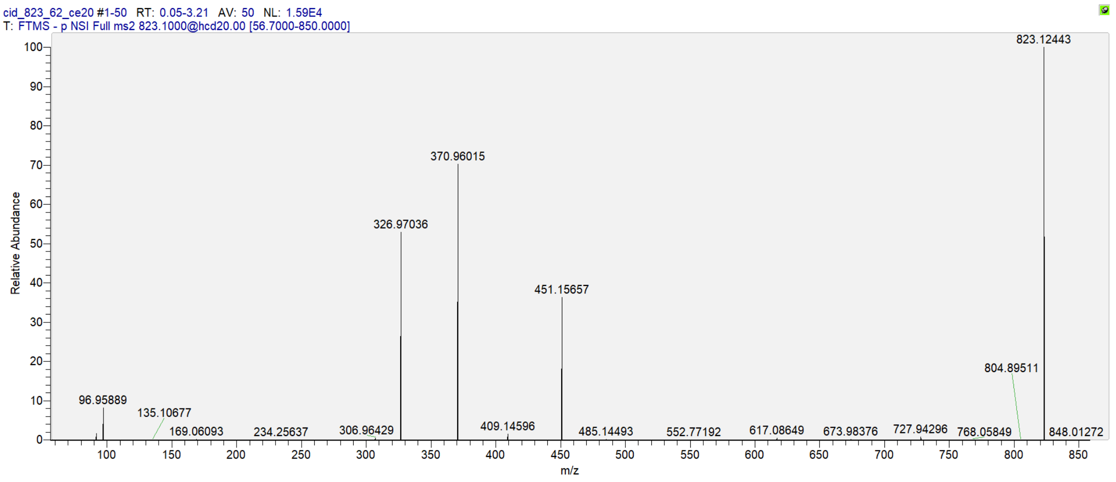


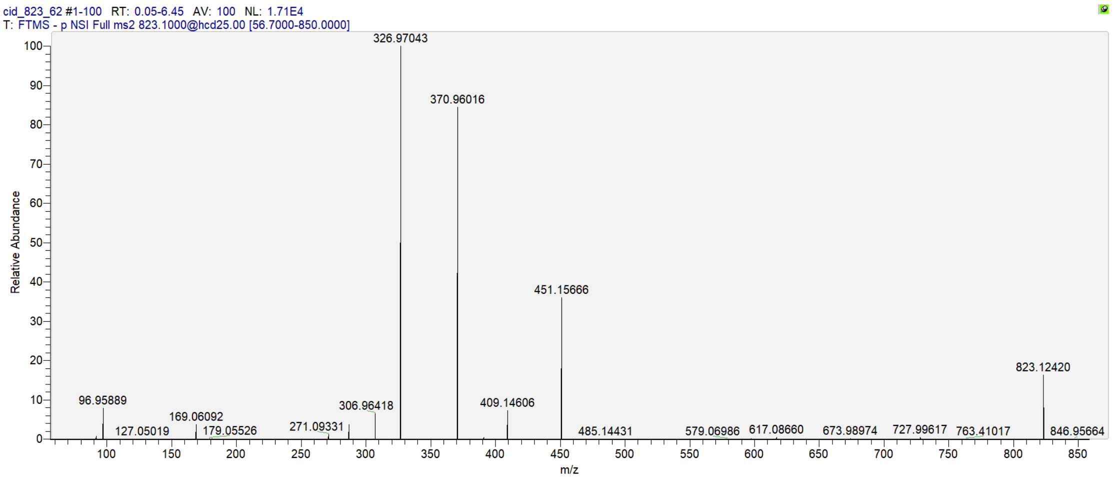


| Product Ions Formula | Theoretical m/z | Observed m/z | Absolute Mass Error (mDa) | Relative Mass Error (ppm) | MS2 Intensity |
|---|---|---|---|---|---|
| $C_5H_7N_2O_2^-$ | 127.05075 | 127.05018 | -0.570 | -4.486 | 874.8 |
| $C_6H_9O_5^-$ | 161.04500 | 161.04450 | -0.500 | -3.105 | 62.8 |
| $C_7H_9N_2O_3^-$ | 169.06132 | 169.06089 | -0.430 | -2.543 | 1590.9 |
| $C_6H_{11}O_6^-$ | 179.05556 | 179.05518 | -0.380 | -2.122 | 141.6 |

| | | | | | |
|---|---|---|---|---|---|
| $C_4F_7^-$ | 180.98882 | 180.98846 | -0.360 | -1.989 | 1905.0 |
| $C_8H_9N_2O_3^-$ | 181.06132 | 181.06098 | -0.340 | -1.878 | 240.5 |
| $C_6F_7^-$ | 204.98882 | 204.98857 | -0.250 | -1.220 | 200.2 |
| $C_9H_{13}N_2O_5^-$ | 229.08245 | 229.08226 | -0.190 | -0.829 | 37.8 |
| $C_6F_7S^-$ | 236.96089 | 236.96096 | 0.070 | 0.295 | 115.6 |
| $C_{11}H_{15}N_2O_6^-$ | 271.09301 | 271.09333 | 0.320 | 1.180 | 394.5 |
| $C_7F_9S^-$ | 286.95770 | 286.95795 | 0.250 | 0.871 | 5666.6 |
| $C_7F_{11}^-$ | 292.98243 | 292.98272 | 0.290 | 0.990 | 1326.7 |
| $C_7HF_{10}S^-$ | 306.96393 | 306.96418 | 0.250 | 0.814 | 1123.1 |
| $C_7H_2F_{11}S^-$ | 326.97016 | 326.97042 | 0.260 | 0.795 | 17302.0 |
| $C_8F_9O_2S^-$ | 330.94753 | 330.94776 | 0.230 | 0.695 | 701.4 |
| $C_8HF_{10}O_2S^-$ | 350.95376 | 350.95399 | 0.230 | 0.655 | 725.3 |
| $C_8H_2F_{11}O_2S^-$ | 370.95999 | 370.96016 | 0.170 | 0.458 | 14866.5 |
| $C_{15}H_{25}N_2O_{11}^-$ | 409.14583 | 409.14602 | 0.190 | 0.464 | 1283.0 |
| $C_{17}H_{27}N_2O_{12}^-$ | 451.15640 | 451.15652 | 0.120 | 0.266 | 639.6 |
| $C_{17}H_{29}N_2O_{12}S^-$ | 485.14412 | 485.14431 | 0.190 | 0.392 | 36.3 (CE=25V) |
| $C_{25}H_{30}F_{11}N_2O_{14}S^-$ | 823.12421 | 823.12420 | -0.01 | -0.012 | 2823.1 (CE=20V) |

**Figure S6.** MS2 spectrum of MS-5:3 FTUCA and its major product ions generated by CID.

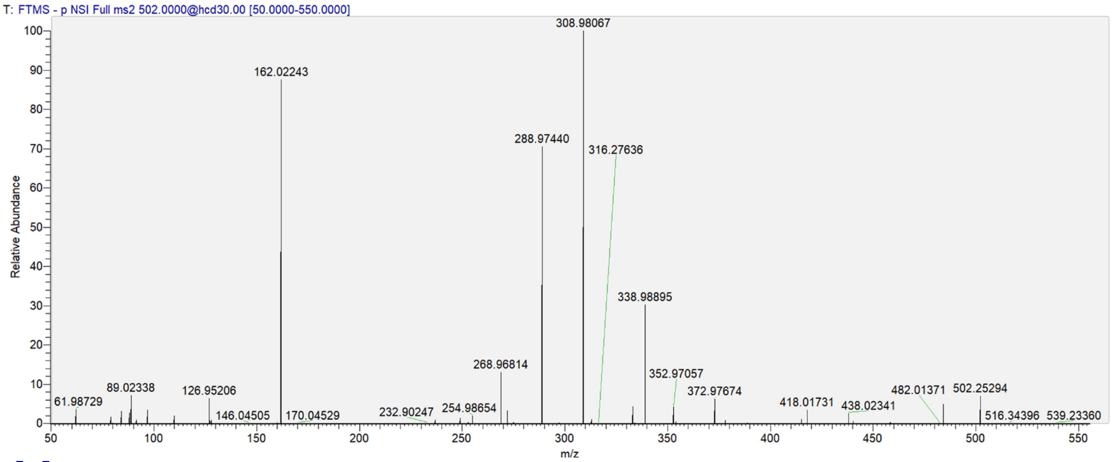


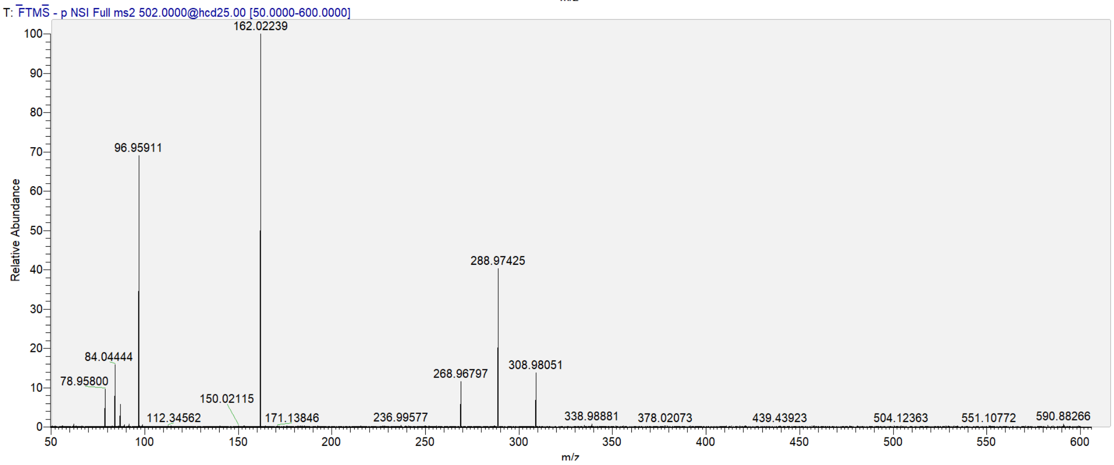


| Product Ions Formula | Theoretical m/z | Observed m/z | Absolute Mass Error (mDa) | Relative Mass Error (ppm) | MS2 Intensity |
|---|---|---|---|---|---|
| $C_4H_6NO^-$ | 84.04494 | 84.04443 | -0.510 | -6.068 | 719.6 |
| $C_3H_6NO_2^-$ | 88.03985 | 88.03935 | -0.500 | -5.679 | 627.6 |
| $C_5H_8NO_3S^-$ | 162.02249 | 162.02243 | -0.060 | -0.370 | 20031.2 |
| $C_7F_7^-$ | 216.98882 | 216.98937 | 0.550 | 2.535 | 119.0 |
| $C_7HF_8^-$ | 236.99505 | 236.99575 | 0.700 | 2.954 | 216.9 |
| $C_7F_7S^-$ | 248.96089 | 248.96167 | 0.780 | 3.133 | 309.5 |
| $C_7F_9^-$ | 254.98563 | 254.98654 | 0.910 | 3.569 | 441.7 |
| $C_7HF_8S^-$ | 268.96712 | 268.96814 | 1.020 | 3.792 | 3011.7 |
| $C_5F_{11}^-$ | 268.98243 | 268.98355 | 1.120 | 4.164 | 130.8 |
| $C_7HF_{10}^-$ | 274.99186 | 274.99300 | 1.140 | 4.146 | 97.9 |
| $C_7H_2F_9S^-$ | 288.97335 | 288.97440 | 1.050 | 3.634 | 16136.2 |
| $C_7H_3F_{10}S^-$ | 308.97958 | 308.98067 | 1.090 | 3.528 | 22849.1 |
| $C_8HF_8O_2S^-$ | 312.95695 | 312.95815 | 1.200 | 3.834 | 257.9 |
| $C_8H_2F_9O_2S^-$ | 332.96318 | 332.96437 | 1.190 | 3.574 | 982.7 |
| $C_8H_2F_{11}O_2^-$ | 338.98791 | 338.98895 | 1.040 | 3.068 | 6914.0 |
| $C_8H_3F_{10}O_2S^-$ | 352.96941 | 352.97057 | 1.160 | 3.286 | 978.8 |
| $C_8H_4F_{11}O_2S^-$ | 372.97564 | 372.97674 | 1.100 | 2.949 | 1451.5 |
| $C_{12}H_7F_7NO_3S^-$ | 378.00349 | 378.00471 | 1.220 | 3.227 | 192.2 |

| | | | | | |
|---|---|---|---|---|---|
| $C_{12}H_9F_9NO_3S^-$ | 418.01594 | 418.01731 | 1.370 | 3.277 | 805.3 |
| $C_{12}H_{10}F_{10}NO_3S^-$ | 438.02217 | 438.02341 | 1.240 | 2.831 | 583.1 |
| $C_{13}H_{11}F_{11}NO_5S^-$ | 502.01823 | 502.01981 | 1.580 | 3.147 | 1290.4 (CE=20V) |

**Figure S7.** MS2 spectrum of AcCyS-5:3 FTCA and its major product ions generated by CID.

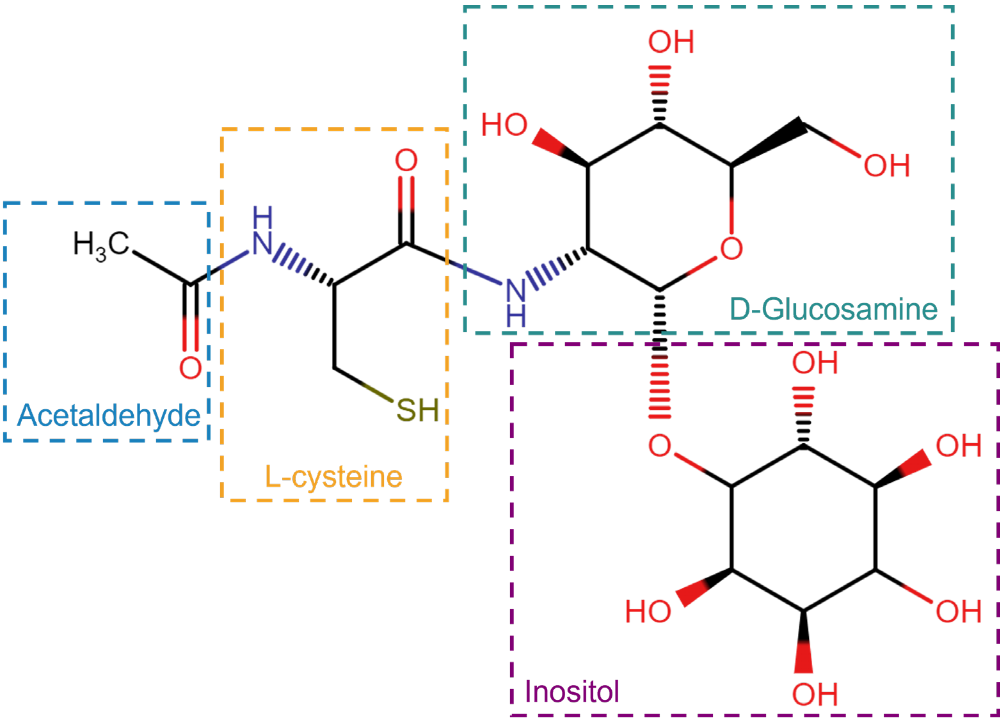


**Figure S8.** Molecular structure of mycothiol (Ac-CyS-GlcN-Ins), highlighting its four moieties that include acetyl group, cysteine, glucosamine, and *myo*-inositol.

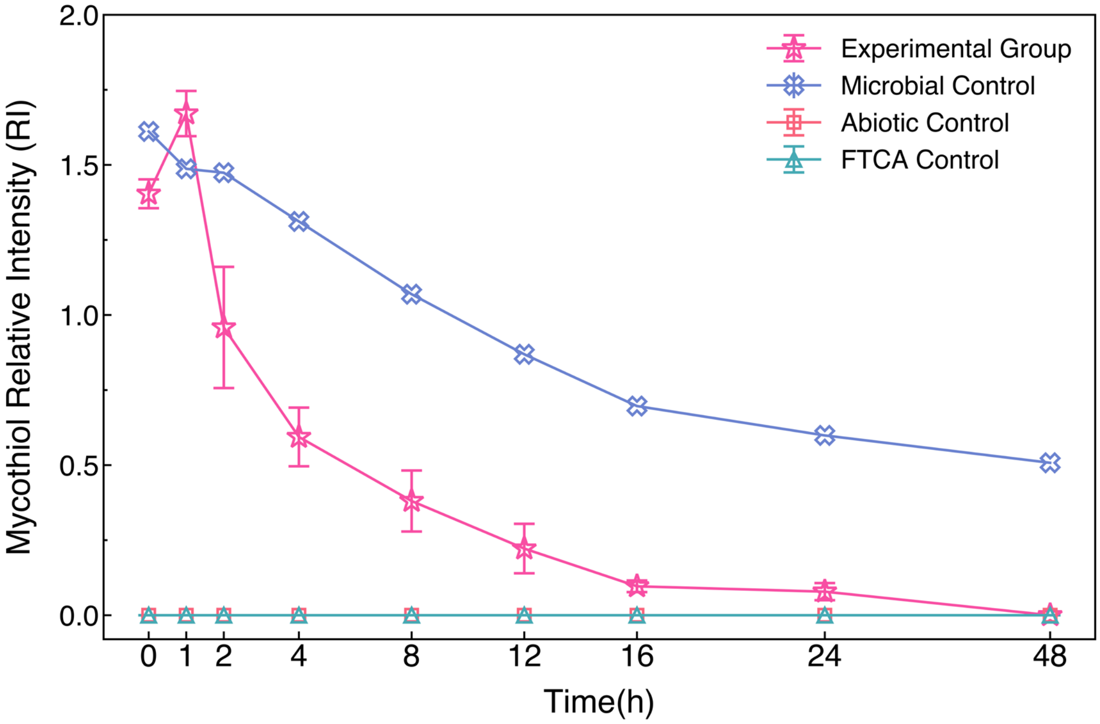


**Figure S9.** MSH concentrations during 6:2 FTCA treatments, as compared to microbial controls, abiotic controls, and analytical controls.

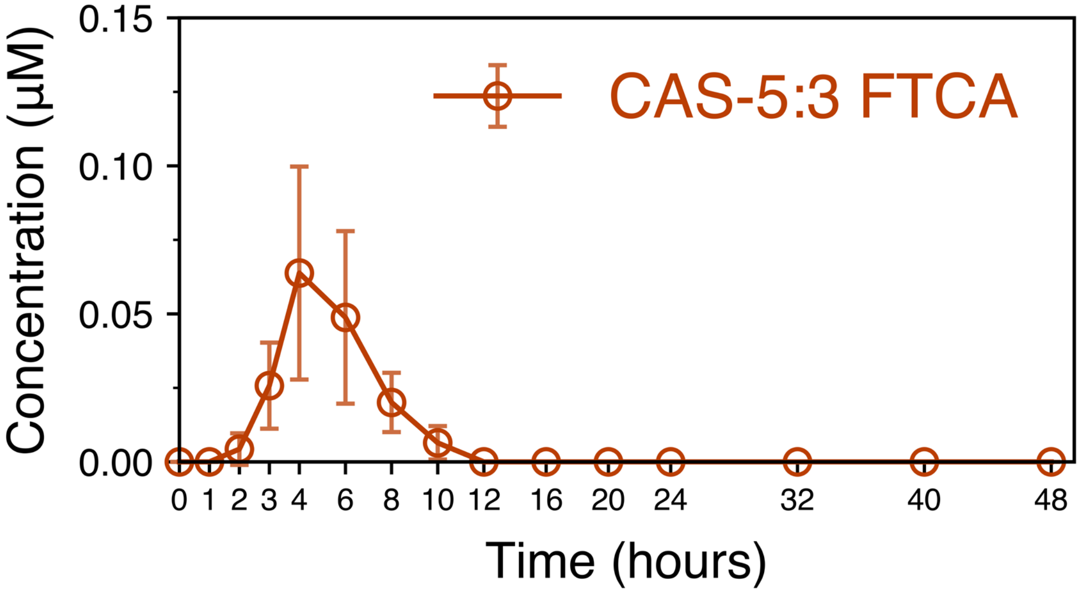


**Figure S10**. Transient accumulation of the cysteamine conjugate CAS-5:3 FTCA in 6:2 FTCA treatment by RHA1. The concentration profile of CAS-5:3 FTCA is similar to CAS-5:3 FTUCA, as short-lived intermediate in the downstream MSH-mediated conjugation pathway.

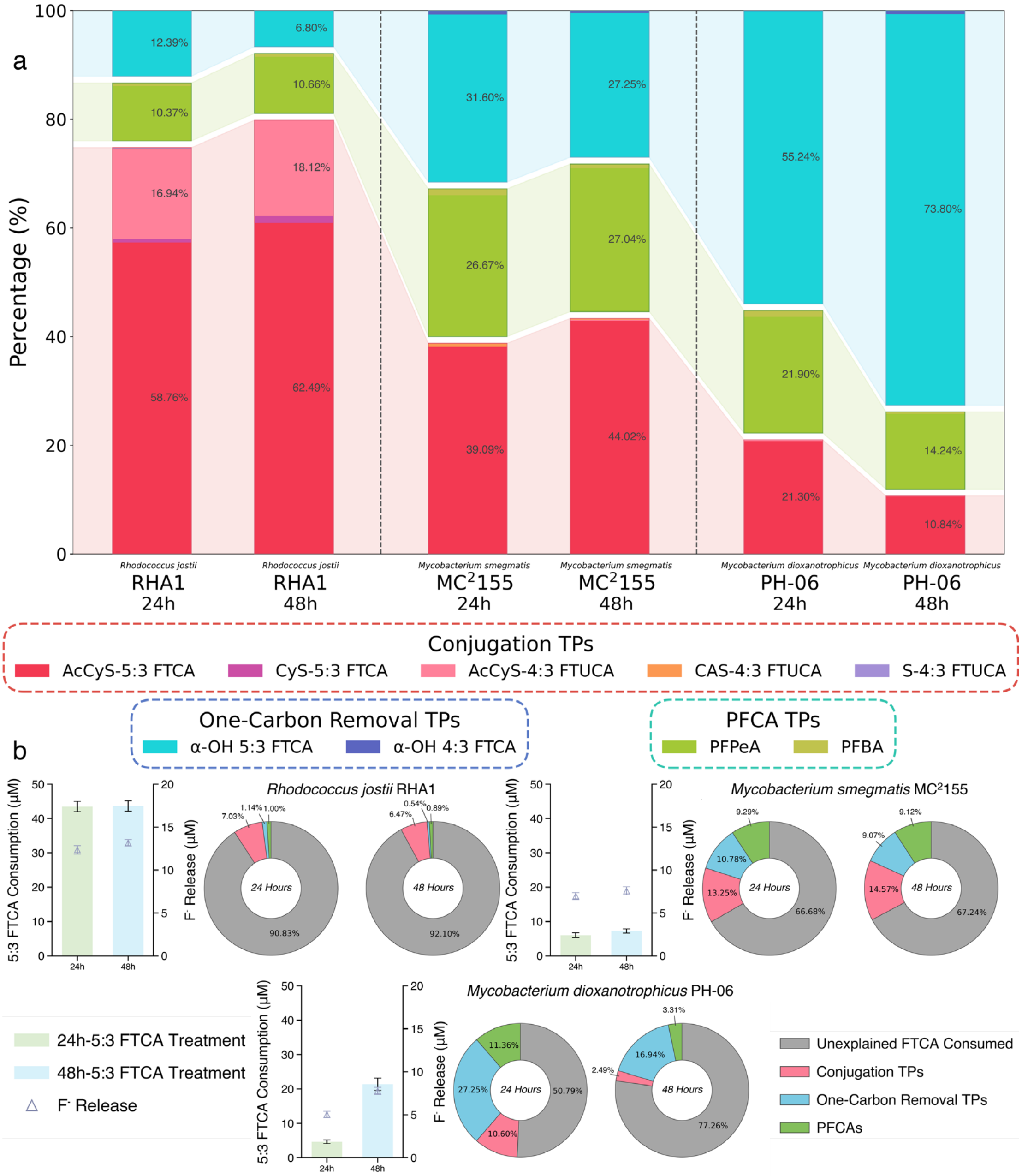


**Figure S11.** Comparison of 5:3 FTCA biotransformation in RHA1, MC$^2$155, and PH-06. (a) TP profiling after 24-h and 48-h exposure to 5:3 FTCA. (b) 5:3 FTCA removal, fluoride release, and the mass discrepancies unaccountable by MSH-mediated conjugation and "one-carbon removal pathways"-derived intermediates or PFCAs.

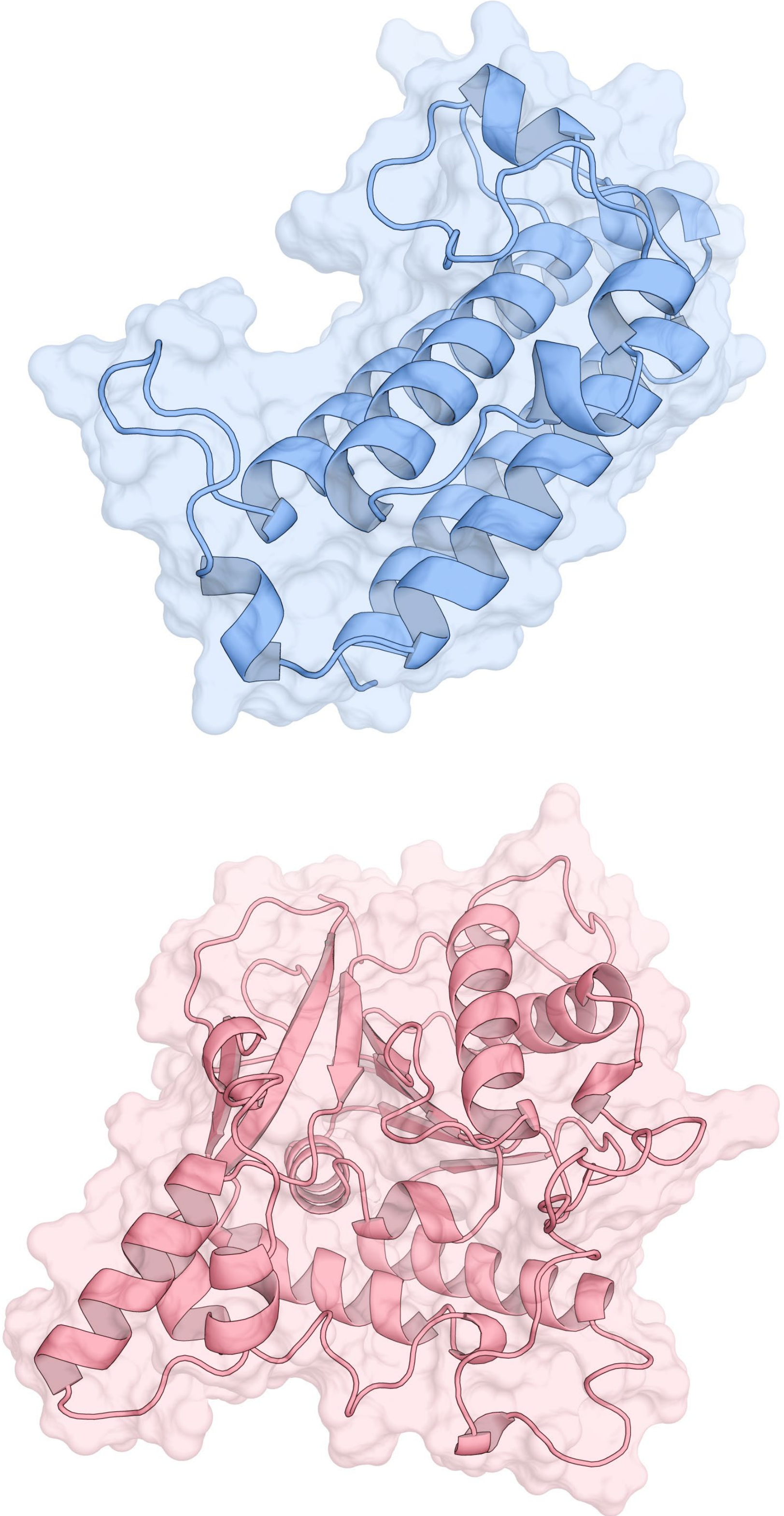

**Figure S12.** Predicted 3-D structures of *R.jostii* RHA1 MST (top) and Mca (bottom) computed by AlphaFold2.

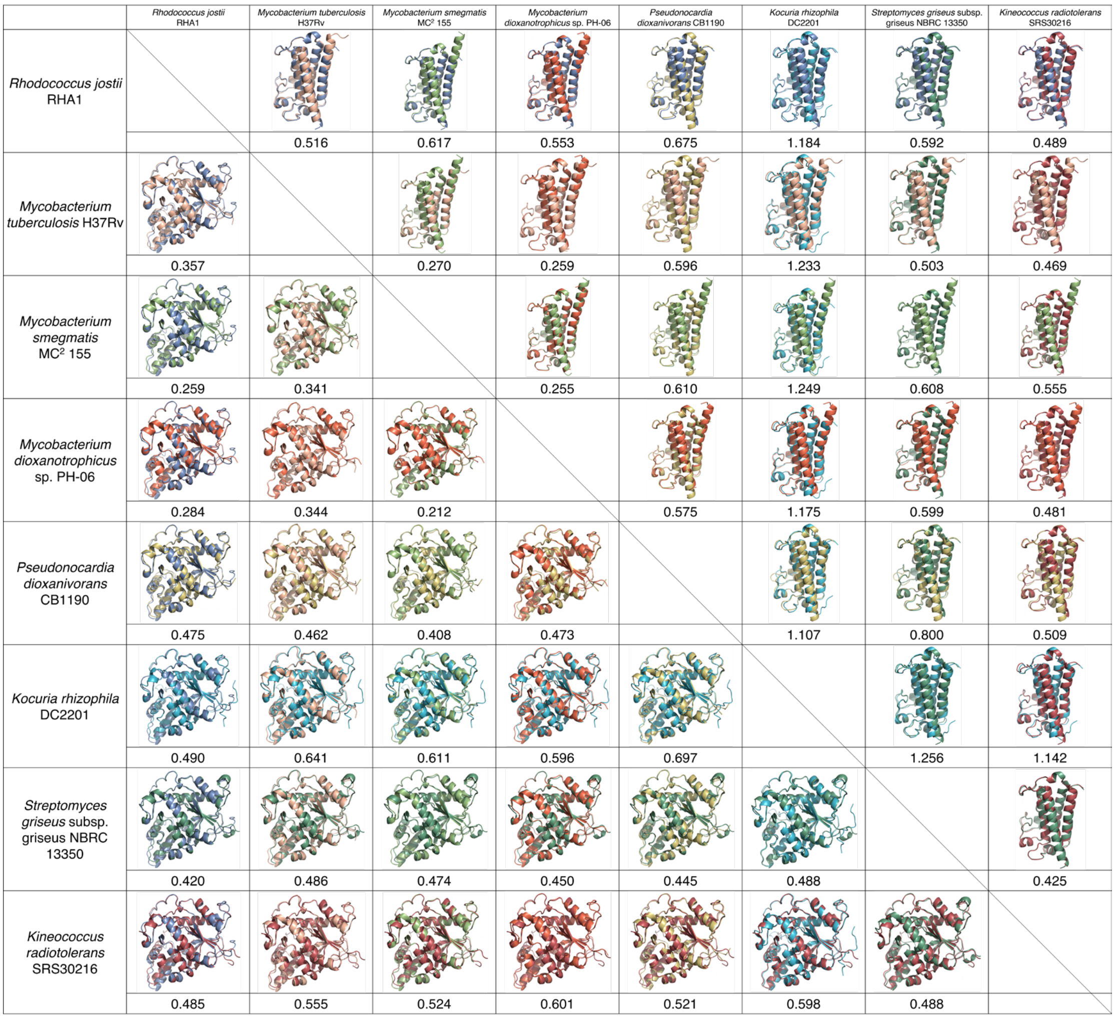


**Figure S13.** Structural conservation of the MSH-conjugation enzymes, MST (top right) and Mca (bottom left). Pairwise structural similarity among eight representative *Actinomycetota* species with the RMSD value of each alignment.

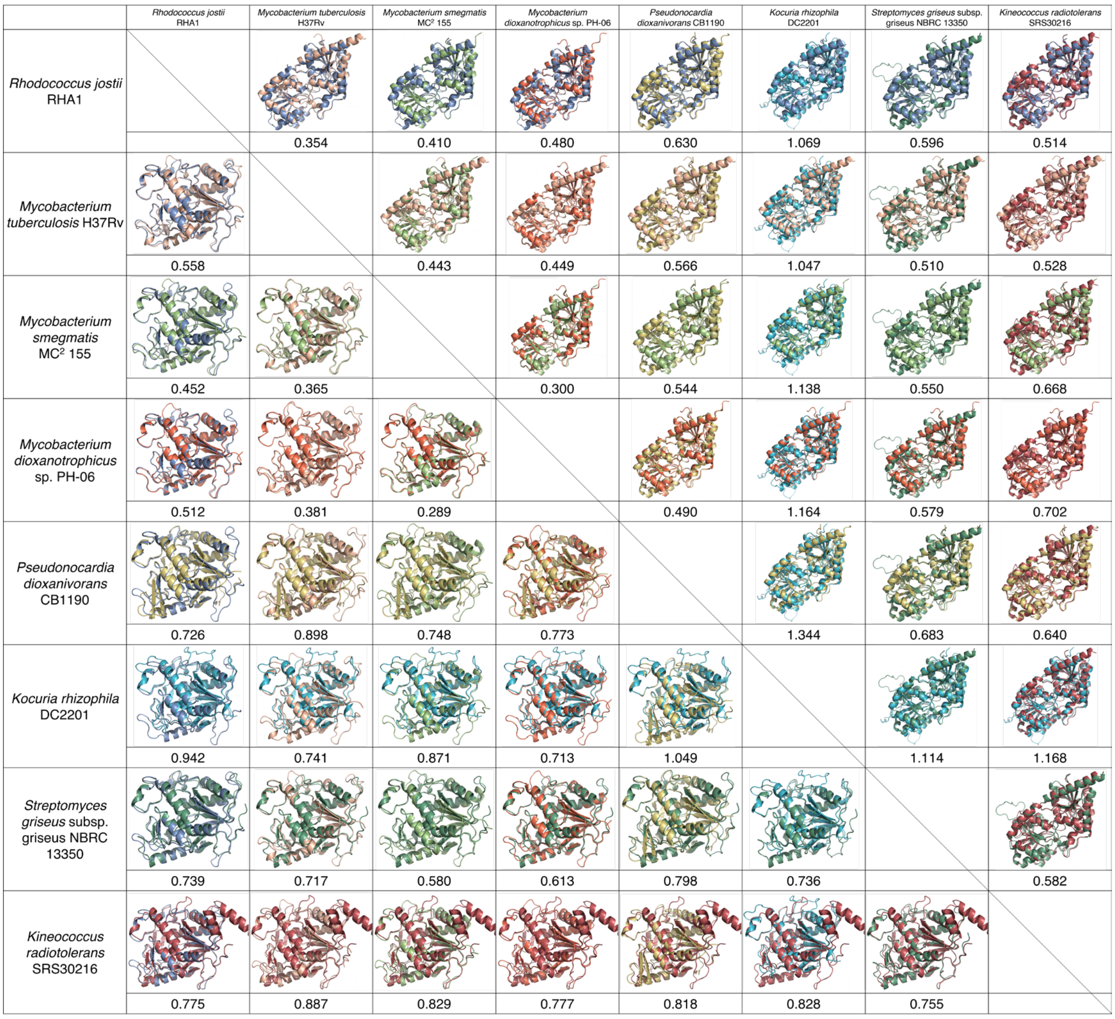


**Figure S14.** Structural conservation of the MSH- biosynthetic enzymes, MshA (top right) and MshB (bottom left). Pairwise structural similarity among eight representative *Actinomycetota* species with the RMSD value of each alignment.

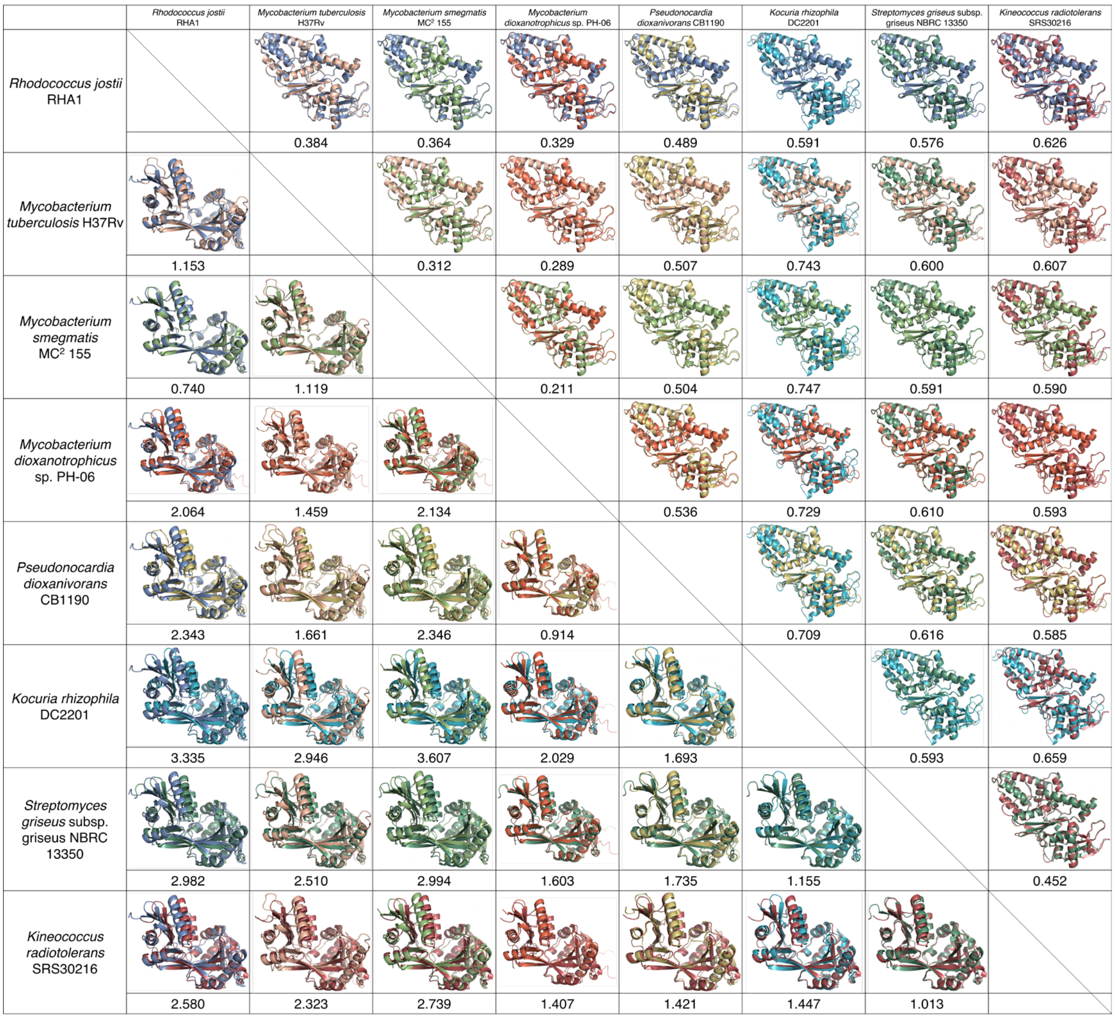


**Figure S15.** Structural conservation of the MSH- biosynthetic enzymes, MshC (top right) and MshD (bottom left). Pairwise structural similarity among eight representative *Actinomycetota* species with the RMSD value of each alignment.

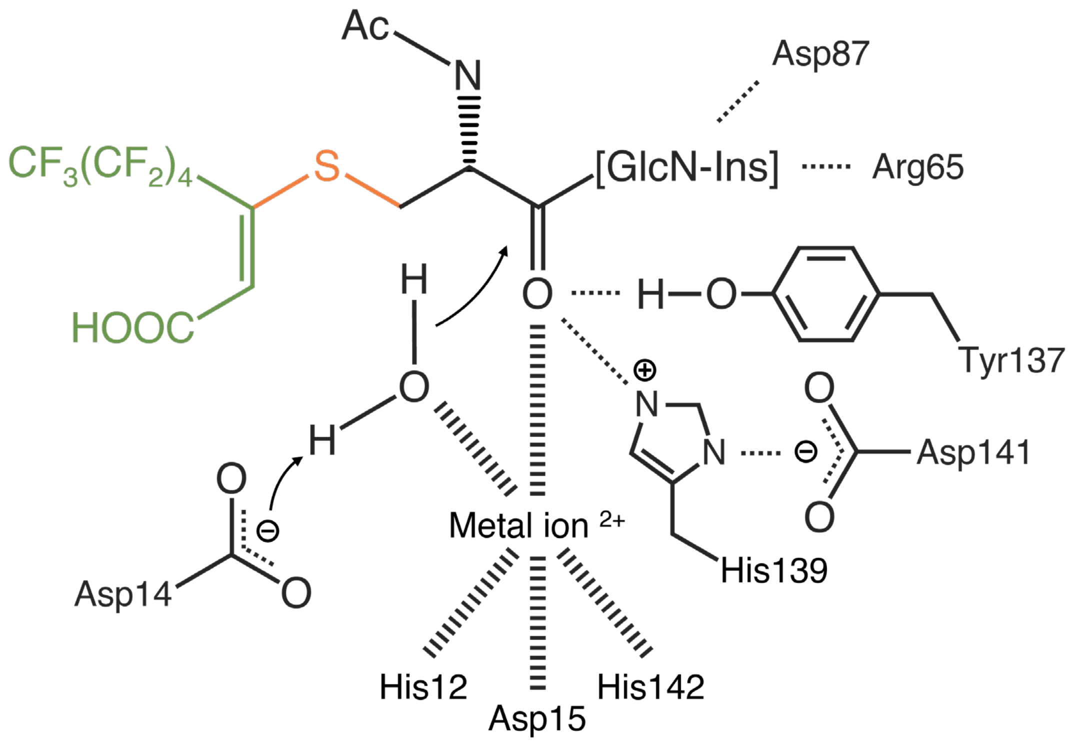


**Figure S16.** Predicted docking of MS-5:3 FTUCA (ligand) in the active site of Mca, highlighting key residues involved in hydrogen bonding and metal coordination.

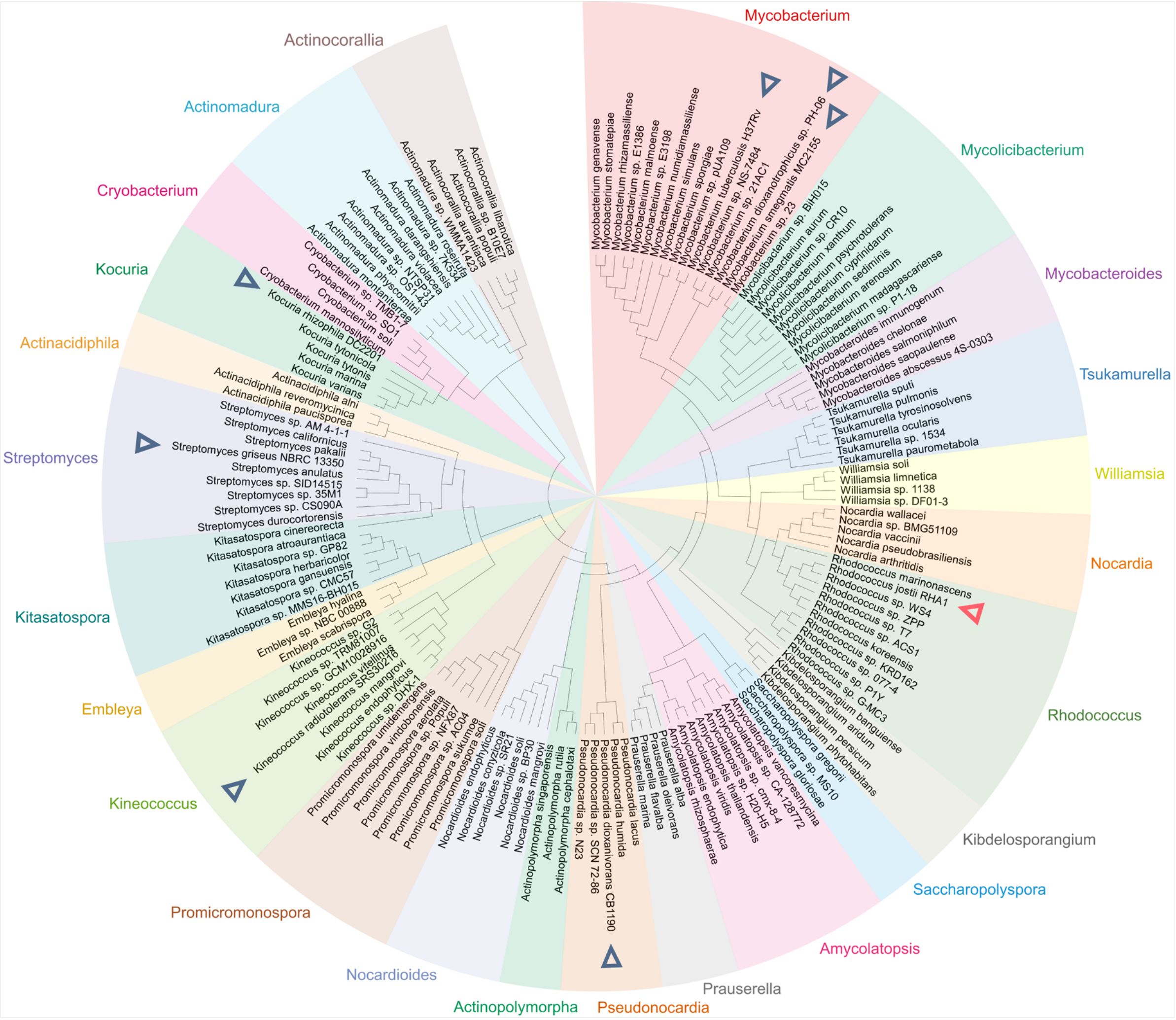


**Figure S17.** Phylogenetic tree (maximum-likelihood) depicting the evolutionary relationships among 150 Mca proteins from 24 genera in the phylum of *Actinomycetota* (equivalent to species shown in the MST phylogenetic tree in Figure 7c).

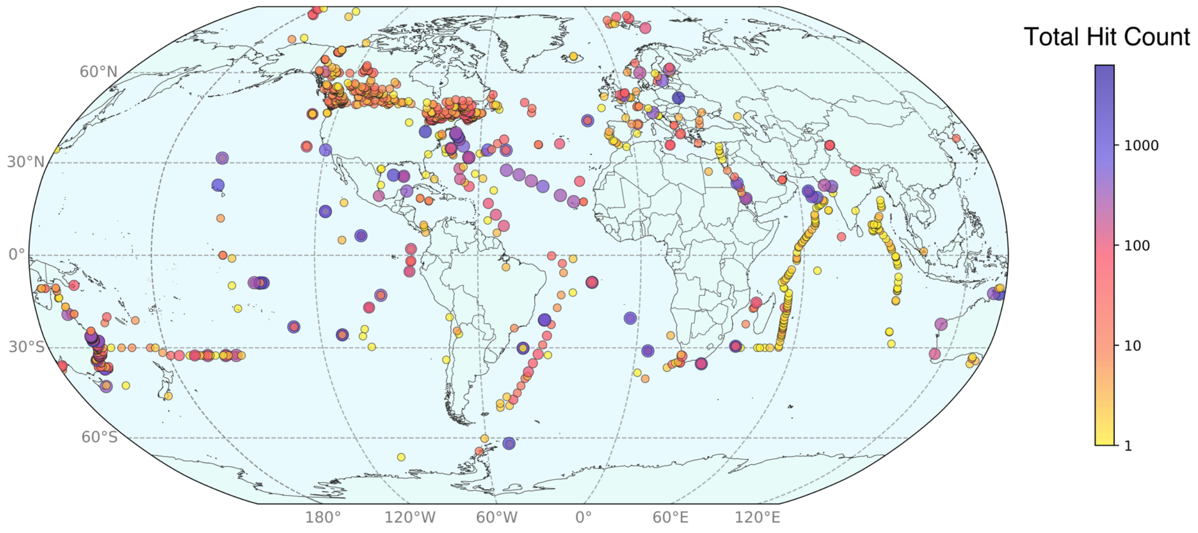


**Figure S18.** Widespread occurrence of MST homologs across the global aquatic metagenomes (marine and freshwater). Each circle represents the location where MST homologs were detected, and circle color corresponds to the number of identified homolog positive hits.

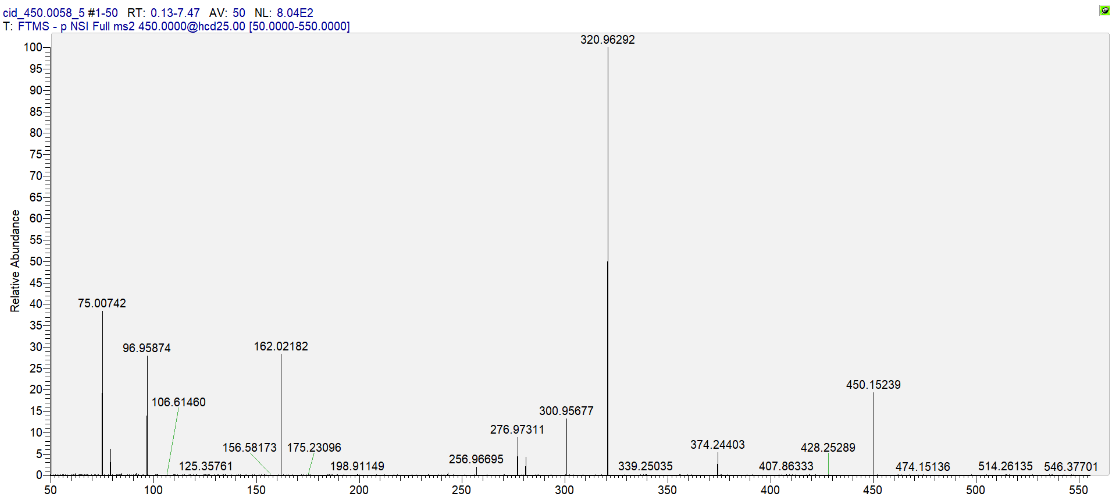


| Product Ions Formula | Theoretical m/z | Observed m/z | Absolute Mass Error (mDa) | Relative Mass Error (ppm) | MS2 Intensity |
|---|---|---|---|---|---|
| $C_5H_8NO_3S^-$ | 162.02249 | 162.02182 | -0.670 | -4.135 | 231.1 |
| $C_6F_7^-$ | 204.98882 | 204.98792 | -0.900 | -4.390 | 8.9 |
| $C_6F_7S^-$ | 236.96089 | 236.96094 | 0.050 | 0.211 | 11.0 |
| $C_6F_9^-$ | 242.98563 | 242.98553 | -0.100 | -0.412 | 15.1 |
| $C_6HF_8S^-$ | 256.96712 | 256.96695 | -0.170 | -0.662 | 26.3 |
| $C_6H_2F_9S^-$ | 276.97335 | 276.97311 | -0.240 | -0.867 | 72.6 |
| $C_7F_7O_2S^-$ | 280.95072 | 280.95047 | -0.250 | -0.890 | 35.6 |
| $C_7HF_8O_2S^-$ | 300.95695 | 300.95677 | -0.180 | -0.598 | 107.8 |
| $C_7H_2F_9O_2S^-$ | 320.96318 | 320.96292 | -0.260 | -0.810 | 813.7 |
| $C_{12}H_9F_9NO_5S^-$ | 450.00577 | 450.00547 MS1 | -0.300 | -0.667 | - |

**Figure S19.** MS2 spectrum of AcCyS-4:3 FTUCA and its major product ions generated by CID.

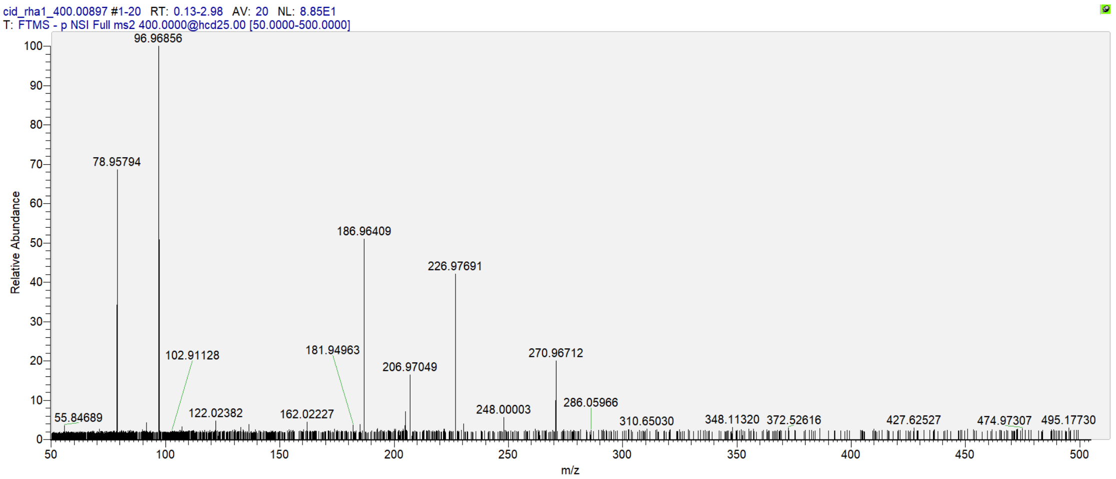


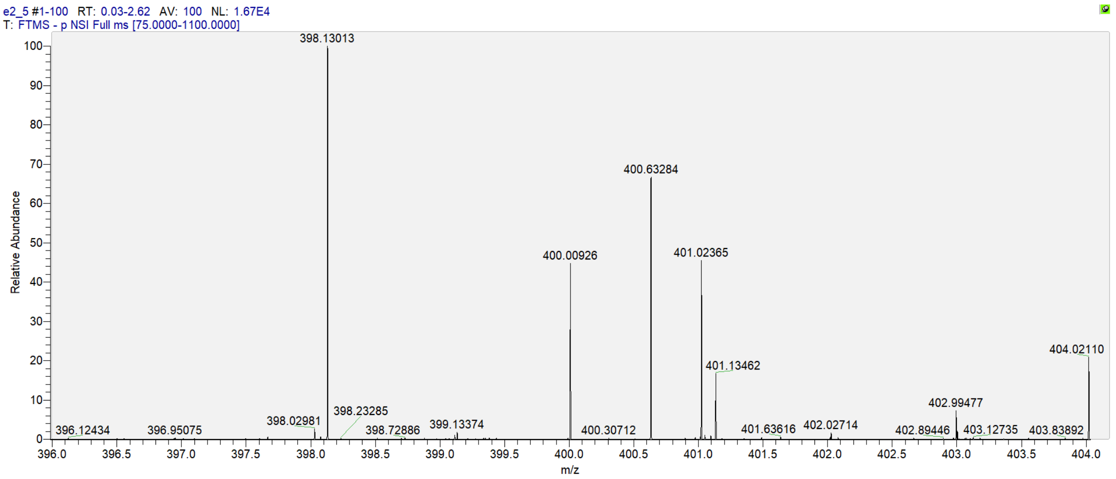


| Product Ions Formula | Theoretical m/z | Observed m/z | Absolute Mass Error (mDa) | Relative Mass Error (ppm) | MS2 Intensity |
|---|---|---|---|---|---|
| $C_5H_8NO_3S^-$ | 162.02249 | 162.02227 | -0.220 | -1.358 | 12.1 |
| $C_5F_5S^-$ | 186.96409 | 186.96409 | 0.000 | 0.000 | 135.5 |
| $C_5HF_6S^-$ | 206.97032 | 206.97049 | 0.170 | 0.821 | 44.0 |
| $C_5H_2F_7S^-$ | 226.97654 | 226.97691 | 0.370 | 1.630 | 112.1 |
| $C_6H_2F_7O_2S^-$ | 270.96637 | 270.96712 | 0.750 | 2.768 | 53.4 |
| $C_{11}H_9F_7NO_5S^-$ | 400.00897 | 400.00926 | 0.290 | 0.725 | MS1 |

**Figure S20.** MS2 spectrum of AcCyS-3:3 FTUCA and its major product ions generated by CID.

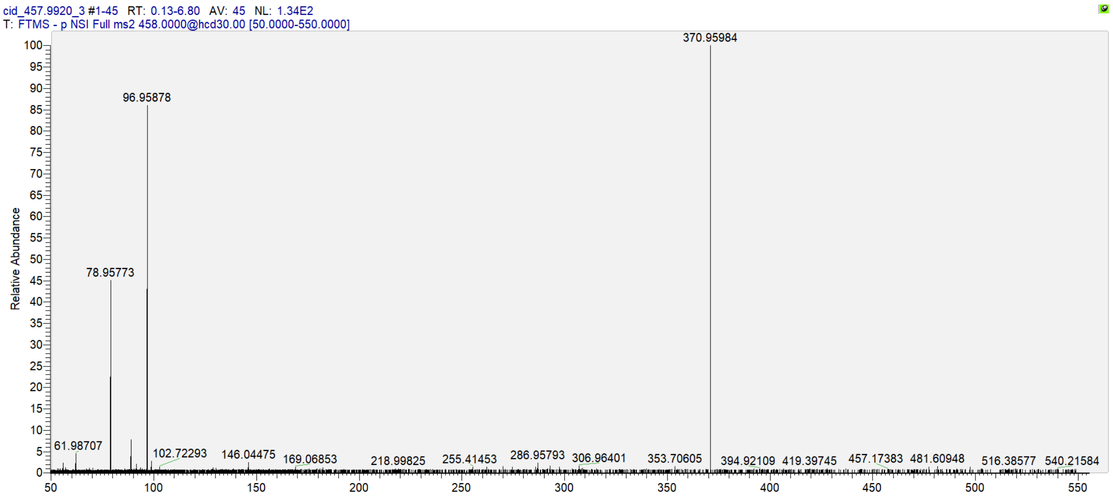


| Product Ions Formula | Theoretical m/z | Observed m/z | Absolute Mass Error (mDa) | Relative Mass Error (ppm) | MS2 Intensity |
|---|---|---|---|---|---|
| $C_7F_9S^-$ | 286.95770 | 286.95793 | 0.230 | 0.802 | 8.2 |
| $C_7F_{11}^-$ | 292.98243 | 292.98225 | -0.180 | -0.614 | 7.3 |
| $C_7HF_{10}S^-$ | 306.96393 | 306.96401 | 0.080 | 0.261 | 7.2 |
| $C_8H_2F_{11}O_2S^-$ | 370.95999 | 370.95984 | -0.150 | -0.404 | 138.3 |
| $C_{11}H_7F_{11}NO_4S^-$ | 457.99201 | 457.99180 MS1 | -0.210 | -0.459 | - |

**Figure S21.** MS2 spectrum of CyS-5:3 FTUCA and its major product ions generated by CID.

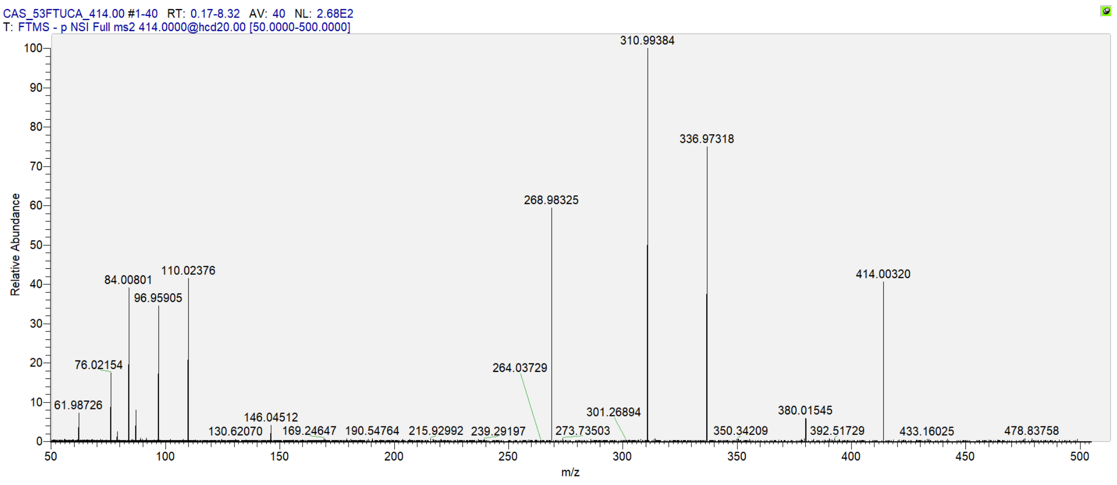


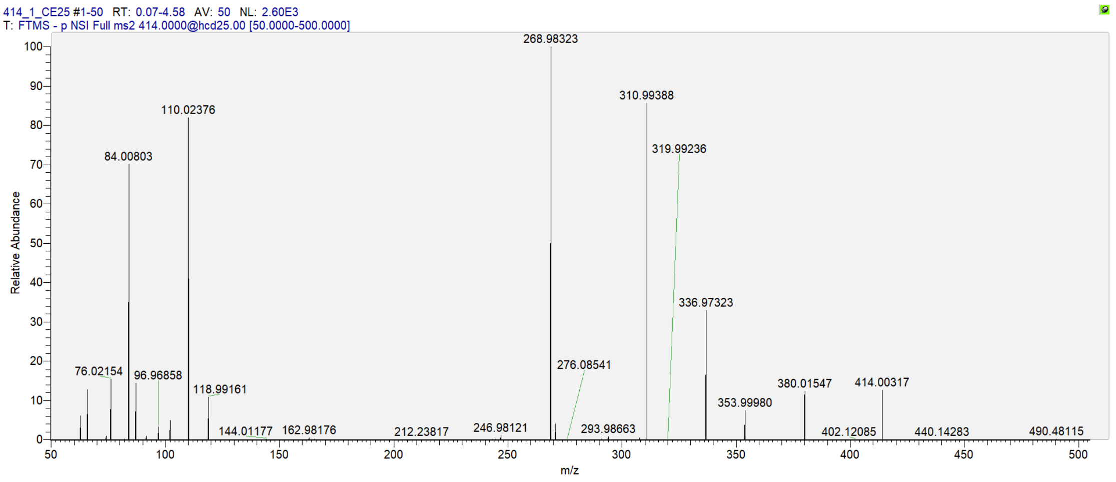


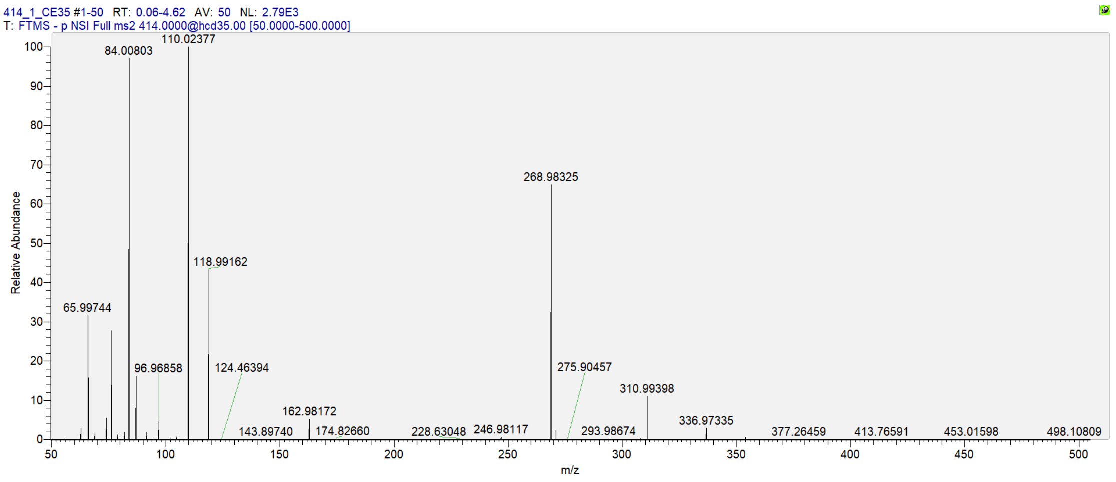


| Product Ions Formula | Theoretical m/z | Observed m/z | Absolute Mass Error (mDa) | Relative Mass Error (ppm) | MS2 Intensity |
|---|---|---|---|---|---|
| $C_2H_4NS^-$ | 74.00645 | 74.00588 | -0.570 | -7.702 | 26.6 |
| $C_2H_6NS^-$ | 76.02210 | 76.02154 | -0.560 | -7.366 | 407.9 |

| | | | | | |
|---|---|---|---|---|---|
| $C_3FO_2^-$ | 86.98823 | 86.98771 | -0.520 | -5.978 | 389.4 |
| $C_2F_5^-$ | 118.99202 | 118.99161 | -0.410 | -3.446 | 289.2 |
| $C_5F_9O^-$ | 246.98054 | 246.98121 | 0.670 | 2.713 | 29.0 |
| $C_5F_{11}^-$ | 268.98243 | 268.98323 | 0.800 | 2.974 | 2609.8 |
| $C_7F_9O^-$ | 270.98054 | 270.98133 | 0.790 | 2.915 | 109.1 |
| $C_8F_{11}O_2^-$ | 336.97226 | 336.97323 | 0.970 | 2.879 | 873.7 |
| $C_{10}H_7F_{11}NO_2S^-$ | 414.00218 | 414.00317 | 0.990 | 2.391 | 338.2 |

**Figure S22.** MS2 spectrum of CAS-5:3 FTUCA and its major product ions generated by CID.

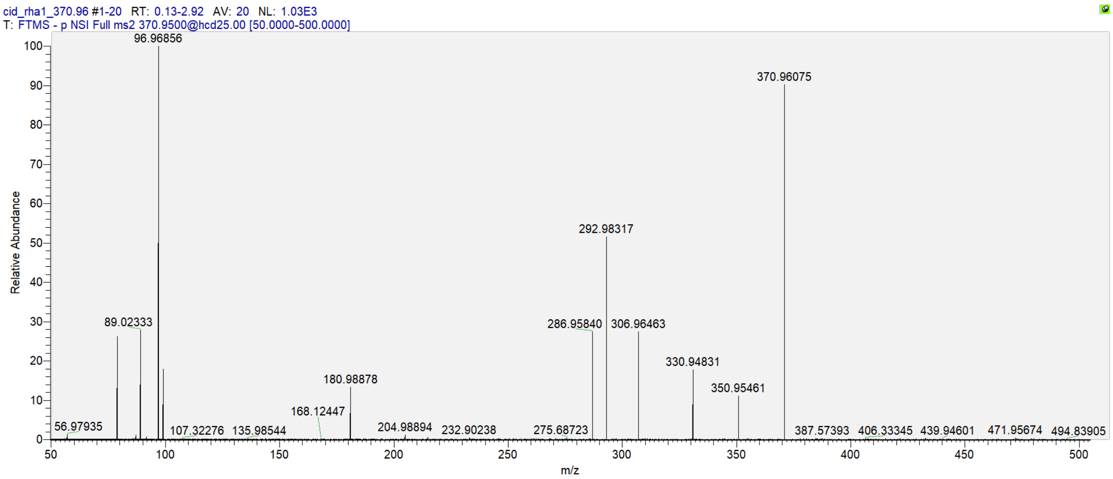


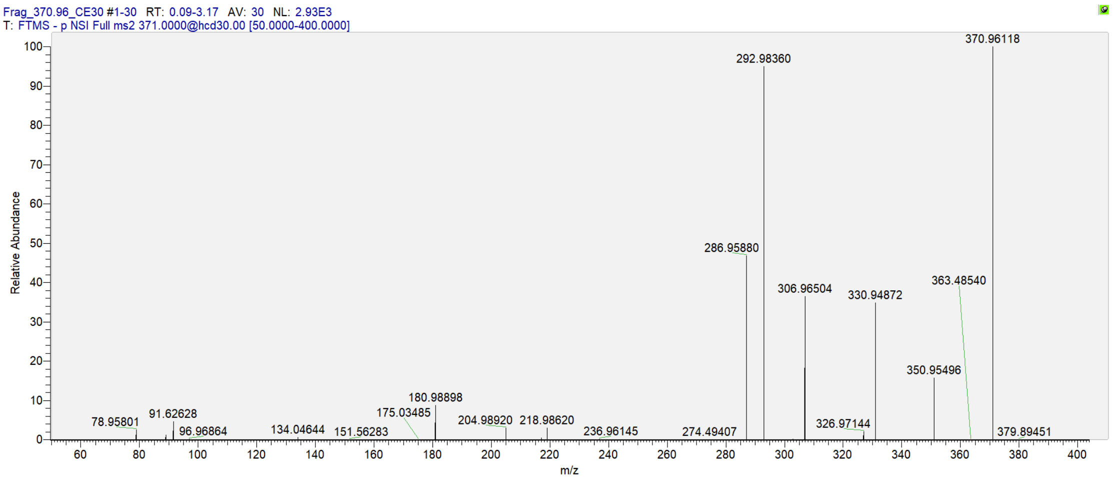


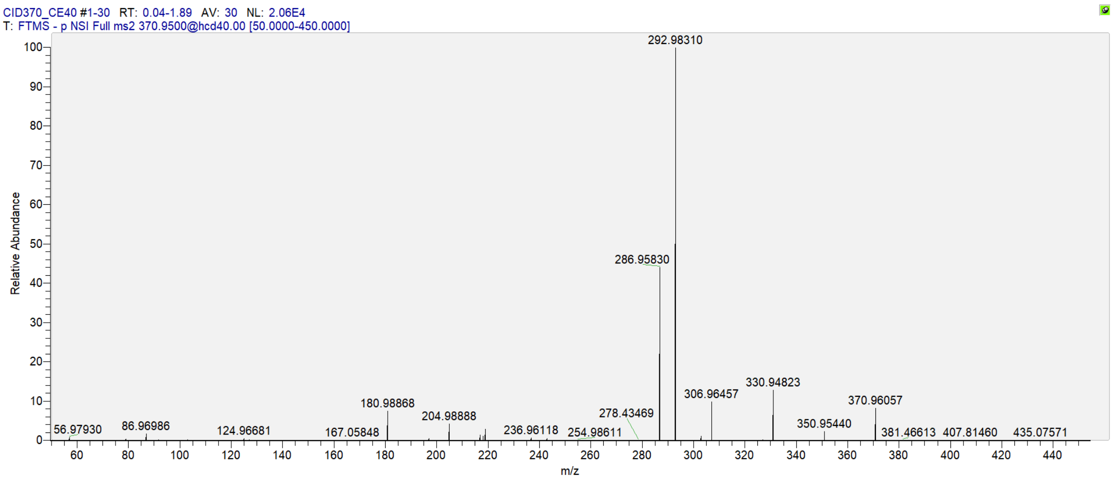


| Product Ions Formula | Theoretical m/z | Observed m/z | Absolute Mass Error (mDa) | Relative Mass Error (ppm) | MS2 Intensity |
|---|---|---|---|---|---|
| $C_2HS^-$ | 56.97990 | 56.97930 | -0.600 | -10.530 | 182.2 |
| $CF_3^-$ | 68.99521 | 68.99454 | -0.670 | -9.711 | 9.8 |

| | | | | | |
|---|---|---|---|---|---|
| $C_3FS^-$ | 86.97047 | 86.96986 | -0.610 | -7.014 | 346.0 |
| $C_3FOS^-$ | 102.96539 | 102.96481 | -0.580 | -5.633 | 40.8 |
| $C_2F_5^-$ | 118.99202 | 118.99150 | -0.520 | -4.370 | 12.5 |
| $C_3F_3S^-$ | 124.96728 | 124.96681 | -0.470 | -3.761 | 123.9 |
| $C_4F_7^-$ | 180.98882 | 180.98868 | -0.140 | -0.774 | 1568.9 |
| $C_4F_7O^-$ | 196.98374 | 196.98372 | -0.020 | -0.102 | 119.6 |
| $C_6F_7^-$ | 204.98882 | 204.98888 | 0.060 | 0.293 | 902.1 |
| $C_7F_7^-$ | 216.98882 | 216.98896 | 0.140 | 0.645 | 299.0 |
| $C_6F_6S^-$ | 217.96249 | 217.96269 | 0.200 | 0.918 | 234.2 |
| $C_4F_9^-$ | 218.98563 | 218.98581 | 0.180 | 0.822 | 609.7 |
| $C_6F_7S^-$ | 236.96089 | 236.96118 | 0.290 | 1.224 | 135.1 |
| $C_6F_9^-$ | 242.98563 | 242.98602 | 0.390 | 1.605 | 115.0 |
| $C_7F_9^-$ | 254.98563 | 254.98611 | 0.480 | 1.882 | 45.2 |
| $C_7F_9O^-$ | 270.98054 | 270.98135 | 0.810 | 2.989 | 32.8 |
| $C_7F_9S^-$ | 286.95770 | 286.95830 | 0.600 | 2.091 | 9097.8 |
| $C_7F_{11}^-$ | 292.98243 | 292.98310 | 0.670 | 2.287 | 20624.6 |
| $C_7F_9OS^-$ | 302.95261 | 302.95316 | 0.550 | 1.815 | 215.8 |
| $C_7HF_{10}S^-$ | 306.96393 | 306.96457 | 0.640 | 2.085 | 2024.9 |
| $C_7H_2F_{11}S^-$ | 326.97016 | 326.97074 | 0.580 | 1.774 | 57.4 |
| $C_8F_9O_2S^-$ | 330.94753 | 330.94823 | 0.700 | 2.115 | 2630.3 |
| $C_8HF_{10}O_2S^-$ | 350.95376 | 350.95440 | 0.640 | 1.824 | 485.6 |
| $C_8H_2F_{11}O_2S^-$ | 370.95999 | 370.96057 | 0.580 | 1.564 | 1695.0 |

**Figure S23.** MS2 spectrum of S-5:3 FTUCA and its major product ions generated by CID.

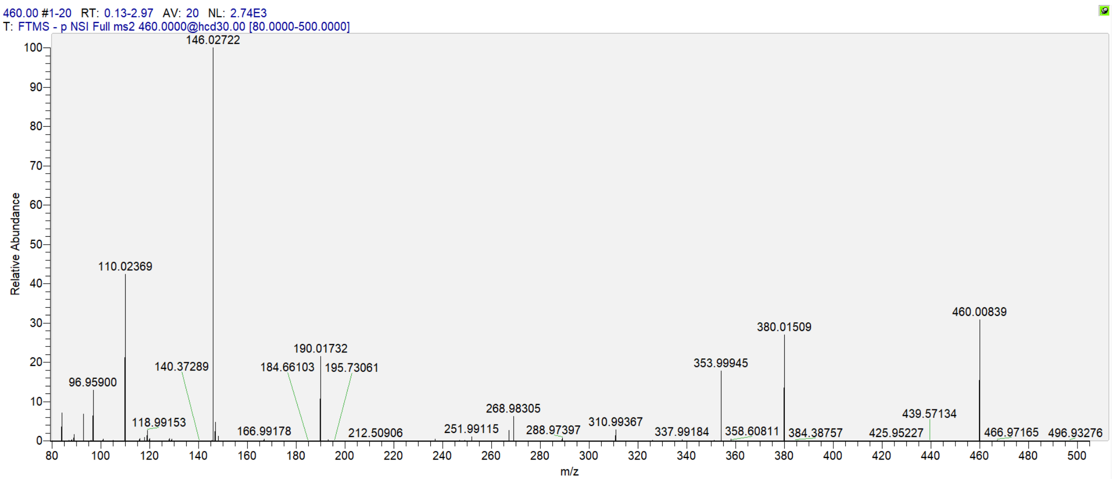


| Product Ions Formula | Theoretical m/z | Observed m/z | Absolute Mass Error (mDa) | Relative Mass Error (ppm) | MS2 Intensity |
|---|---|---|---|---|---|
| $C_3H_2NO_2^-$ | 84.00855 | 84.00796 | -0.590 | -7.023 | 199.9 |
| $C_3H_6NO_2^-$ | 88.03985 | 88.03930 | -0.550 | -6.247 | 18.5 |
| $C_3H_5O_2S^-$ | 105.00103 | 105.00047 | -0.560 | -5.333 | 17.2 |
| $C_3H_4NO_2S^-$ | 117.99627 | 117.99577 | -0.500 | -4.237 | 28.2 |
| $C_2F_5^-$ | 118.99202 | 118.99153 | -0.490 | -4.118 | 75.4 |
| $C_3H_6NO_2S^-$ | 120.01192 | 120.01143 | -0.490 | -4.083 | 27.5 |
| $C_5H_8NO_2S^-$ | 146.02757 | 146.02722 | -0.350 | -2.397 | 2742.5 |
| $C_6F_5^-$ | 166.99202 | 166.99178 | -0.240 | -1.437 | 24.8 |
| $C_6H_8NO_4S^-$ | 190.01740 | 190.01732 | -0.080 | -0.421 | 608.1 |
| $C_5F_7^-$ | 192.98882 | 192.98875 | -0.070 | -0.363 | 21.2 |
| $C_6F_5S^-$ | 198.96409 | 198.96410 | 0.010 | 0.050 | 12.6 |
| $C_7F_7^-$ | 216.98882 | 216.98901 | 0.190 | 0.876 | 12.9 |
| $C_7HF_8^-$ | 236.99505 | 236.99534 | 0.290 | 1.224 | 25.1 |
| $C_7F_7S^-$ | 248.96089 | 248.96137 | 0.480 | 1.928 | 17.3 |
| $C_9H_3F_5NS^-$ | 251.99064 | 251.99115 | 0.510 | 2.024 | 31.4 |
| $C_7HF_8S^-$ | 268.96712 | 268.96776 | 0.640 | 2.379 | 44.1 |
| $C_5F_{11}^-$ | 268.98243 | 268.98305 | 0.620 | 2.305 | 175.4 |
| $C_7H_2F_9S^-$ | 288.97335 | 288.97397 | 0.620 | 2.146 | 30.7 |
| $C_{11}H_9F_{11}NO_4S^-$ | 460.00766 | 460.00839 | 0.730 | 1.587 | 860.4 |

**Figure S24.** MS2 spectrum of CyS-5:3 FTCA and its major product ions generated by CID.

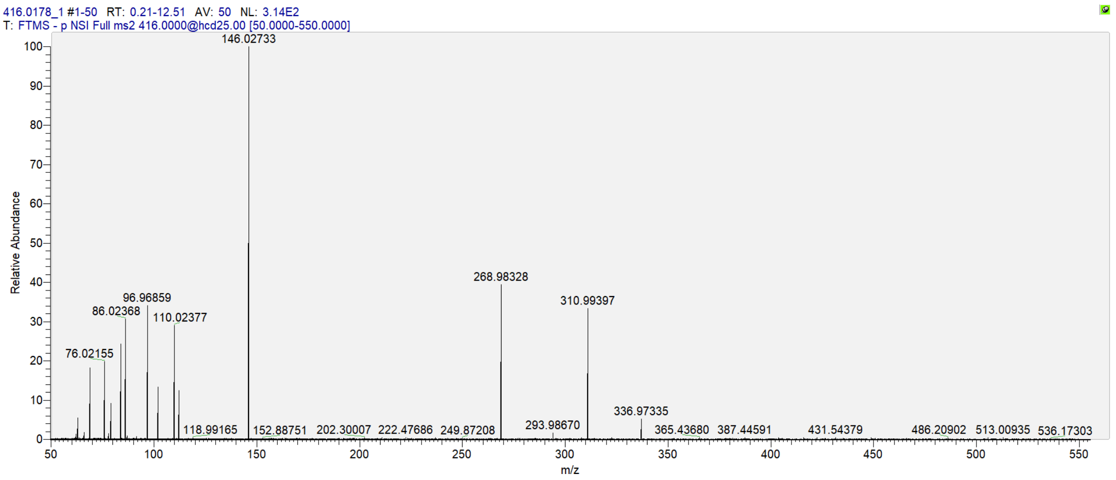


| Product Ions Formula | Theoretical m/z | Observed m/z | Absolute Mass Error (mDa) | Relative Mass Error (ppm) | MS2 Intensity |
|---|---|---|---|---|---|
| $C_3HO_2^-$ | 68.99765 | 68.99710 | -0.550 | -7.971 | 57.3 |
| $C_2H_6NS^-$ | 76.02210 | 76.02155 | -0.550 | -7.235 | 65.1 |
| $C_2F_5^-$ | 118.99202 | 118.99165 | -0.370 | -3.109 | 9.7 |
| $C_5H_8NO_2S^-$ | 146.02757 | 146.02733 | -0.240 | -1.644 | 314.5 |
| $C_6H_6F_5O_2S^-$ | 237.00087 | 237.00122 | 0.350 | 1.477 | 9.4 |
| $C_5F_{11}^-$ | 268.98243 | 268.98328 | 0.850 | 3.160 | 123.8 |
| $C_8F_{11}O_2^-$ | 336.97226 | 336.97335 | 1.090 | 3.235 | 26.7 |
| $C_{10}H_9F_{11}NO_2S^-$ | 416.01783 | 416.01879 | 0.960 | 2.380 | 6283.2 MS1 |

**Figure S25.** MS2 spectrum of CAS-5:3 FTCA and its major product ions generated by CID.

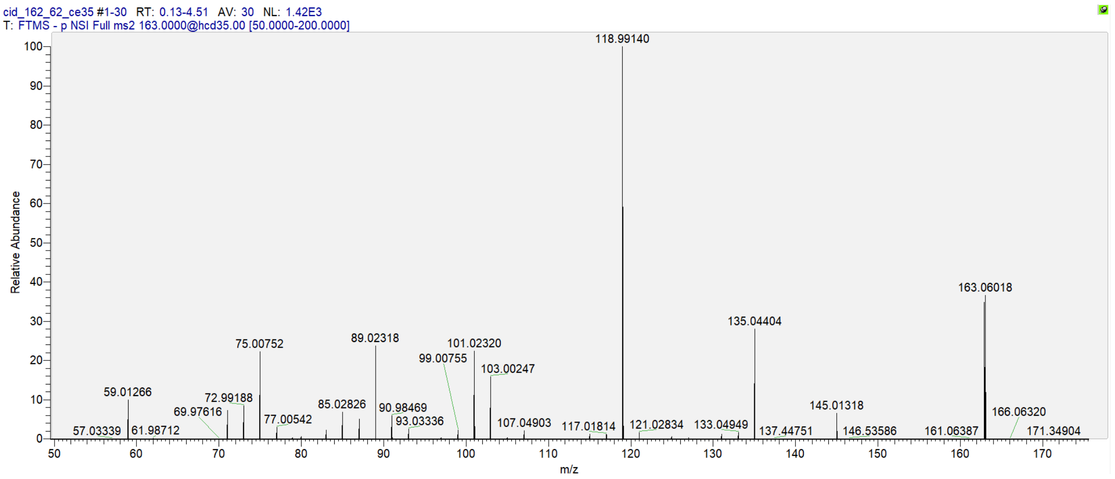


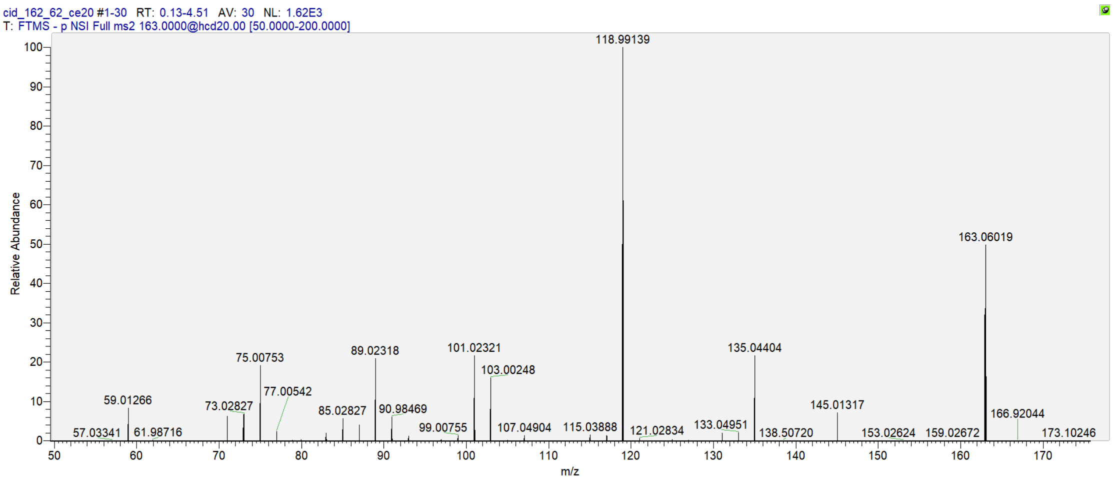


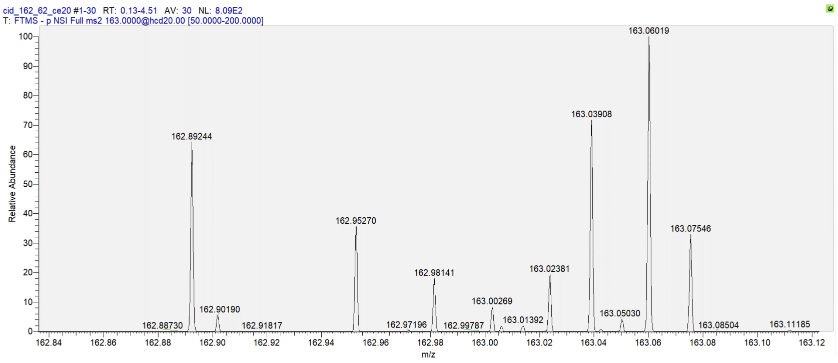


| Product Ions Formula | Theoretical m/z | Observed m/z | Absolute Mass Error (mDa) | Relative Mass Error (ppm) | MS2 Intensity |
|---|---|---|---|---|---|
| $CF_3^-$ | 68.99521 | 68.99451 | -0.700 | -10.146 | 6.0 |
| $C_2F_5^-$ | 118.99202 | 118.99140 | -0.620 | -5.210 | 1431.3 |
| $C_3F_5O_2^-$ | 162.98185 | 162.98141 | -0.440 | -2.700 | 145.7 (CE=20V) |

**Figure S26.** MS2 spectrum of PFPrA and its major product ions generated by CID.

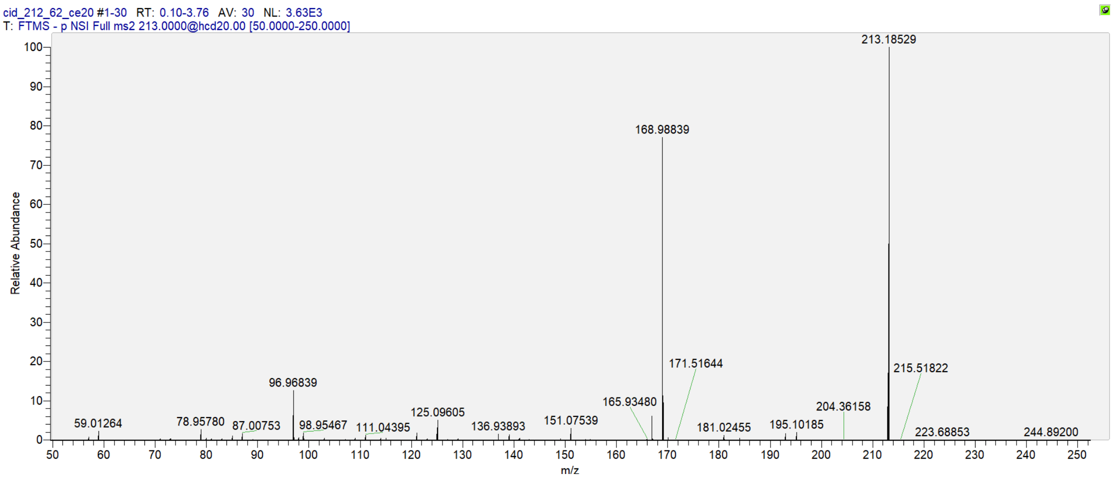


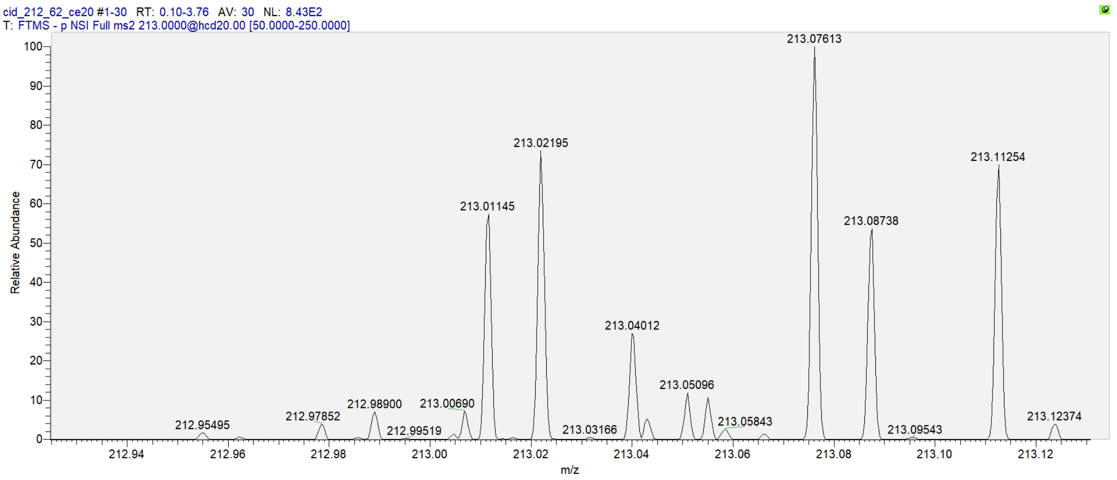


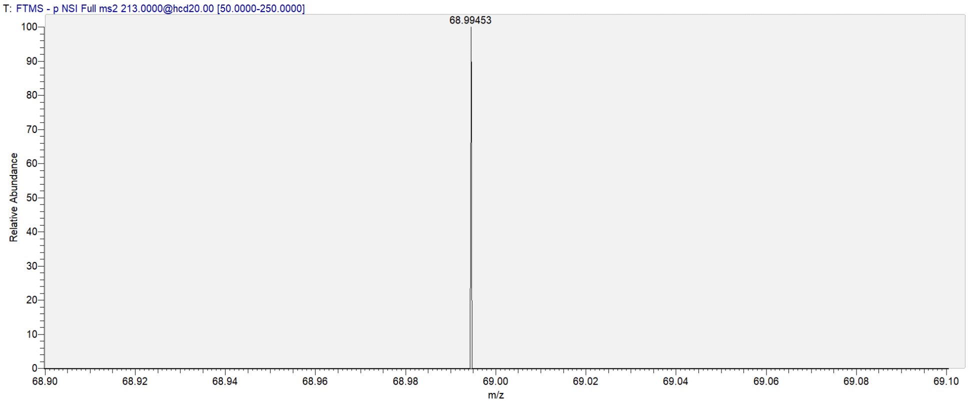


| Product Ions Formula | Theoretical m/z | Observed m/z | Absolute Mass Error (mDa) | Relative Mass Error (ppm) | MS2 Intensity |
|---|---|---|---|---|---|
| $CF_3^-$ | 68.99521 | 68.99453 | -0.680 | -9.856 | 28.0 |
| $C_3F_7^-$ | 168.98882 | 168.98839 | -0.430 | -2.545 | 2813.8 |
| $C_4F_7O_2^-$ | 212.97865 | 212.97852 | -0.130 | -0.610 | 33.9 |

**Figure S27.** MS2 spectrum of PFBA and its major product ions generated by CID.

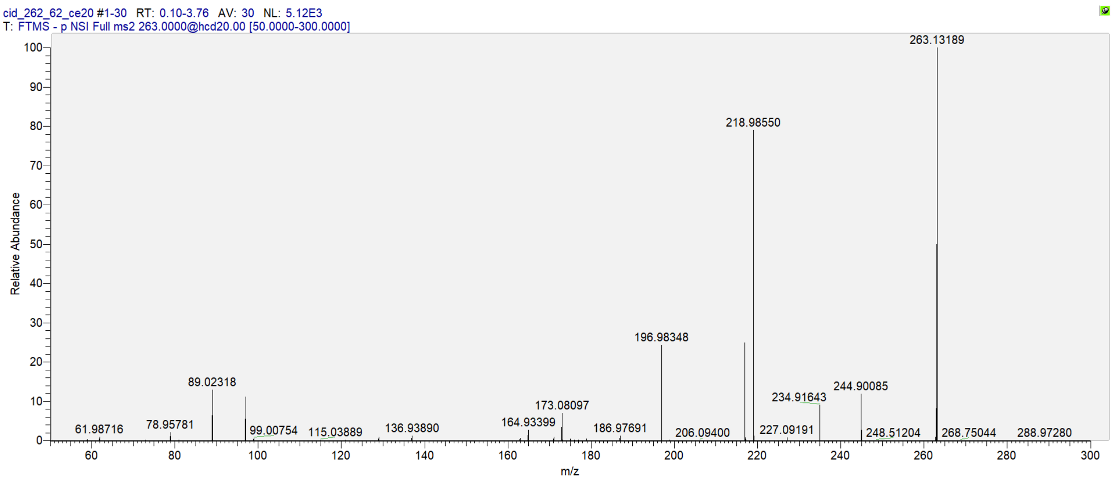


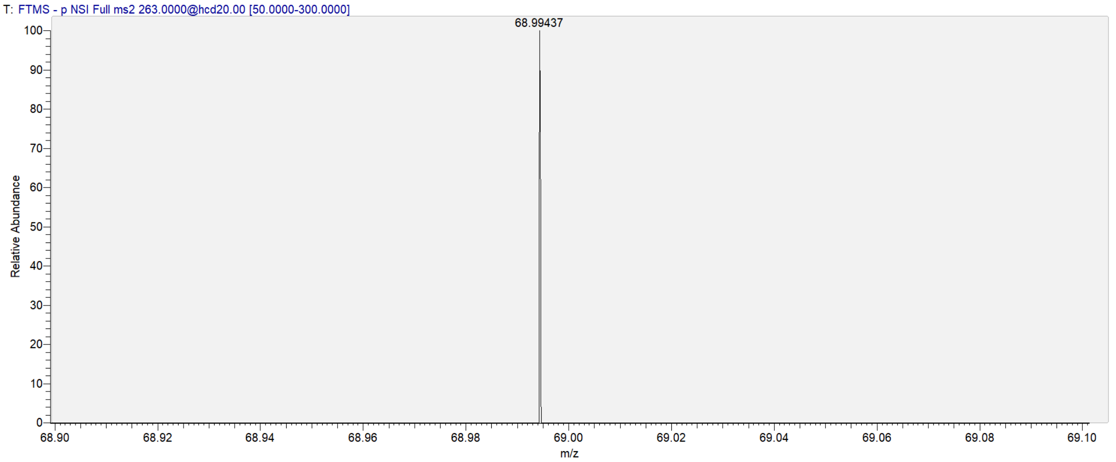


| Product Ions Formula | Theoretical m/z | Observed m/z | Absolute Mass Error (mDa) | Relative Mass Error (ppm) | MS2 Intensity |
|---|---|---|---|---|---|
| $CF_3^-$ | 68.99521 | 68.99437 | -0.840 | -12.175 | 10.1 |
| $C_4F_7O^-$ | 196.98374 | 196.98348 | -0.260 | -1.320 | 1260.4 |
| $C_4F_9^-$ | 218.98563 | 218.98550 | -0.130 | -0.594 | 4055.9 |
| $C_5F_9O_2^-$ | 262.97546 | 262.97568 | 0.220 | 0.837 | 104.5 |

**Figure S28.** MS2 spectrum of PFPeA and its major product ions generated by CID.

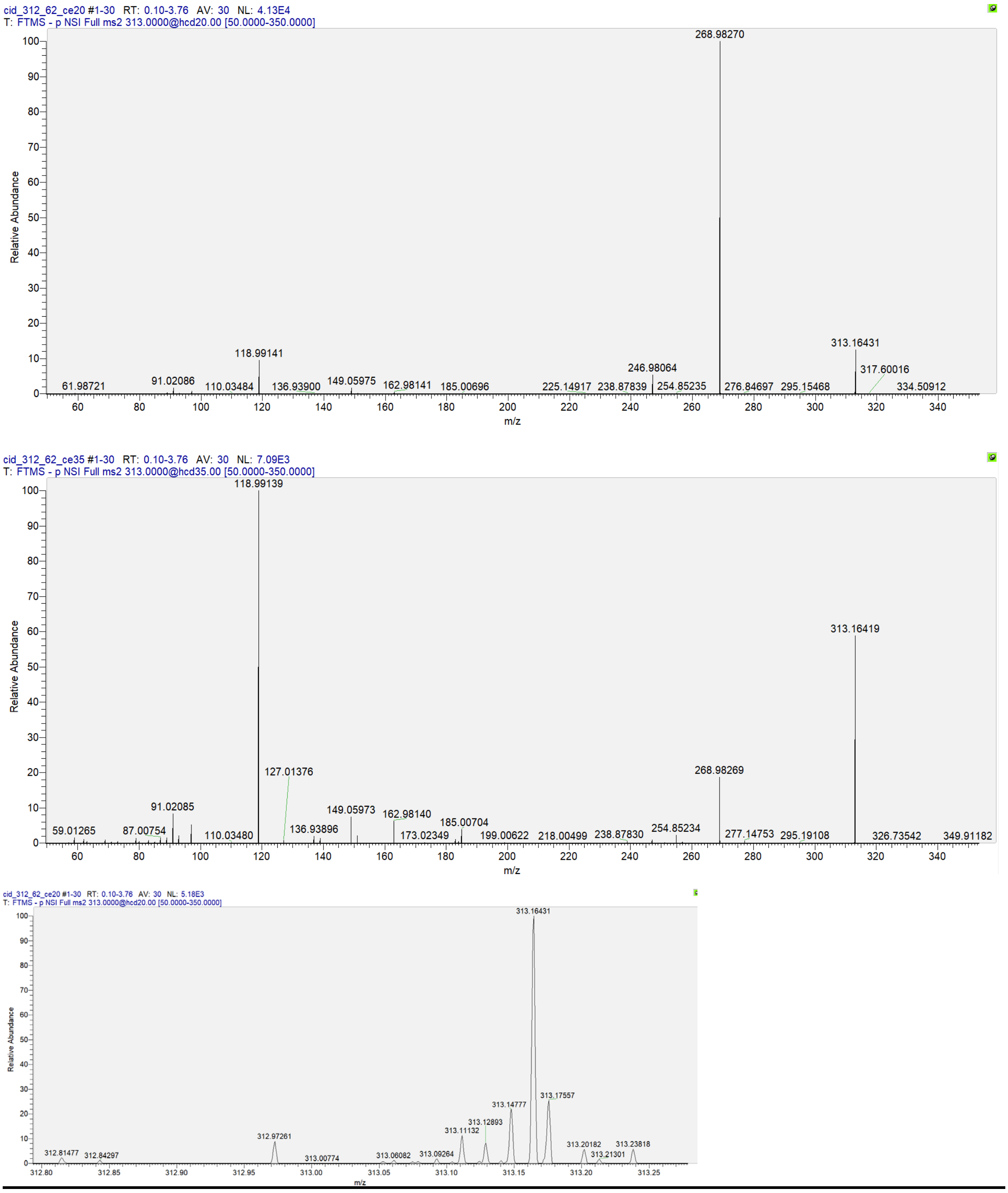


| Product Ions Formula | Theoretical m/z | Observed m/z | Absolute Mass Error (mDa) | Relative Mass Error (ppm) | MS2 Intensity |
|---|---|---|---|---|---|
| $CF_3^-$ | 68.99521 | 68.99453 | -0.680 | -9.856 | 68.2 (CE=35V) |
| $C_2F_5^-$ | 118.99202 | 118.99141 | -0.610 | -5.126 | 4098.5 |
| $C_3F_5O_2^-$ | 162.98185 | 162.98141 | -0.440 | -2.700 | 240.7 |
| $C_5F_9O^-$ | 246.98054 | 246.98064 | 0.100 | 0.405 | 2300.3 |
| $C_5F_{11}^-$ | 268.98243 | 268.98270 | 0.270 | 1.004 | 42305.1 |
| $C_6F_{11}O_2^-$ | 312.97226 | 312.97261 | 0.350 | 1.118 | 468.0 |

**Figure S29.** MS2 spectrum of PFHxA and its major product ions generated by CID.

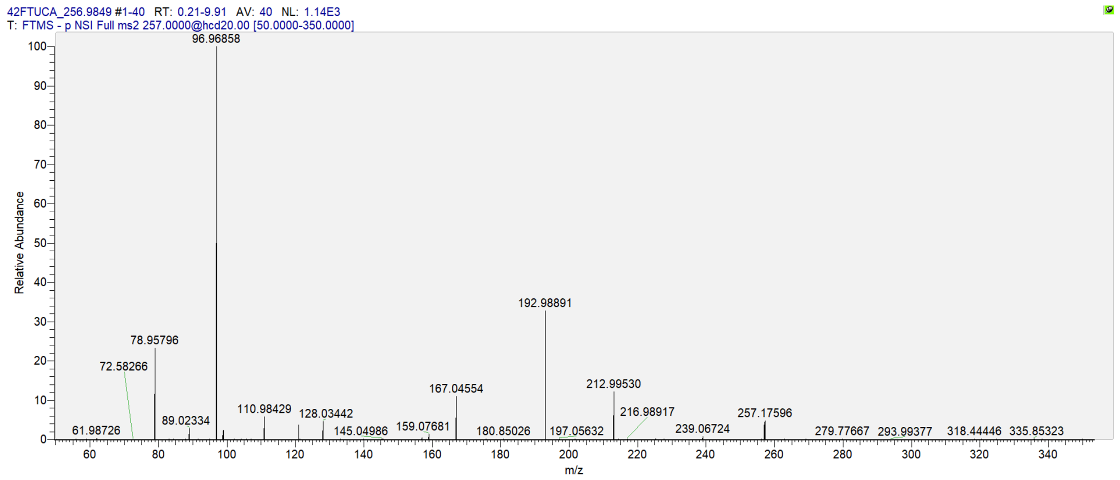


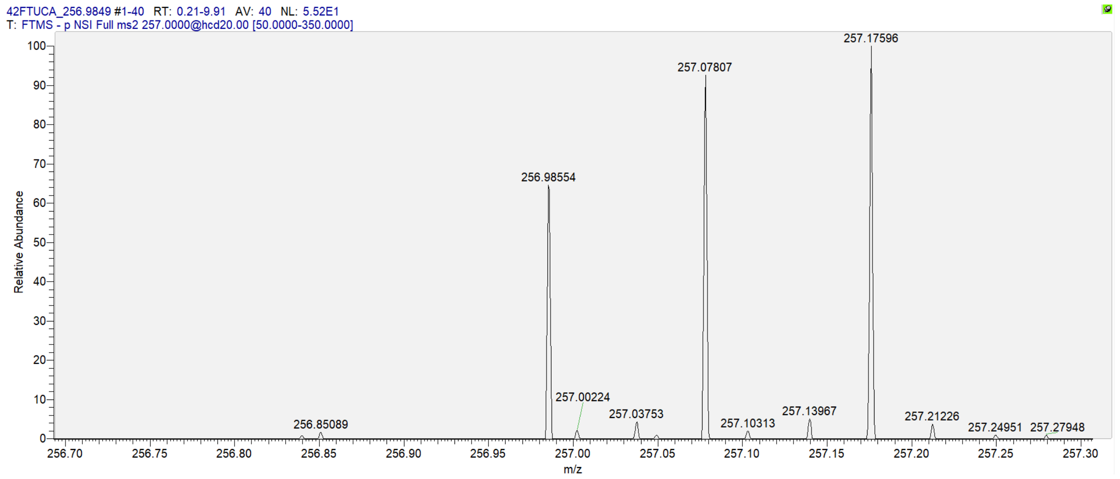


| Product Ions Formula | Theoretical m/z | Observed m/z | Absolute Mass Error (mDa) | Relative Mass Error (ppm) | MS2 Intensity |
|---|---|---|---|---|---|
| $C_5F_7^-$ | 192.98882 | 192.98891 | 0.090 | 0.466 | 375.9 |
| $C_5HF_8^-$ | 212.99505 | 212.99530 | 0.250 | 1.174 | 141.5 |
| $C_6HF_8O_2^-$ | 256.98488 | 256.98554 | 0.660 | 2.568 | 36.8 |

**Figure S30.** MS2 spectrum of 4:2 FTUCA and its major product ions generated by CID.

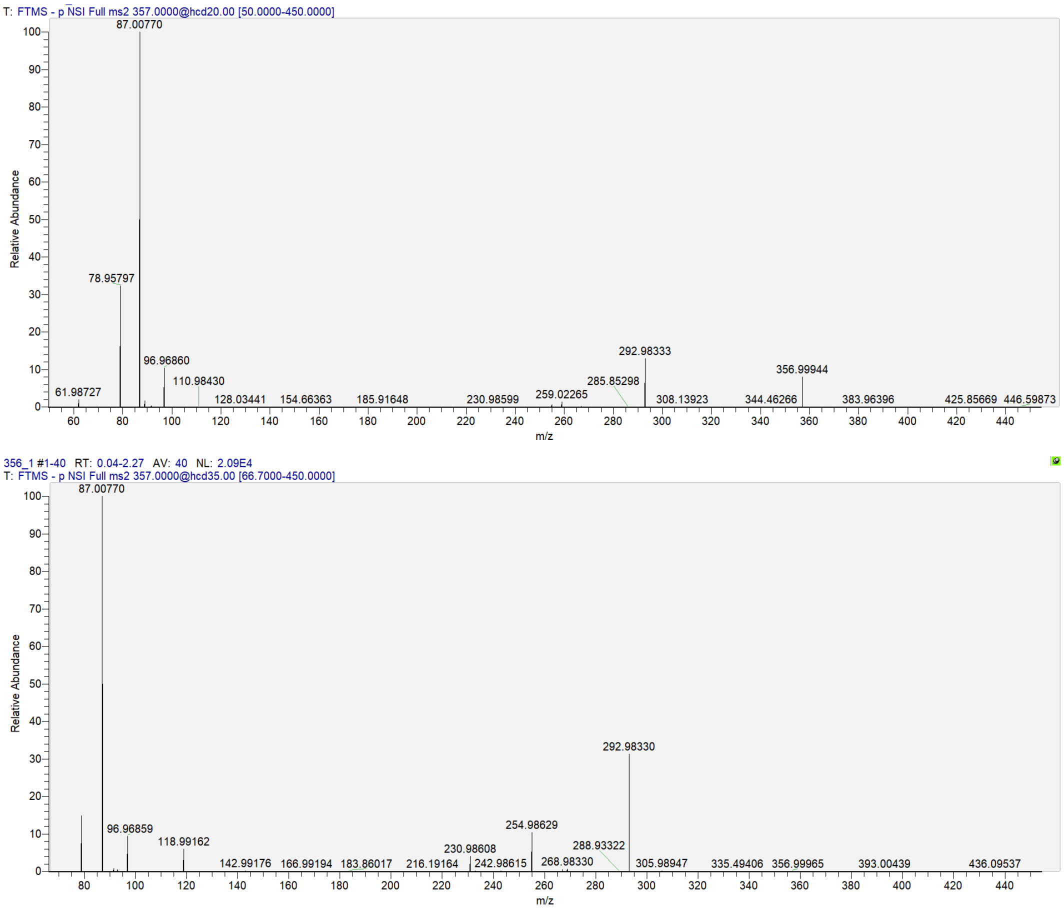


| Product Ions Formula | Theoretical m/z | Observed m/z | Absolute Mass Error (mDa) | Relative Mass Error (ppm) | MS2 Intensity |
|---|---|---|---|---|---|
| $CF_3^-$ | 68.99521 | 68.99465 | -0.560 | -8.117 | 20.2 |
| $C_3H_3O_3^-$ | 87.00822 | 87.00770 | -0.520 | -5.976 | 21003.4 |
| $C_3F_3^-$ | 92.99521 | 92.99470 | -0.510 | -5.484 | 97.5 |
| $C_2F_5^-$ | 118.99202 | 118.99162 | -0.400 | -3.362 | 1270.4 |
| $C_4F_5^-$ | 142.99202 | 142.99176 | -0.260 | -1.818 | 36.8 |
| $C_6F_5^-$ | 166.99202 | 166.99194 | -0.080 | -0.479 | 23.9 |
| $C_6F_7^-$ | 204.98882 | 204.98911 | 0.290 | 1.415 | 18.6 |
| $C_5F_9^-$ | 230.98563 | 230.98608 | 0.450 | 1.948 | 859.8 |
| $C_7F_7O^-$ | 232.98374 | 232.98423 | 0.490 | 2.103 | 58.6 |
| $C_6F_9^-$ | 242.98563 | 242.98615 | 0.520 | 2.140 | 56.6 |
| $C_7HF_8O^-$ | 252.98996 | 252.99058 | 0.620 | 2.451 | 29.9 |
| $C_7F_9^-$ | 254.98563 | 254.98629 | 0.660 | 2.588 | 2175.4 |
| $C_5F_{11}^-$ | 268.98243 | 268.98330 | 0.870 | 3.234 | 153.3 |
| $C_7F_{11}^-$ | 292.98243 | 292.98330 | 0.870 | 2.969 | 6583.3 |
| $C_7H_3F_{10}O^-$ | 293.00242 | 293.00318 | 0.760 | 2.594 | 21.6 |
| $C_8H_4F_{11}O_3^-$ | 356.99848 | 356.99944 | 0.960 | 2.689 | 202.4 (CE=20V) |

**Figure S31.** MS2 spectrum of α-OH 5:3 FTCA and its major product ions generated by CID.

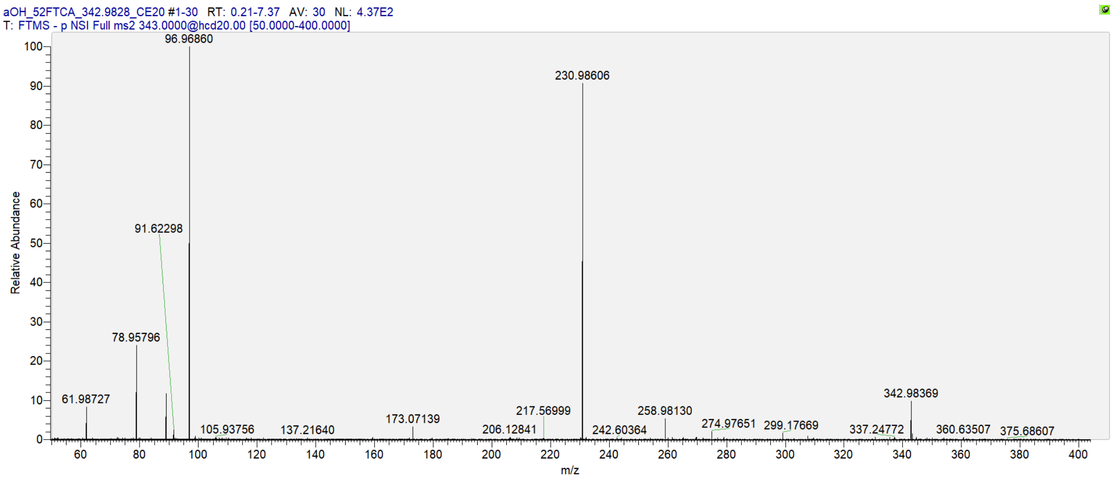


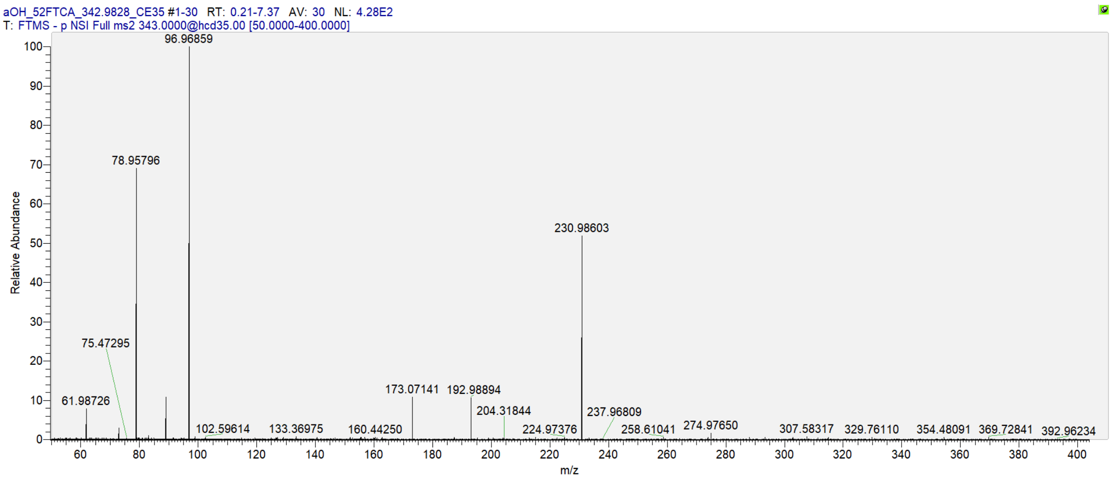


| Product Ions Formula | Theoretical m/z | Observed m/z | Absolute Mass Error (mDa) | Relative Mass Error (ppm) | MS2 Intensity |
|---|---|---|---|---|---|
| $C_2HO_3^-$ | 72.99257 | 72.99200 | -0.570 | -7.809 | 12.9 (CE=35V) |
| $C_5F_7^-$ | 192.98882 | 192.98894 | 0.120 | 0.622 | 46.0 (CE=35V) |
| $C_5F_9^-$ | 230.98563 | 230.98606 | 0.430 | 1.862 | 408.4 |
| $C_6F_9O^-$ | 258.98054 | 258.98130 | 0.760 | 2.935 | 24.7 |
| $C_6F_9O_2^-$ | 274.97546 | 274.97651 | 1.050 | 3.819 | 19.6 |
| $C_6HF_{10}O^-$ | 278.98677 | 278.98758 | 0.810 | 2.903 | 5.9 |
| $C_7H_2F_{11}O_3^-$ | 342.98283 | 342.98369 | 0.860 | 2.507 | 43.4 |

**Figure S32.** MS2 spectrum of α-OH 5:2 FTCA and its major product ions generated by CID.

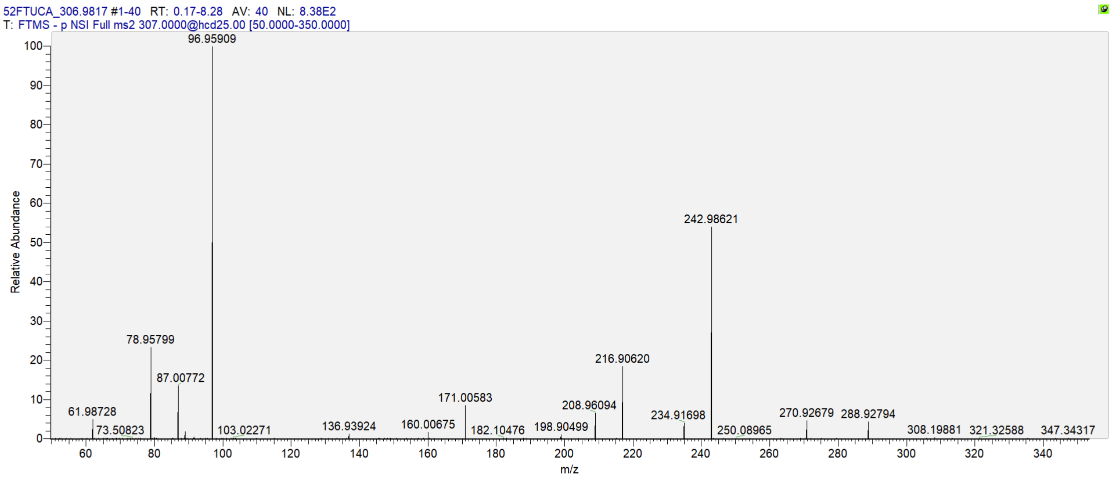


| Product Ions Formula | Theoretical m/z | Observed m/z | Absolute Mass Error (mDa) | Relative Mass Error (ppm) | MS2 Intensity |
|---|---|---|---|---|---|
| $C_3F_7^-$ | 168.98882 | 168.98897 | 0.150 | 0.888 | 11.2 |
| $C_6F_9^-$ | 242.98563 | 242.98621 | 0.580 | 2.387 | 457.3 |
| $C_7HF_{10}O_2^-$ | 306.98169 | 306.98248 | 0.790 | 2.573 | 31322.9 MS1 |

**Figure S33.** MS2 spectrum of 5:2 FTUCA and its major product ions generated by CID.

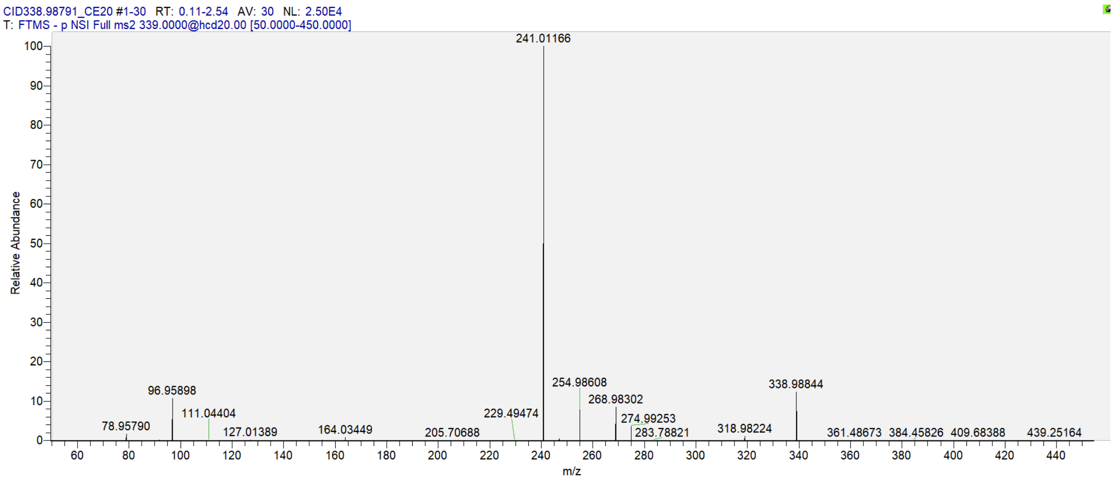


| Product Ions Formula | Theoretical m/z | Observed m/z | Absolute Mass Error (mDa) | Relative Mass Error (ppm) | MS2 Intensity |
|---|---|---|---|---|---|
| $C_5F_9O^-$ | 246.98054 | 246.98093 | 0.390 | 1.579 | 117.6 |
| $C_7HF_8O^-$ | 252.98996 | 252.99027 | 0.310 | 1.225 | 20.2 |
| $C_7F_9^-$ | 254.98563 | 254.98608 | 0.450 | 1.765 | 2010.0 |
| $C_5F_{11}^-$ | 268.98243 | 268.98302 | 0.590 | 2.193 | 2131.1 |
| $C_7HF_{10}^-$ | 274.99186 | 274.99253 | 0.670 | 2.436 | 931.1 |
| $C_8F_9O_2^-$ | 298.97546 | 298.97601 | 0.550 | 1.840 | 12.0 |
| $C_8HF_{10}O_2^-$ | 318.98169 | 318.98224 | 0.550 | 1.724 | 268.6 |
| $C_8F_{11}O_2^-$ | 336.97226 | 336.97279 | 0.530 | 1.573 | 28.1 |
| $C_8H_2F_{11}O_2^-$ | 338.98791 | 338.98844 | 0.530 | 1.563 | 3173.0 |

**Figure S34.** MS2 spectrum of 5:3 FTUCA and its major product ions generated by CID.

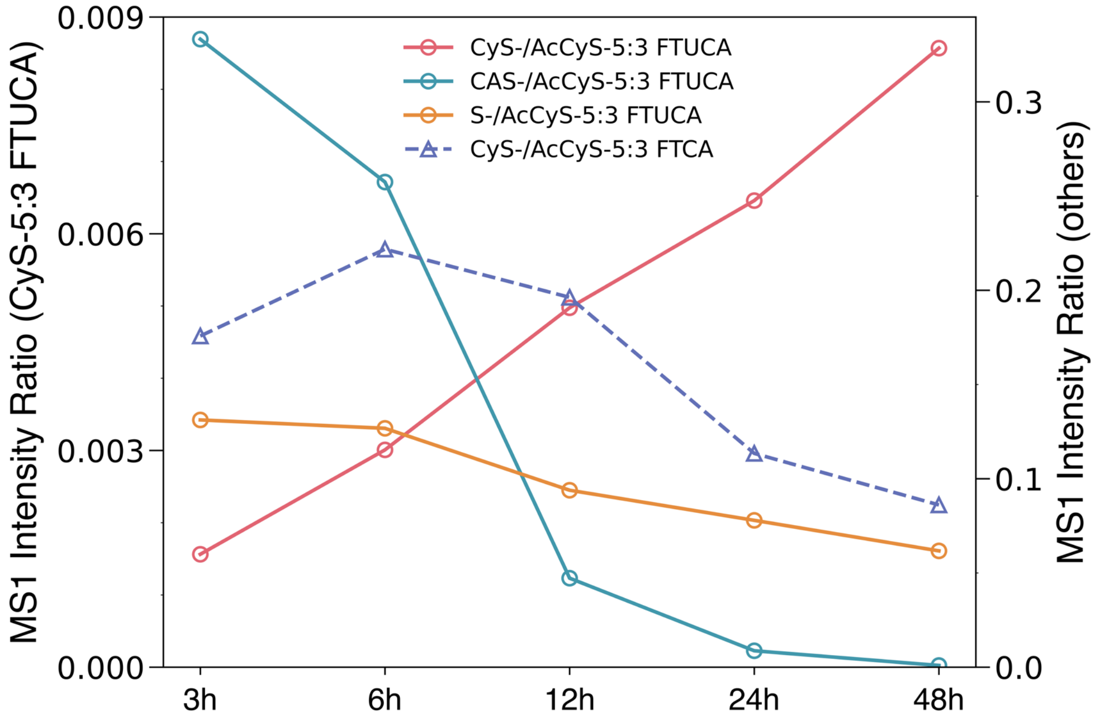


**Figure S35.** Ion intensity ratios between primary mercapturic acid derivatives (AcCyS-R) and their downstream thiol-containing conjugates (CyS-, CAS, and S-conjugates).

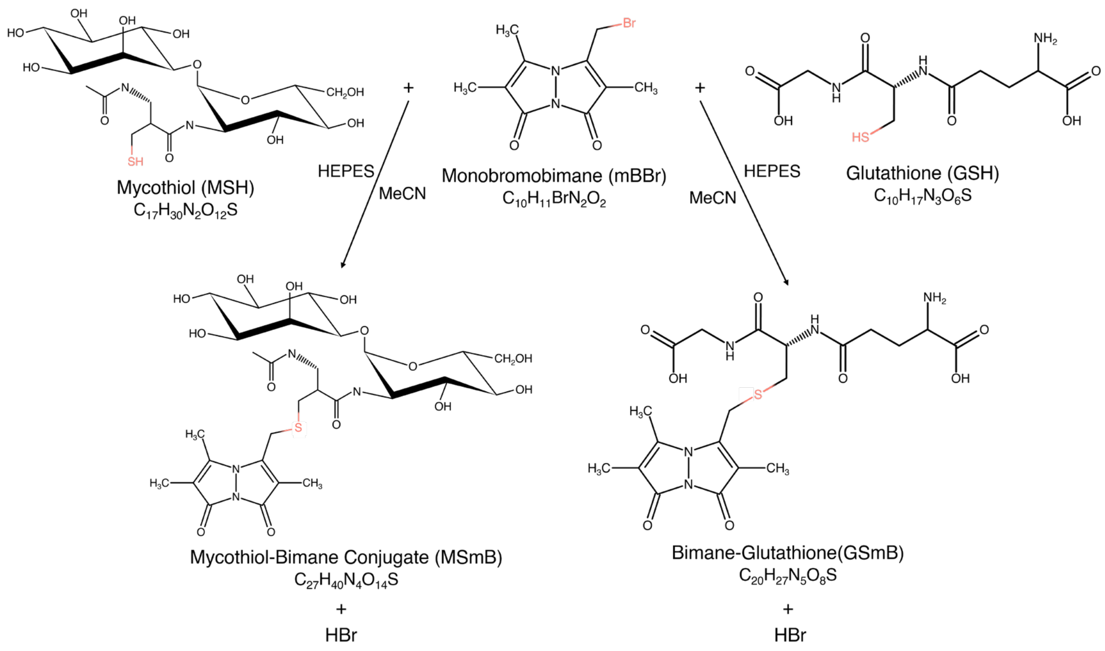


**Figure S36.** Derivatization of thiols (i.e., GSH and MSH) with monobromobimane (mBBr). The $S_N2$ reaction of mBBr with the thiol groups to produce thioether conjugates, MS-mBBr and GS-mBBr.

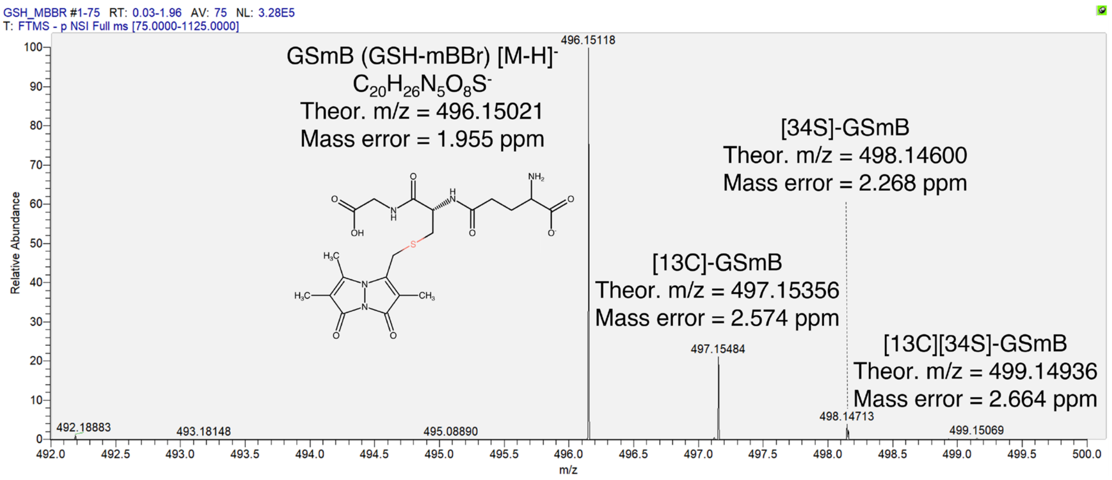


**Figure S37.** MS1 spectrum of GS-mBBr conjugate.

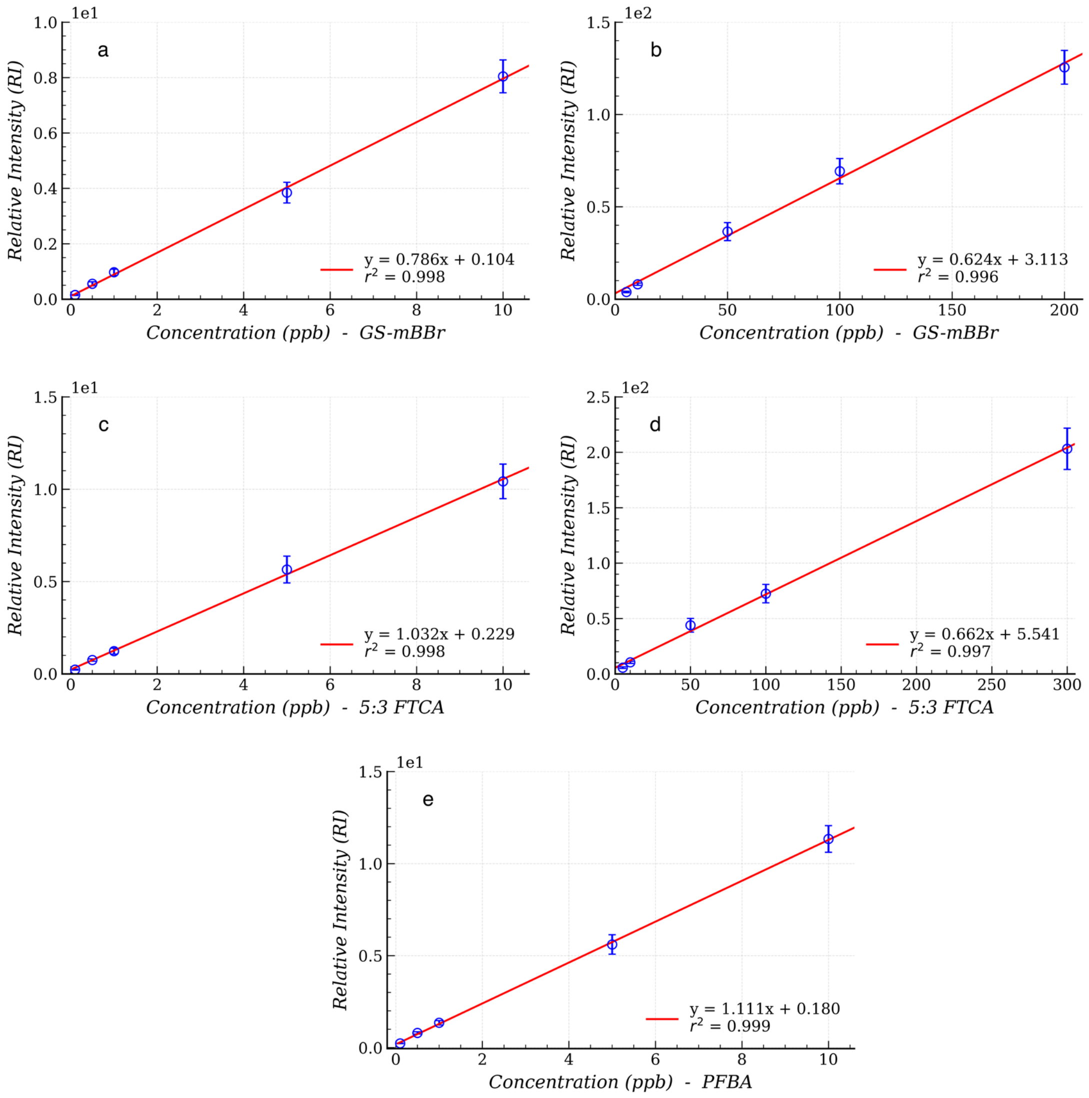


**Figure S38.** Calibration curves for semi-quantification. (a, b) Two segment (low- and high-concentration) linear calibration curves for the GS-mBBr. (c, d) Two segment (low- and high-concentration) linear calibration curves for the 5:3 FTCA. (e) Low-concentration curve for the PFBA calibration standard.

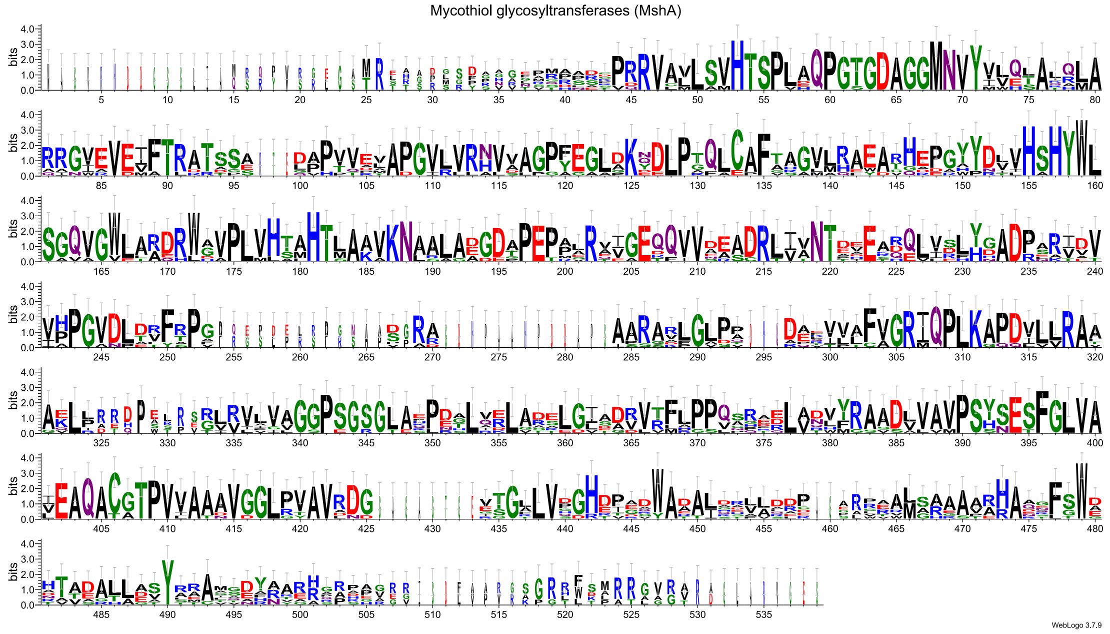


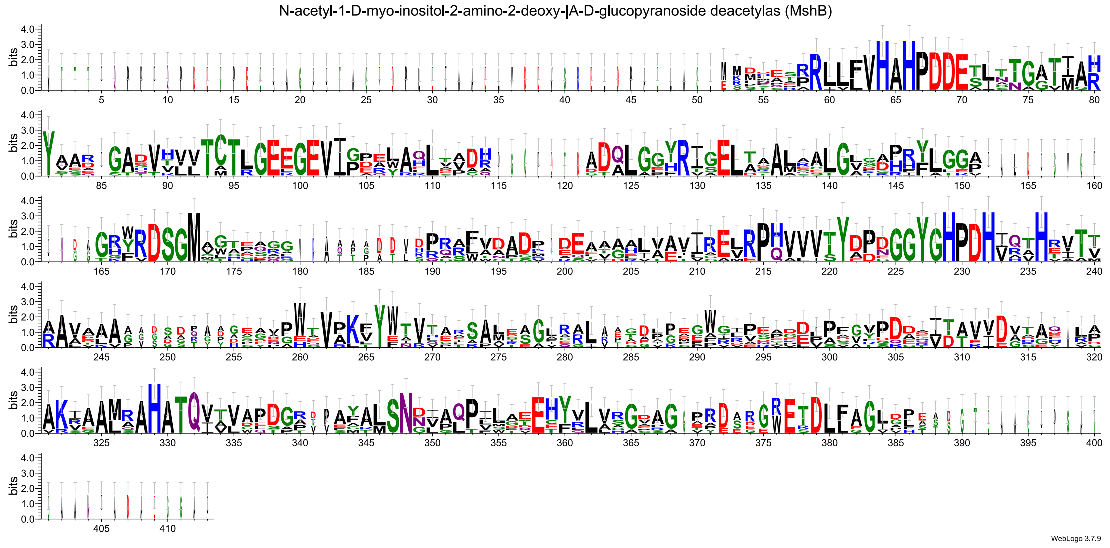


**Figure S39.** Logos of MshA (top) and MshB (bottom) protein sequence exhibiting conservativity from 8 representative *Actinomycetota* strains.

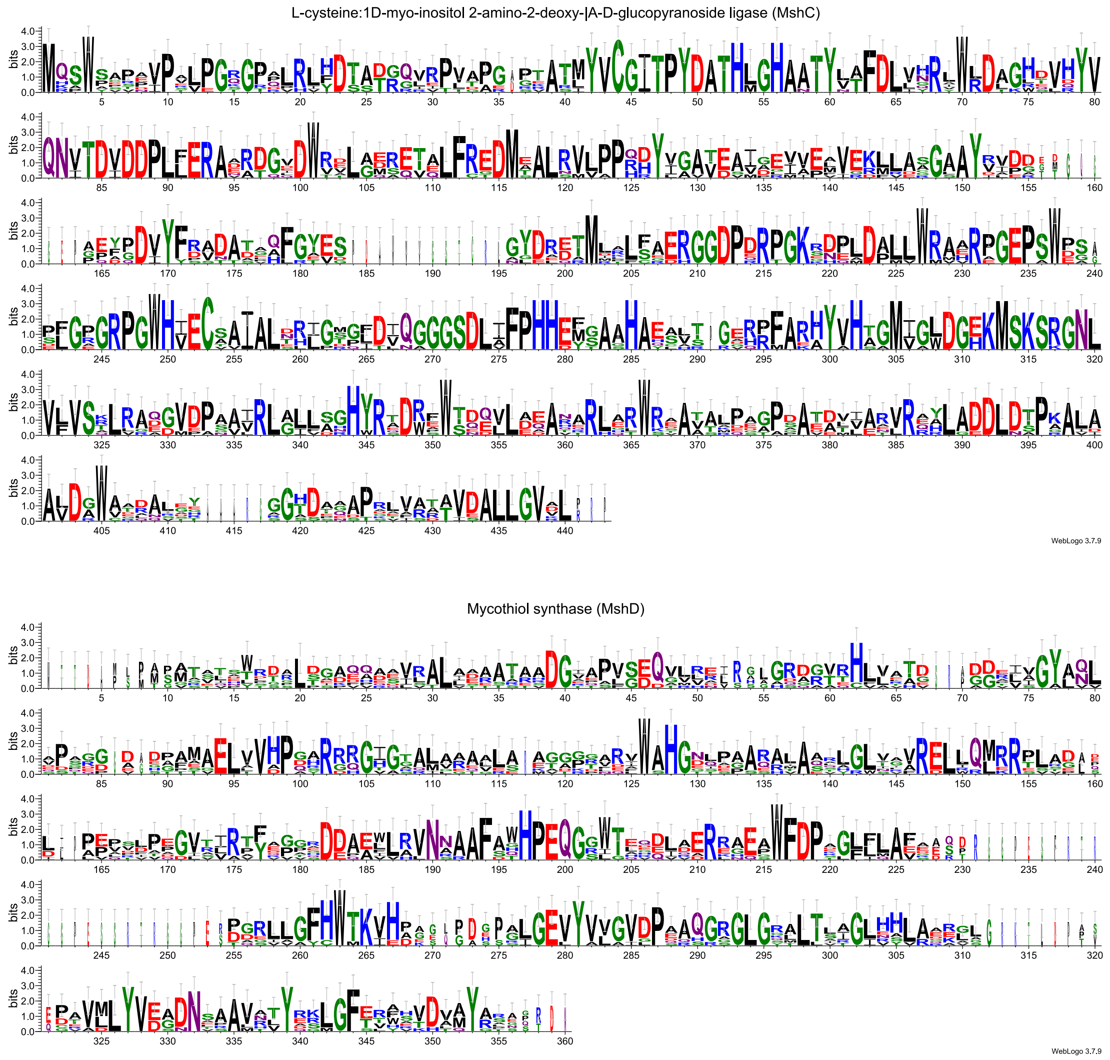


**Figure S40.** Logos of MshC (top) and MshD (bottom) protein sequence exhibiting conservativity from 8 representative *Actinomycetota* strains.

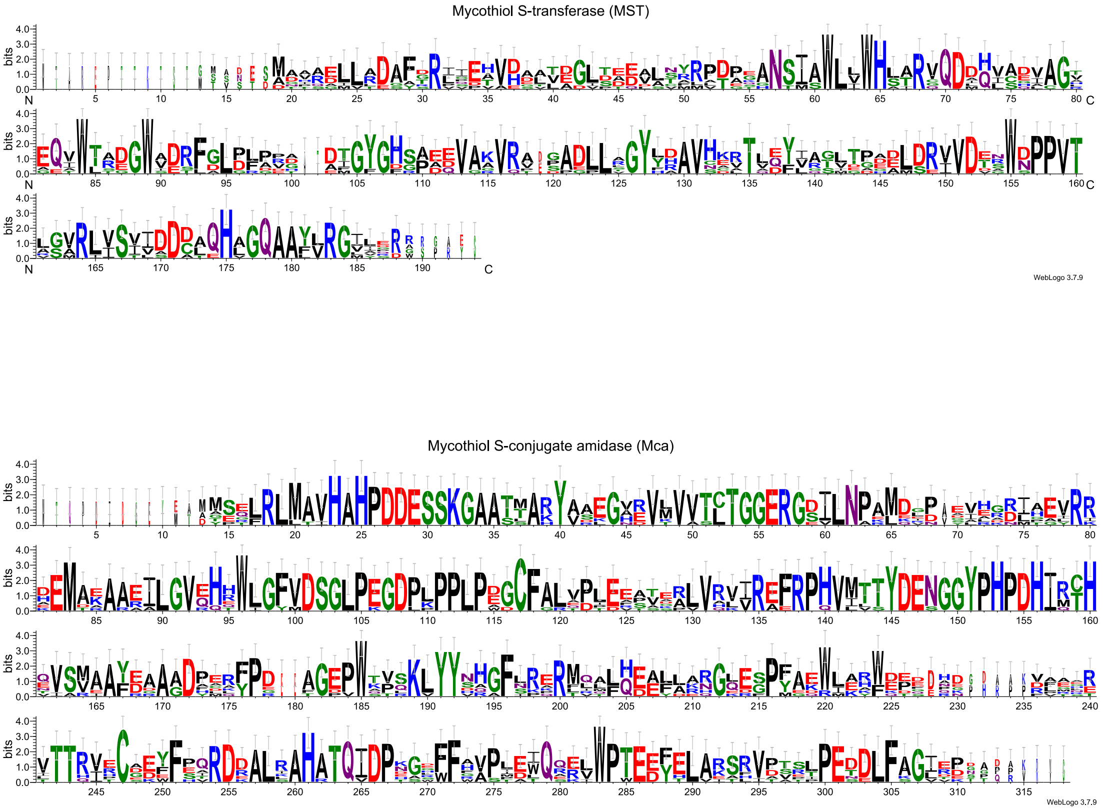


**Figure S41.** Logos of MST (top) and Mca (bottom) protein sequence exhibiting conservativity from 8 representative *Actinomycetota* strains.

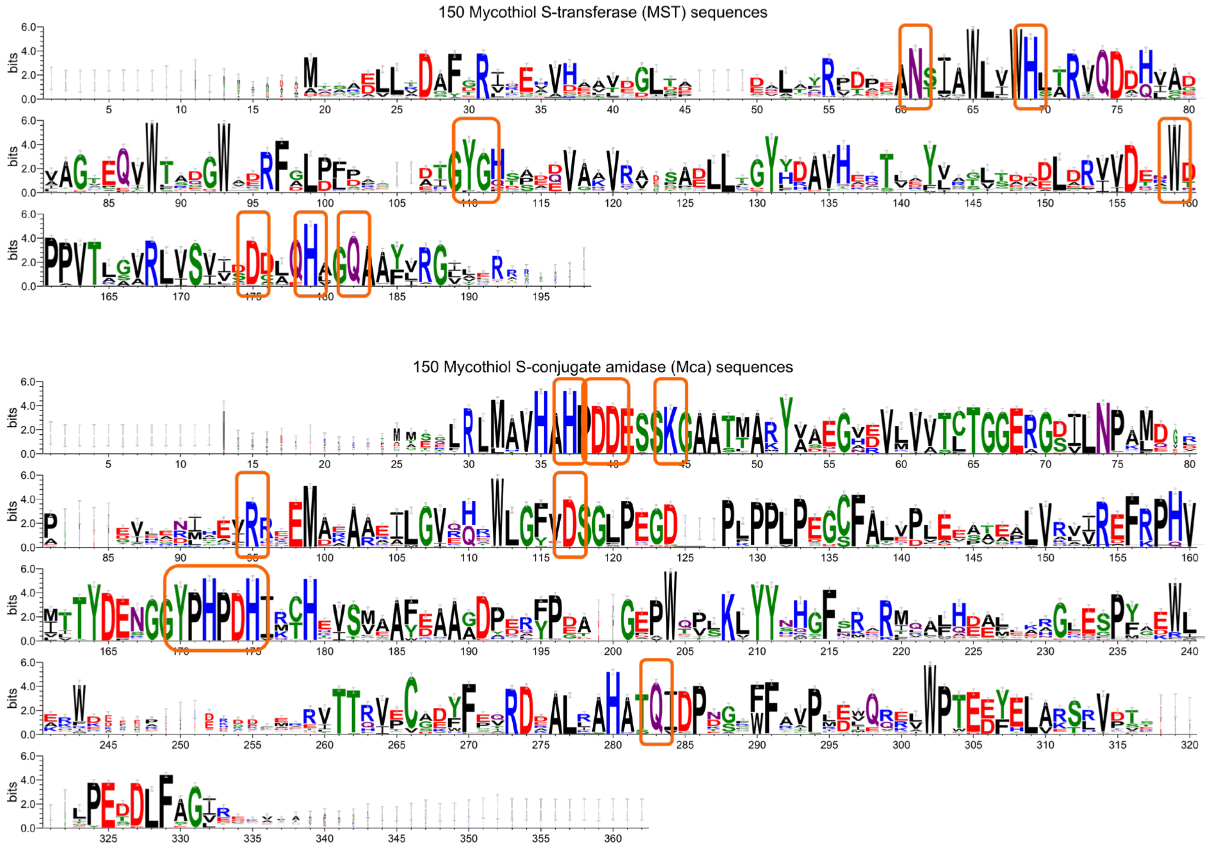


**Figure S42.** A broad sequence conservativity of MST and Mca catalytic residues. Logos were generated for MST (top) and Mca (bottom) from an alignment of 150 *Actinomycetota* strains, respectively.

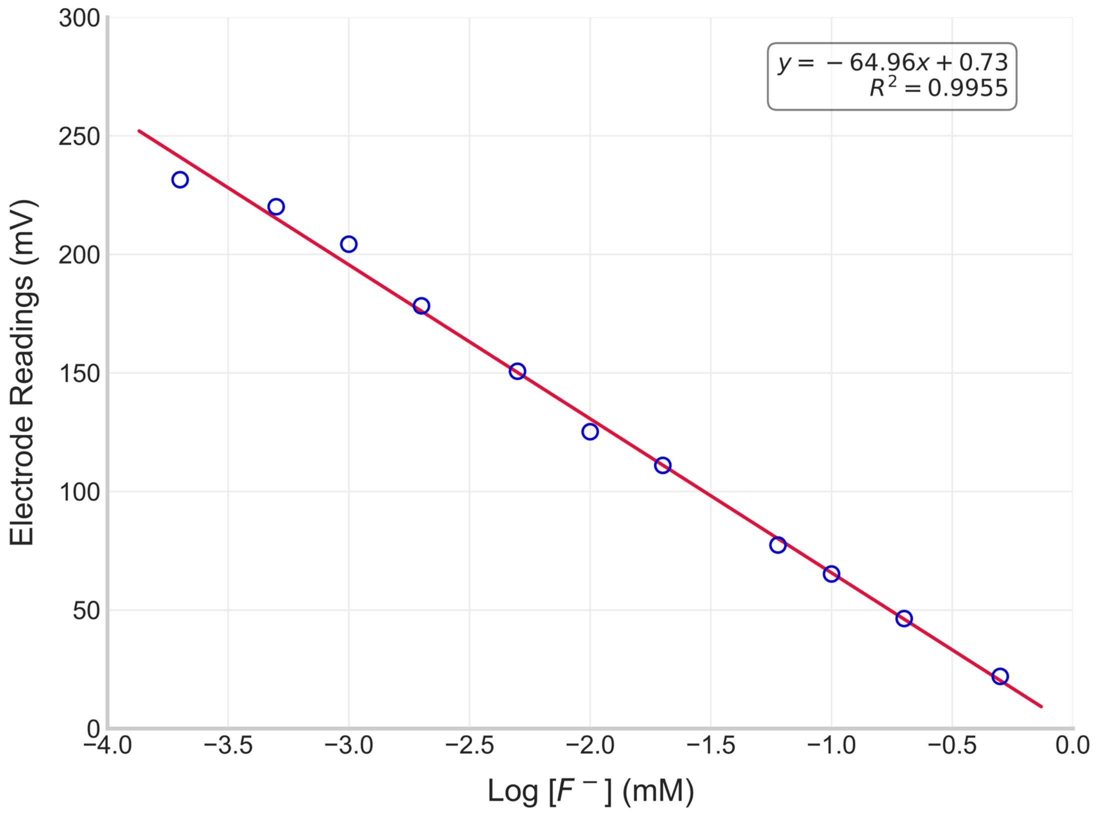


**Figure S43.** $F^-$ calibration curve.

**Table S1.** Bond-dissociation energies (BDEs) calculated for 6:2 and 5:3 FTCA [12, 33].

| Bonds in 6:2 FTCA | BDE kcal/mol | Bonds in 5:3 FTCA | BDE kcal/mol |
|---|---|---|---|
| Carboxyl C - Cα | 73.5 | Carboxyl C - Cα | 71.6 |
| Cα-β | 88.5 | Cα-β | 84.7 |
| Cβ-γ | 78.2 | Cβ-γ | 89.6 |
| Cγ-δ | 78.2 | Cγ-δ | 81.3 |
| Cδ-ε | 77.6 | Cδ-ε | 78.8 |
| Cε-ζ | 65.6 | Cε-ζ | 80.2 |
| Cζ-η | 84.7 | Cζ-η | 88.0 |
| C-H (Cα) | 95.2 | C-H (Cα) | 92.7 |
| C-F (Cβ) | 105.3 | C-H (Cβ) | 95.8 |
| C-F (Cγ) | 106.6 | C-F (Cγ) | 107.7 |
| C-F (Cδ) | 102.8 | C-F (Cδ) | 106.1 |
| C-F (Cε) | 102.6 | C-F (Cε) | 104.9 |
| C-F (Cζ) | 105.2 | C-F (Cζ) | 105.7 |
| C-F (Cη) | 115.4 | C-F (Cη) | 116.1 |

**Table S2.** Summary of previous studies that have reported PFAS biotransformation pathways that involves LMW thiols.

| Thiol/ Conjugating Agent | PFAS Precursor | Predicted Conjugation Site | Conjugation Mechanism | Pathway Importance | Reference |
|---|---|---|---|---|---|
| Glutathione (GSH) | 6:2 FTOH | Carboxylate group | Conjugative transformation at terminal functional group | Minor | Tseng et al., 2014 [34] |
| **Enzymes and Organisms**: *Phanerochaete chrysosporium* (white-rot fungus)<br>**Key Findings**: First evidence of identifying unique fungal-derived conjugation metabolites; biotransformation of 6:2 FTOH resulted in a high yield of 5:3 FTCA and a lower yield of terminal PFCAs and GSH conjugates | | | | | |
| Glutathione (GSH) | 6:2 FTOH | β-carbon | GSH conjugation | Minor | Zhang et al., 2020 [35] |
| **Enzymes and Organisms**: Alcohol and aldehyde dehydrogenase in soybean seedlings; glutathione S-transferase (GST)<br>**Key Findings**: Identification of GSH conjugates from 6:2 FTOH in plants; pathways of 6:2 FTOH biotransformation led to the dominant generation of primary PFCAs and 5:3 FTCA, accompanied by minor amount of GSH conjugates | | | | | |
| Glutathione (GSH) | 6:2 and 4:2 FTOH | β-carbon | GSH conjugation | Minor | Bhardwaj et al., 2024 [36] |
| **Enzymes and Organisms**: *Dietzia aurantiaca* strain J3; long-chain-fatty-acid-CoA ligases, acyl-CoA dehydrogenases, 3-hydroxyacyl-CoA dehydrogenase, glutathione S-transferase (GST)<br>**Key Findings**: Biotransformation of FTOHs by a pure bacterial culture formed GSH conjugates with pure bacterial strain treatment | | | | | |
| Glutathione (GSH) | 8:2 FTOH | β-carbon | GSH conjugation | Minor | Fasano et al., 2006 [37]<br>Fasano et al., 2009 [38]<br>Martin et al., 2005 [39]<br>Nabb et al., 2007 [40] |
| **Enzymes and Organisms**: Rat hepatocytes; Cytochrome P450 (P450) and glutathione transferases (GST)<br>**Key Findings**: *In vivo* and *in vitro* characterization of 8:2 FTOH biotransformation metabolism revealed the formation of PFCAs and glutathione conjugates | | | | | |
| Coenzyme A (CoA) | PFMeUPA | Carboxylate group | CoA ligation at the terminal functional group | Major | Yu et al., 2024 [41] |
| **Enzymes and Organisms**: CoA ligase (CarA) in *Acetobacterium* spp. (*A. bakii*)<br>**Key Findings**: CoA conjugates were formed in the process of reductive defluorination of PFMeUPA | | | | | |
| Coenzyme A (CoA) | 1:4 FTCA and 2:3 FTCA; 1:5 FTCA, 2:4 FTCA and 3:3 FTCA | Carboxylate group | Formation of CoA adduct at terminal functional group | Major | Mothersole et al., 2023 [42]<br>Mothersole et al., 2024 [43] |
| **Enzymes and Organisms**: *Gordonia* sp. strain NB4-1Y; Acyl-CoA synthetase (ACS)<br>**Key Findings**: Provided evidence for enzyme-catalyzed formation of PFAS-CoA adducts; reported enzyme selectivity based on chain length and fluorination level | | | | | |
| Mycothiol (MSH) | n:2 FTCA and n:3 FTUCA | β-carbon | SN2, E2 reaction to perform β-oxidation reaction at β-carbons of n:2 FTCAs; Michael addition at the β-carbons of n:3 FTUCAs | Major | This study |

**Table S3.** List of two FTCA parent compounds.

| Name | Acronym | Molecular Formula | Chemical Structure | Supplier | Molecular Weight | Anion *m/z* |
|---|---|---|---|---|---|---|
| 3,3,4,4,5,5,6,6,7,7,8,8,8-tridecafluorooctanoic acid | 6:2 FTCA | $C_8H_3F_{13}O_2$ | 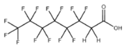 | SynQuest Labs, Inc. | 378.09 g/mol | 376.9847 |
| 2H,2H,3H,3H-perfluorooctanoic acid | 5:3 FTCA | $C_8H_5F_{11}O_2$ | 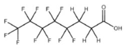 | SynQuest Labs, Inc. | 342.11 g/mol | 341.0036 |

**Table S4.** Experimental setup for FTCA biotransformation assays.

| *Experiment ID* | *Batch Type* | *Cell Type* | *Medium* | *FTCA Stock Addition* | *Analyte ConC.* | *Replicate* |
|---|---|---|---|---|---|---|
| T-62 | Treatment | Live RHA1 | 20× Diluted AMS | 20 μL 6:2 FTCA stock | 40 μM | |
| T-53 | Treatment | Live RHA1 | 20× Diluted AMS | 20 μL 5:3 FTCA stock | 40 μM | |
| AC-62 | Abiotic Control | Killed RHA1 | 20× Diluted AMS | 20 μL 6:2 FTCA stock | 40 μM | |
| AC-53 | Abiotic Control | Killed RHA1 | 20× Diluted AMS | 20 μL 5:3 FTCA stock | 40 μM | |
| AC-mix | Abiotic Control | Killed RHA1 | - | 20 μL of each FTCA stock | 40 μM of each FTCA | |
| AB-AMS | Abiotic Blank | Killed RHA1 | 20× Diluted AMS | - | - | |
| FTCA-62 | Analytical Control | - | 20× Diluted AMS | 20 μL 6:2 FTCA stock | 40 μM | |
| FTCA-53 | Analytical Control | - | 20× Diluted AMS | 20 μL 5:3 FTCA stock | 40 μM | Triplicate |
| FTCA-mix | Analytical Control | - | - | 20 μL of each FTCA stock | 40 μM of each FTCA | |
| MB | Microbial Blank | Live RHA1 | - | - | - | |
| MB-AMS | Microbial Blank | Live RHA1 | 20× Diluted AMS | - | - | |
| LRB-AMS | Lab Reagent Blank-AMS | - | 20× Diluted AMS | - | - | |
| LRB-MeOH | Lab Reagent Blank-MeOH | - | 20× Diluted AMS | - | - | |
| LRB-MeCN | Lab Reagent Blank-MeCN | - | 20× Diluted AMS | - | - | |
| Bg | Background | - | - | - | - | |

**Table S5.** List of TPs identified in FTCA biotransformation assays with their MS characteristics.

| Name | Molecular Formula | Anion Exact Mass | Observed Anion Mass | Mass error (ppm) | Max. Intensity | Time point of max. Intensity | Conf. level | CID |
|---|---|---|---|---|---|---|---|---|
| AcCyS-5:3 FTUCA | $C_{13}H_{10}F_{11}NO_5S$ | 500.00258 | 500.00333 | 1.500 | 5e6 | 5th round | 2b | Y |
| AcCyS-4:3 FTUCA | $C_{12}H_{10}F_9NO_5S$ | 450.00577 | 450.00647 | 1.556 | 2e5 | 3rd round* | 2b | Y |
| AcCyS-3:3 FTUCA | $C_{11}H_{10}F_7NO_5S$ | 400.00897 | 400.00954 | 1.425 | 4e4 | 5th round | 2b | Y |
| AcCyS-2:3 FTUCA | $C_{10}H_{10}F_5NO_5S$ | 350.01216 | 350.01286 | 1.999 | 6e3 | 1st round | 3d | N |
| MS-5:3 FTUCA | $C_{25}H_{31}F_{11}N_2O_{14}S$ | 823.12421 | 823.12477 | 0.680 | 6e4 | 5th round | 2b | Y |
| MS-3:3 FTUCA | $C_{23}H_{31}F_7N_2O_{14}S$ | 723.13060 | 723.13167 | 1.480 | 1e4 | 3rd round | 3d | N |
| PFHxA | $C_6HF_{11}O_2$ | 312.97226 | 312.97282 | 1.789 | 8e5 | 5th round | 1a | Y |
| PFPeA | $C_5HF_9O_2$ | 262.97546 | 262.97587 | 1.559 | 1e5 | 5th round | 1a | Y |
| PFBA | $C_4HF_7O_2$ | 212.97865 | 212.97872 | 0.329 | 4e4 | 5th round | 1a | Y |
| PFPrA | $C_3HF_5O_2$ | 162.98185 | 162.98152 | -2.025 | 1e4 | 5th round | 2c | Y |
| 6:2 FTUCA | $C_8H_2F_{12}O_2$ | 356.97849 | 356.97902 | 1.485 | - | Initial | 2b | Y |
| 5:2 FTUCA | $C_7H_2F_{10}O_2$ | 306.98169 | 306.98224 | 1.792 | 1e4 | 4th round | 3b | Y |
| 4:2 FTUCA | $C_6H_2F_8O_2$ | 256.98488 | 256.98536 | 1.868 | 6e4 | 5th round | 4 | Y |
| CyS-5:3 FTUCA | $C_{11}H_8F_{11}NO_4S$ | 457.99201 | 457.99246 | 0.983 | 2e4 | 5th round | 2b | Y |
| CyS-4:3 FTUCA | $C_{10}H_8F_9NO_4S$ | 407.99521 | 407.99556 | 0.858 | 9e3 | 3rd round* | 3d | N |
| S-5:3 FTUCA | $C_8H_3F_{11}O_2S$ | 370.95999 | 370.96031 | 0.863 | 3e5 | 5th round | 2b | Y |
| S-4:3 FTUCA | $C_7H_3F_9O_2S$ | 320.96318 | 320.96372 | 1.682 | 2e4 | 3rd round* | 3d | N |
| S-3:3 FTUCA | $C_6H_3F_7O_2S$ | 270.96637 | 270.96683 | 1.698 | 9e3 | 3rd round | 3d | N |
| CAS-5:3 FTUCA | $C_{10}H_8F_{11}NO_2S$ | 414.00218 | 414.00302 | 2.029 | 5e5 | SR-6h | 2b | Y |
| CAS-4:3 FTUCA | $C_9H_8F_9NO_2S$ | 364.00538 | 364.00624 | 2.363 | 2e4 | SR-4h* | 3d | N |
| CAS-3:3 FTUCA | $C_8H_8F_7NO_2S$ | 314.00857 | 314.00967 | 3.503 | 4e3 | SR-12h | 3d | N |
| 5:3 FTUCA | $C_8H_3F_{11}O_2$ | 338.98791 | 338.98856 | 1.917 | 2e5 | SR-4h | 2b | Y |
| 4:3 FTUCA | $C_7H_3F_9O_2$ | 288.99111 | 288.99168 | 1.972 | 9e3 | 4th round | 3d | N |
| 3:3 FTUCA | $C_6H_3F_7O_2$ | 238.99430 | 238.99477 | 1.967 | 7e3 | 1st round | 3d | N |
| 5:3 FTCA | $C_8H_5F_{11}O_2$ | 341.00356 | 341.00405 | 1.437 | 1e5 | SR-2h | 1a | Y |
| 4:3 FTCA | $C_7H_5F_9O_2$ | 291.00676 | 291.00758 | 2.818 | 8e3 | 4th round | 3d | N |
| α-OH 5:3 FTCA | $C_8H_5F_{11}O_3$ | 356.99848 | 356.99939 | 2.549 | 1e5 | SR-4h* | 3a | Y |
| α-OH 4:3 FTCA | $C_7H_5F_9O_3$ | 307.00167 | 307.00244 | 2.508 | 2e4 | SR-3h* | 4 | N |
| AcCyS-5:3 FTCA | $C_{13}H_{12}F_{11}NO_5S$ | 502.01823 | 502.01890 | 1.335 | 7e5 | 3rd round* | 2b | Y |

| | | | | | | | | |
|---|---|---|---|---|---|---|---|---|
| AcCyS-4:3 FTCA | $C_{12}H_{12}F_9NO_5S$ | 452.02142 | 452.02271 | 2.854 | 7e3 | 1st round | 3d | N |
| CyS-5:3 FTCA | $C_{11}H_{10}F_{11}NO_4S$ | 460.00766 | 460.00871 | 2.283 | 2e4 | 5th round | 2b | Y |
| CyS-4:3 FTCA | $C_{10}H_{10}F_9NO_4S$ | 410.01086 | 410.01163 | 1.878 | 5e3 | SR-24h | 3d | N |
| CAS-5:3 FTCA | $C_{10}H_{10}F_{11}NO_2S$ | 416.01783 | 416.01879 | 2.308 | 1e4 | SR-4h | 2b | Y |
| S-5:3 FTCA | $C_8H_5F_{11}O_2S$ | 372.97564 | 372.97612 | 1.099 | 5e3 | 4th round | 4 | N |
| α-OH 5:2 FTCA | $C_7H_2F_{11}O_3^-$ | 342.98283 | 342.98365 | 2.391 | 7e4 | SR-16h | 2b | Y |

**Table S6.** Detailed time-series fluorine mass recovery during 48-h biotransformation of 6:2 FTCA by RHA1.

| *6:2 FTCA Treatment* | 0h | 2h | 4h | 6h | 8h | 10h | 12h | 16h | 20h | 24h | 32h | 40h | 48h |
|---|---|---|---|---|---|---|---|---|---|---|---|---|---|
| Δ6:2 FTCA (µM) | 0.00 | 20.76 | 32.42 | 37.39 | 39.12 | 39.78 | 39.81 | 39.83 | 39.82 | 39.83 | 39.83 | 39.82 | 39.82 |
| Conjugation TPs (µM) | 0.00 | 10.53 | 19.08 | 21.41 | 21.52 | 21.02 | 20.70 | 19.18 | 17.67 | 17.35 | 16.02 | 15.16 | 14.93 |
| Conjugation TPs/Δ6:2 FTCA (µM/µM) | - | 50.72% | 58.86% | 57.25% | 55.00% | 52.83% | 51.98% | 48.16% | 44.37% | 43.56% | 40.22% | 38.06% | 37.49% |
| Conjugation TPs/Total 6:2 FTCA Consumed (µM/µM) | 0.00 | 26.44% | 47.91% | 53.75% | 54.04% | 52.77% | 51.97% | 48.16% | 44.37% | 43.56% | 40.22% | 38.07% | 37.49% |
| PFCAs TPs (µM) | 0.00 | 0.13 | 0.22 | 0.42 | 0.72 | 1.26 | 1.67 | 1.82 | 1.87 | 1.89 | 1.97 | 2.12 | 2.28 |
| PFCA TPs /Δ6:2 FTCA (µM/µM) | - | 0.60% | 0.68% | 1.12% | 1.84% | 3.18% | 4.18% | 4.56% | 4.69% | 4.74% | 4.96% | 5.33% | 5.73% |
| PFCA TPs /Total 6:2 FTCA Consumed (µM/µM) | 0.00 | 0.31% | 0.55% | 1.06% | 1.81% | 3.18% | 4.18% | 4.56% | 4.69% | 4.74% | 4.96% | 5.33% | 5.73% |
| “One-Carbon Removal” TPs (µM) | 0.00 | 2.74 | 3.35 | 2.98 | 1.69 | 0.96 | 0.81 | 0.74 | 0.80 | 0.82 | 0.89 | 0.92 | 0.93 |
| “One-Carbon Removal” TPs /Δ6:2 FTCA (µM/µM) | - | 13.21% | 10.35% | 7.96% | 4.31% | 2.42% | 2.04% | 1.86% | 2.00% | 2.07% | 2.23% | 2.31% | 2.34% |
| “One-Carbon Removal” TPs /Total 6:2 FTCA Consumed (µM/µM) | 0.00 | 6.89% | 8.42% | 7.48% | 4.24% | 2.42% | 2.04% | 1.86% | 2.00% | 2.07% | 2.23% | 2.31% | 2.34% |
| Measured TPs/Δ6:2 FTCA (µM/µM) | - | 64.53% | 69.88% | 66.34% | 61.15% | 58.43% | 58.21% | 54.57% | 51.06% | 50.36% | 47.40% | 45.70% | 45.56% |
| Measured TPs /Total 6:2 FTCA Consumed (µM/µM) | 0.00% | 33.64% | 56.88% | 62.28% | 60.08% | 58.36% | 58.19% | 54.57% | 51.06% | 50.36% | 47.40% | 45.70% | 45.56% |
| $F^-$ Release (µM) | 0.00 | 43.84 | 63.66 | 71.23 | 75.86 | 78.59 | 81.05 | 83.34 | 84.51 | 85.34 | 86.48 | 87.51 | 87.96 |
| $F^-$ Release/Δ6:2 FTCA (µM/µM) | - | 2.11 | 1.96 | 1.90 | 1.94 | 1.98 | 2.04 | 2.09 | 2.12 | 2.14 | 2.17 | 2.20 | 2.21 |
| $F^-$ Release /Total 6:2 FTCA Consumed (µM/µM) | 0.00 | 1.10 | 1.60 | 1.79 | 1.90 | 1.97 | 2.04 | 2.09 | 2.12 | 2.14 | 2.17 | 2.20 | 2.21 |
| Theoretical $F^-$ Release from Conjugation TPs (µM) | 0.00 | 21.06 | 38.16 | 42.84 | 43.15 | 42.37 | 41.86 | 38.79 | 35.80 | 35.15 | 32.47 | 30.74 | 30.30 |
| Theoretical $F^-$ Release from Conjugation TPs/Δ$F^-$ Release (µM/µM) | - | 48.02% | 59.95% | 60.15% | 56.88% | 53.91% | 51.65% | 46.54% | 42.36% | 41.19% | 37.55% | 35.13% | 34.45% |
| Theoretical $F^-$ Release from Conjugation TPs/Total $F^-$ Release (µM/µM) | 0.00% | 23.94% | 43.38% | 48.70% | 49.05% | 48.17% | 47.59% | 44.10% | 40.69% | 39.96% | 36.92% | 34.95% | 34.45% |
| Fluorine Mass Precent (FMP) of Conjugation TPs (%) | 0.00% | 22.37% | 40.54% | 45.47% | 45.70% | 44.59% | 43.89% | 40.66% | 37.45% | 36.77% | 33.95% | 32.13% | 31.64% |
| Theoretical $F^-$ Release from PFCA TPs (µM) | 0.00 | 0.25 | 0.45 | 0.90 | 1.66 | 3.05 | 3.97 | 4.34 | 4.52 | 4.60 | 4.88 | 5.24 | 5.61 |
| Theoretical $F^-$ Release from PFCA TPs /Δ$F^-$ Release (µM/µM) | - | 0.58% | 0.71% | 1.27% | 2.18% | 3.88% | 4.90% | 5.21% | 5.35% | 5.39% | 5.65% | 5.99% | 6.38% |
| Theoretical $F^-$ Release from PFCA TPs /Total $F^-$ Release (µM/µM) | 0.00% | 0.29% | 0.51% | 1.02% | 1.88% | 3.46% | 4.51% | 4.93% | 5.14% | 5.23% | 5.55% | 5.96% | 6.38% |
| Fluorine Mass Precent (FMP) of PFCAs TPs (%) | 0.00% | 0.26% | 0.46% | 0.88% | 1.49% | 2.59% | 3.42% | 3.72% | 3.82% | 3.85% | 4.01% | 4.31% | 4.64% |
| Theoretical $F^-$ Release from "One-Carbon Removal" TPs (µM) | 0.00 | 5.49 | 6.71 | 5.97 | 3.52 | 2.38 | 2.30 | 2.14 | 2.29 | 2.42 | 2.56 | 2.64 | 2.71 |
| Theoretical $F^-$ Release from "One-Carbon Removal" TPs/Δ$F^-$ Release (µM/µM) | - | 12.51% | 10.54% | 8.38% | 4.65% | 3.03% | 2.84% | 2.57% | 2.71% | 2.83% | 2.97% | 3.02% | 3.08% |
| Theoretical $F^-$ Release from "One-Carbon Removal" TPs/Total $F^-$ Release (µM/µM) | 0.00% | 6.24% | 7.63% | 6.79% | 4.01% | 2.71% | 2.62% | 2.44% | 2.60% | 2.75% | 2.92% | 3.01% | 3.08% |
| Fluorine Mass Precent (FMP) of "One-Carbon Removal" TPs (%) | 0.00% | 5.83% | 7.13% | 6.32% | 3.56% | 1.96% | 1.59% | 1.44% | 1.56% | 1.60% | 1.73% | 1.80% | 1.82% |
| Theoretical $F^-$ Release from All TPs /Δ$F^-$ Release (µM/µM) | - | 61.12% | 71.19% | 69.80% | 63.71% | 60.82% | 59.39% | 54.32% | 50.41% | 49.41% | 46.16% | 44.14% | 43.91% |
| Theoretical $F^-$ Release from All TPs /Total $F^-$ Release (µM/µM) | 0.00% | 30.46% | 51.52% | 56.52% | 54.94% | 54.34% | 54.72% | 51.47% | 48.43% | 47.93% | 45.38% | 43.92% | 43.91% |
| Fluorine Mass Recovery (FMR, %) | - | 84.81% | 79.02% | 72.55% | 67.15% | 64.42% | 64.57% | 61.92% | 59.15% | 58.70% | 56.39% | 55.15% | 55.09% |

**Table S7.** Detailed time-series fluorine mass recovery during 48-h biotransformation of 5:3 FTCA by RHA1.

| *5:3 FTCA Treatment* | 0h | 2h | 4h | 6h | 8h | 10h | 12h | 16h | 20h | 24h | 32h | 40h | 48h |
|---|---|---|---|---|---|---|---|---|---|---|---|---|---|
| Δ5:3 FTCA (μM) | 0.00 | 40.23 | 43.78 | 43.74 | 43.67 | 43.59 | 43.54 | 43.58 | 43.55 | 43.49 | 43.56 | 43.62 | 43.65 |
| Conjugation TPs (μM) | 0.00 | 0.56 | 2.08 | 2.66 | 3.17 | 3.33 | 3.46 | 3.21 | 3.09 | 3.07 | 3.02 | 2.90 | 2.82 |
| Conjugation TPs/Δ5:3 FTCA (μM/μM) | - | 1.39% | 4.74% | 6.08% | 7.25% | 7.64% | 7.94% | 7.37% | 7.09% | 7.05% | 6.93% | 6.66% | 6.47% |
| Conjugation TPs/Total 5:3 FTCA Consumed (μM/μM) | 0.00 | 1.28% | 4.76% | 6.09% | 7.25% | 7.63% | 7.92% | 7.36% | 7.08% | 7.03% | 6.92% | 6.65% | 6.47% |
| PFCAs TPs (μM) | 0.00 | 0.32 | 0.37 | 0.41 | 0.44 | 0.46 | 0.47 | 0.46 | 0.44 | 0.44 | 0.42 | 0.41 | 0.39 |
| PFCA TPs /Δ5:3 FTCA (μM/μM) | - | 0.79% | 0.84% | 0.93% | 1.02% | 1.06% | 1.08% | 1.05% | 1.02% | 1.01% | 0.97% | 0.94% | 0.89% |
| PFCA TPs /Total 5:3 FTCA Consumed (μM/μM) | 0.00 | 0.73% | 0.84% | 0.94% | 1.02% | 1.06% | 1.08% | 1.04% | 1.01% | 1.00% | 0.97% | 0.94% | 0.89% |
| "One-Carbon Removal" TPs (μM) | 0.00 | 0.59 | 0.62 | 0.63 | 0.63 | 0.63 | 0.60 | 0.54 | 0.51 | 0.50 | 0.40 | 0.32 | 0.23 |
| "One-Carbon Removal" TPs /Δ5:3 FTCA (μM/μM) | - | 1.46% | 1.41% | 1.44% | 1.45% | 1.45% | 1.37% | 1.23% | 1.18% | 1.14% | 0.92% | 0.73% | 0.54% |
| "One-Carbon Removal" TPs /Total 5:3 FTCA Consumed (μM/μM) | 0.00 | 1.35% | 1.41% | 1.45% | 1.45% | 1.44% | 1.37% | 1.23% | 1.18% | 1.14% | 0.92% | 0.73% | 0.54% |
| Measured TPs/Δ5:3 FTCA (μM/μM) | - | 3.64% | 6.99% | 8.46% | 9.72% | 10.15% | 10.39% | 9.65% | 9.29% | 9.20% | 8.82% | 8.33% | 7.90% |
| Measured TPs /Total 5:3 FTCA Consumed (μM/μM) | 0.00% | 3.36% | 7.01% | 8.48% | 9.72% | 10.13% | 10.36% | 9.64% | 9.27% | 9.17% | 8.81% | 8.32% | 7.90% |
| $F^-$ Release (μM) | - | 4.84 | 7.50 | 9.00 | 10.05 | 10.69 | 11.27 | 11.83 | 12.13 | 12.36 | 12.67 | 12.96 | 13.17 |
| $F^-$ Release/Δ5:3 FTCA (μM/μM) | - | 0.12 | 0.17 | 0.21 | 0.23 | 0.25 | 0.26 | 0.27 | 0.28 | 0.28 | 0.29 | 0.30 | 0.30 |
| $F^-$ Release /Total 5:3 FTCA Consumed (μM/μM) | - | 0.11 | 0.17 | 0.21 | 0.23 | 0.24 | 0.26 | 0.27 | 0.28 | 0.28 | 0.29 | 0.30 | 0.30 |
| Theoretical $F^-$ Release from Conjugation TPs (μM) | 0.00 | 0.65 | 1.56 | 1.79 | 1.77 | 1.71 | 1.68 | 1.52 | 1.42 | 1.38 | 1.30 | 1.28 | 1.25 |
| Theoretical $F^-$ Release from Conjugation TPs/Δ$F^-$ Release (μM/μM) | - | 13.42% | 20.80% | 19.86% | 17.57% | 16.02% | 14.87% | 12.84% | 11.67% | 11.15% | 10.22% | 9.84% | 9.49% |
| Theoretical $F^-$ Release from Conjugation TPs/Total $F^-$ Release (μM/μM) | 0.00 | 4.93% | 11.85% | 13.58% | 13.40% | 13.01% | 12.73% | 11.53% | 10.75% | 10.46% | 9.84% | 9.69% | 9.49% |
| Fluorine Mass Precent (FMP) of Conjugation TPs (%) | 0.00% | 1.15% | 4.43% | 5.72% | 6.89% | 7.27% | 7.57% | 7.05% | 6.78% | 6.74% | 6.65% | 6.39% | 6.21% |
| Theoretical $F^-$ Release from PFCA TPs (μM) | 0.00 | 0.66 | 0.76 | 0.86 | 0.94 | 0.98 | 1.00 | 0.96 | 0.93 | 0.92 | 0.89 | 0.86 | 0.82 |
| Theoretical $F^-$ Release from PFCA TPs /Δ$F^-$ Release (μM/μM) | - | 13.54% | 10.14% | 9.51% | 9.34% | 9.13% | 8.84% | 8.16% | 7.71% | 7.46% | 7.00% | 6.67% | 6.26% |
| Theoretical $F^-$ Release from PFCA TPs /Total $F^-$ Release (μM/μM) | 0.00 | 4.98% | 5.78% | 6.50% | 7.12% | 7.41% | 7.57% | 7.32% | 7.10% | 7.00% | 6.74% | 6.56% | 6.26% |
| Fluorine Mass Precent (FMP) of PFCAs TPs (%) | 0.00 | 0.59% | 0.68% | 0.76% | 0.82% | 0.85% | 0.87% | 0.84% | 0.82% | 0.81% | 0.78% | 0.76% | 0.72% |
| Theoretical $F^-$ Release from "One-Carbon Removal" TPs (μM) | 0.00 | 0.18 | 0.09 | 0.05 | 0.01 | 0.00 | 0.00 | 0.00 | 0.00 | 0.00 | 0.00 | 0.00 | 0.00 |
| Theoretical $F^-$ Release from "One-Carbon Removal" TPs/Δ$F^-$ Release (μM/μM) | - | 3.80% | 1.18% | 0.50% | 0.14% | 0.00% | 0.00% | 0.00% | 0.00% | 0.00% | 0.00% | 0.00% | 0.00% |
| Theoretical $F^-$ Release from "One-Carbon Removal" TPs/Total $F^-$ Release (μM/μM) | 0.00 | 1.40% | 0.67% | 0.34% | 0.11% | 0.00% | 0.00% | 0.00% | 0.00% | 0.00% | 0.00% | 0.00% | 0.00% |
| Fluorine Mass Precent (FMP) of "One-Carbon Removal" TPs (%) | 0.00 | 1.31% | 1.39% | 1.44% | 1.45% | 1.44% | 1.37% | 1.23% | 1.18% | 1.14% | 0.92% | 0.73% | 0.54% |
| Theoretical $F^-$ Release from All TPs /Δ$F^-$ Release (μM/μM) | - | 30.77% | 32.13% | 29.87% | 27.04% | 25.16% | 23.72% | 20.99% | 19.38% | 18.60% | 17.23% | 16.51% | 15.75% |
| Theoretical $F^-$ Release from All TPs /Total $F^-$ Release (μM/μM) | 0.00 | 11.31% | 18.30% | 20.43% | 20.63% | 20.42% | 20.30% | 18.85% | 17.85% | 17.46% | 16.58% | 16.25% | 15.75% |
| Fluorine Mass Recovery (FMR, %) | - | 11.88% | 7.77% | 9.58% | 11.19% | 11.94% | 12.40% | 11.73% | 11.54% | 11.63% | 11.18% | 10.64% | 10.21% |

**Table S8.** Parameters of calibration regression for three calibration standards.

| | WLS Regression (1/x weighting) | Linear Regression low Conc. Segment | Linear Regression high Conc. Segment | Intersection (ppb) |
|---|---|---|---|---|
| GS-mBBr | $m = 0.7896$<br>$b = 0.0907$ | $m = 0.7856$<br>$b = 0.1039$ | $m = 0.6238$<br>$b = 3.1126$ | 18.6 |
| 5:3FTCA | $m = 1.0581$<br>$b = 0.1419$ | $m = 1.0319$<br>$b = 0.2289$ | $m = 0.6621$<br>$b = 5.5406$ | 14.4 |
| PFBA | $m = 1.1211$<br>$b = 0.1451$ | $m = 1.1107$<br>$b = 0.1795$ | | |

**Table S9.** Gene and protein accession numbers for MST, Mca, and other MSH biosynthesis enzymes in eight representative *Actinomycetota*.

| *Rhodococcus jostii* RHA1 | | |
|---|---|---|
| Genome accession | GCF_000014565.1 | |
| | Gene Identifier (Locus Tag) | Protein Identifier (RefSeq) |
| *Mycothiol S-transferases (MST)* | RHA1_RS14880 | WP_009475824.1 |
| *Mycothiol S-conjugate amidase (Mca)* | RHA1_RS28575 | WP_011597950.1 |
| *Mycothiol glycosyltransferase (MshA)* | RHA1_RS10115 | WP_011594914.1 |
| *GlcNAc-Ins deacetylase (MshB)* | RHA1_RS28995 | WP_011598013.1 |
| *GlcN-Ins ligase (MshC)* | RHA1_RS04190 | WP_011594098.1 |
| *Mycothiol acetyltransferase (MshD)* | RHA1_RS23745 | WP_011597161.1 |

| *Mycobacterium tuberculosis* H37Rv | | |
|---|---|---|
| Genome accession | GCF_000195955.2 | |
| | Gene Identifier (Locus Tag) | Protein Identifier (RefSeq) |
| *Mycothiol S-transferases (MST)* | Rv0443 | CCP43174.1 |
| *Mycothiol S-conjugate amidase (Mca)* | Rv1082 [44] | CCP43833.1 |
| *Mycothiol glycosyltransferase (MshA)* | Rv0486 [45] | CCP43220.1 |
| *GlcNAc-Ins deacetylase (MshB)* | Rv1170 | CCP43926.1 |
| *GlcN-Ins ligase (MshC)* | Rv2130c | CCP44905.1 |
| *Mycothiol acetyltransferase (MshD)* | Rv0819 | CCP43567.1 |

| *Mycolicibacterium smegmatis* MC$^2$ 155 | | |
|---|---|---|
| Genome accession | GCF_000015005.1 | |
| | Gene Identifier (Locus Tag) | Protein Identifier (RefSeq) |
| *Mycothiol S-transferases (MST)* | MSMEG_RS04325 | WP_003892314.1 |
| *Mycothiol S-conjugate amidase (Mca)* | MSMEG_RS25365 | WP_003896662.1 |
| *Mycothiol glycosyltransferase (MshA)* | MSMEG_RS04550 | WP_011727296.1 |
| *GlcNAc-Ins deacetylase (MshB)* | MSMEG_RS24740 | WP_011730335.1 |
| *GlcN-Ins ligase (MshC)* | MSMEG_RS20270 | WP_011729628.1 |
| *Mycothiol acetyltransferase (MshD)* | MSMEG_RS27875 | WP_014878453.1 |

| *Mycobacterium dioxanotrophicus* PH-06 | | |
|---|---|---|
| Genome accession | GCF_002157835.1 | |
| | Gene Identifier (Locus Tag) | Protein Identifier (RefSeq) |
| *Mycothiol S-transferases (MST)* | BTO20_RS04775 | WP_087073828.1 |
| *Mycothiol S-conjugate amidase (Mca)* | BTO20_RS08225 | WP_087074893.1 |

| | | |
|---|---|---|
| *Mycothiol glycosyltransferase (MshA)* | BTO20_RS04990 | WP_087073896.1 |
| *GlcNAc-Ins deacetylase (MshB)* | BTO20_RS08885 | WP_087081834.1 |
| *GlcN-Ins ligase (MshC)* | BTO20_RS13925 | WP_087082060.1 |
| *Mycothiol acetyltransferase (MshD)* | BTO20_RS30600 | WP_087079624.1 |

| *Pseudonocardia dioxanivorans* CB1190 | | |
|---|---|---|
| Genome accession | GCF_000196675.2 | |
| | Gene Identifier (Locus Tag) | Protein Identifier (RefSeq) |
| *Mycothiol S-transferases (MST)* | PSED_RS18380 | WP_013675746.1 |
| *Mycothiol S-conjugate amidase (Mca)* | PSED_RS04845 | WP_013673135.1 |
| *Mycothiol glycosyltransferase (MshA)* | PSED_RS28370 | WP_013677672.1 |
| *GlcNAc-Ins deacetylase (MshB)* | PSED_RS05150 | WP_013673196.1 |
| *GlcN-Ins ligase (MshC)* | PSED_RS16765 | WP_013675440.1 |
| *Mycothiol acetyltransferase (MshD)* | PSED_RS02365 | WP_013672669.1 |

| *Kocuria rhizophila* DC2201 | | |
|---|---|---|
| Genome accession | GCF_000010285.1 | |
| | Gene Identifier (Locus Tag) | Protein Identifier (RefSeq) |
| *Mycothiol S-transferases (MST)* | KRH_RS10695 | WP_012399228.1 |
| *Mycothiol S-conjugate amidase (Mca)* | KRH_RS08700 | WP_012398837.1 |
| *Mycothiol glycosyltransferase (MshA)* | KRH_RS03205 | WP_012397734.1 |
| *GlcNAc-Ins deacetylase (MshB)* | KRH_RS03970 | WP_041297315.1 |
| *GlcN-Ins ligase (MshC)* | KRH_RS06820 | WP_012398460.1 |
| *Mycothiol acetyltransferase (MshD)* | KRH_RS02755 | WP_012397644.1 |

| *Streptomyces griseus* subsp. *griseus* NBRC 13350 | | |
|---|---|---|
| Genome accession | GCF_000010605.1 | |
| | Gene Identifier (Locus Tag) | Protein Identifier (RefSeq) |
| *Mycothiol S-transferases (MST)* | SGR_RS34255 | WP_012382273.1 |
| *Mycothiol S-conjugate amidase (Mca)* | SGR_RS12630 | WP_012379296.1 |
| *Mycothiol glycosyltransferase (MshA)* | SGR_RS19815 | WP_042496821.1 |
| *GlcNAc-Ins deacetylase (MshB)* | SGR_RS11845 | WP_012379173.1 |
| *GlcN-Ins ligase (MshC)* | SGR_RS29140 | WP_012381568.1 |
| *Mycothiol acetyltransferase (MshD)* | SGR_RS19560 | WP_012380250.1 |

| *Kineococcus radiotolerans* SRS30216 | | |
|---|---|---|
| Genome accession | GCF_000017305.1 | |
| | Gene Identifier (Locus Tag) | Protein Identifier (RefSeq) |

| | | |
|---|---|---|
| *Mycothiol S-transferases (MST)* | KRAD_RS15915 | WP_012086671.1 |
| *Mycothiol S-conjugate amidase (Mca)* | KRAD_RS05600 | WP_041291922.1 |
| *Mycothiol glycosyltransferase (MshA)* | KRAD_RS06675 | WP_012084772.1 |
| *GlcNAc-Ins deacetylase (MshB)* | KRAD_RS05430 | WP_012084524.1 |
| *GlcN-Ins ligase (MshC)* | KRAD_RS01885 | WP_011981539.1 |
| *Mycothiol acetyltransferase (MshD)* | KRAD_RS06730 | WP_012084784.1 |

**Table S10.** Characteristics of MST homologs from representative *Actinomycetota*, using *M.tb.* H37Rv as a model strain

| *MST*-contained strain | Locus Tags | Amino Acid Residues | Molecular Mass (kDa) | Amino Acid Identity (As compared to H37Rv sequence) | Functional Domains/Motifs |
|---|---|---|---|---|---|
| *Rhodococcus jostii* RHA1 | RHA1_ RS14880 | 168 | 18.58 | 52.50% | IPR007061 |
| *Mycobacterium tuberculosis* H37Rv | Rv0443 | 171 | 18.96 | - | IPR007061 IPR027369 |
| *Mycobacterium smegmatis* MC$^2$155 | MSMEG_ RS04325 | 185 | 20.52 | 76.83% | IPR007061 |
| *Mycobacterium dioxanotrophicus* sp. PH-06 | BTO20_ RS04775 | 172 | 19.13 | 76.69% | IPR007061 |
| *Pseudonocardia dioxanivorans* CB1190 | PSED_ RS18380 | 175 | 18.86 | 52.47% | - |
| *Kocuria rhizophila* DC2201 | KRH_ RS10695 | 173 | 18.58 | 45.12% | IPR007061 |
| *Streptomyces griseus* subsp. griseus NBRC 13350 | SGR_ RS34255 | 169 | 18.04 | 45.62% | - |
| *Kineococcus radiotolerans* SRS30216 | KRAD_ RS15915 | 168 | 18.20 | 48.75% | IPR007061 |

**Table S11.** Characteristics of Mca homologs from representative *Actinomycetota*, using *M.tb.* H37Rv as a model strain

| *Mca*-contained strain | Locus Tags | Amino Acid Residues | Molecular Mass (kDa) | Amino Acid Identity (As compared to H37Rv sequence) | Functional Domains/Motifs |
|---|---|---|---|---|---|
| *Rhodococcus jostii* RHA1 | RHA1_ RS28575 | 292 | 32.94 | 69.58% | IPR017811 IPR003737 |
| *Mycobacterium tuberculosis* H37Rv | Rv1082 | 288 | 32.74 | - | IPR017811 IPR003737 IPR024078 |
| *Mycobacterium smegmatis* MC$^2$155 | MSMEG_ RS25365 | 288 | 32.68 | 78.12% | IPR017811 IPR003737 |
| *Mycobacterium dioxanotrophicus* sp. PH-06 | BTO20_ RS08225 | 288 | 32.61 | 78.75% | IPR017811 IPR003737 |
| *Pseudonocardia dioxanivorans* CB1190 | PSED_ RS04845 | 310 | 34.62 | 58.74% | IPR017811 IPR003737 |
| *Kocuria rhizophila* DC2201 | KRH_ RS08700 | 303 | 33.80 | 52.72% | IPR017811 IPR003737 |
| *Streptomyces griseus* subsp. griseus NBRC 13350 | SGR_ RS12630 | 293 | 32.98 | 53.31% | IPR017811 IPR003737 |
| *Kineococcus radiotolerans* SRS30216 | KRAD_ RS05600 | 298 | 33.43 | 57.14% | IPR017811 IPR003737 |

**Table S12.** Characteristics of MshA homologs from representative *Actinomycetota*, using *M.tb.* H37Rv as a model strain

| *MshA*-contained strain | Locus Tags | Amino Acid Residues | Molecular Mass (kDa) | Amino Acid Identity (As compared to H37Rv sequence) | Functional Domains/Motifs |
|---|---|---|---|---|---|
| *Rhodococcus jostii* RHA1 | RHA1_RS10115 | 452 | 48.25 | 69.71% | IPR017814 |
| *Mycobacterium tuberculosis* H37Rv | Rv0486 | 480 | 50.55 | - | IPR017814 IPR001296 IPR028098 |
| *Mycobacterium smegmatis* MC$^2$155 | MSMEG_RS04550 | 434 | 45.94 | 78.93% | IPR017814 |
| *Mycobacterium dioxanotrophicus* sp. PH-06 | BTO20_RS04990 | 444 | 47.23 | 78.99% | IPR017814 |
| *Pseudonocardia dioxanivorans* CB1190 | PSED_RS28370 | 457 | 48.36 | 63.61% | IPR017814 |
| *Kocuria rhizophila* DC2201 | KRH_RS03205 | 446 | 47.26 | 38.95% | IPR050194 |
| *Streptomyces griseus* subsp. griseus NBRC 13350 | SGR_RS19815 | 463 | 48.95 | 56.17% | IPR017814 |
| *Kineococcus radiotolerans* SRS30216 | KRAD_RS06675 | 435 | 46.12 | 57.88% | IPR017814 |

**Table S13.** Characteristics of MshB homologs from representative *Actinomycetota*, using *M.tb.* H37Rv as a model strain

| *MshB*-contained strain | Locus Tags | Amino Acid Residues | Molecular Mass (kDa) | Amino Acid Identity (As compared to H37Rv sequence) | Functional Domains/Motifs |
|---|---|---|---|---|---|
| *Rhodococcus jostii* RHA1 | RHA1_ RS28995 | 292 | 30.78 | 57.62% | IPR017810 IPR003737 |
| *Mycobacterium tuberculosis* H37Rv | Rv1170 | 303 | 31.75 | - | IPR017810 IPR003737 IPR024078 |
| *Mycobacterium smegmatis* MC$^2$ 155 | MSMEG_ RS24740 | 290 | 30.82 | 66.33% | IPR017810 IPR003737 |
| *Mycobacterium dioxanotrophicus* sp. PH-06 | BTO20_ RS08885 | 288 | 30.77 | 62.71% | IPR017810 IPR003737 |
| *Pseudonocardia dioxanivorans* CB1190 | PSED_ RS05150 | 291 | 30.19 | 43.73% | IPR017810 IPR003737 |
| *Kocuria rhizophila* DC2201 | KRH_ RS03970 | 277 | 28.86 | 35.97% | IPR003737 |
| *Streptomyces griseus* subsp. griseus NBRC 13350 | SGR_ RS11845 | 321 | 33.41 | 45.60% | IPR017810 IPR003737 |
| *Kineococcus radiotolerans* SRS30216 | KRAD_ RS05430 | 361 | 37.75 | 41.90% | IPR017810 IPR003737 |

**Table S14.** Characteristics of MshC homologs from representative *Actinomycetota*, using *M.tb.* H37Rv as a model strain

| *MshC*-contained strain | Locus Tags | Amino Acid Residues | Molecular Mass (kDa) | Amino Acid Identity (As compared to H37Rv sequence) | Functional Domains/Motifs |
|---|---|---|---|---|---|
| *Rhodococcus jostii* RHA1 | RHA1_ RS04190 | 415 | 45.82 | 68.60% | IPR017812 IPR024909 |
| *Mycobacterium tuberculosis* H37Rv | Rv2130c | 414 | 45.60 | - | IPR017812 IPR024909 IPR014729 |
| *Mycobacterium smegmatis* MC$^2$ 155 | MSMEG_ RS20270 | 412 | 45.40 | 78.99% | IPR017812 IPR024909 |
| *Mycobacterium dioxanotrophicus* sp. PH-06 | BTO20_ RS13925 | 412 | 44.97 | 78.99% | IPR017812 IPR024909 |
| *Pseudonocardia dioxanivorans* CB1190 | PSED_ RS16765 | 412 | 45.14 | 63.53% | IPR017812 IPR024909 |
| *Kocuria rhizophila* DC2201 | KRH_ RS06820 | 438 | 47.11 | 50.25% | IPR017812 IPR024909 |
| *Streptomyces griseus* subsp. griseus NBRC 13350 | SGR_ RS29140 | 409 | 44.33 | 57.35% | IPR017812 IPR024909 |
| *Kineococcus radiotolerans* SRS30216 | KRAD_ RS01885 | 419 | 44.73 | 55.74% | IPR017812 IPR024909 |

**Table S15.** Characteristics of MshD homologs from representative *Actinomycetota*, using *M.tb.* H37Rv as a model strain

| *MshD*-contained strain | Locus Tags | Amino Acid Residues | Molecular Mass (kDa) | Amino Acid Identity (As compared to H37Rv sequence) | Functional Domains/Motifs |
|---|---|---|---|---|---|
| *Rhodococcus jostii* RHA1 | RHA1_ RS23745 | 305 | 32.8 | 47.77% | IPR017813 IPR050276 |
| *Mycobacterium tuberculosis* H37Rv | Rv0819 | 315 | 33.6 | - | IPR017813 IPR050276 IPR016181 IPR000182 |
| *Mycobacterium smegmatis* MC$^2$155 | MSMEG_ RS27875 | 295 | 31.9 | 59.61% | IPR017813 IPR050276 |
| *Mycobacterium dioxanotrophicus* sp. PH-06 | BTO20_ RS30600 | 330 | 35.22 | 53.39% | IPR017813 IPR050276 |
| *Pseudonocardia dioxanivorans* CB1190 | PSED_ RS02365 | 300 | 31.93 | 49.51% | IPR017813 IPR050832 |
| *Kocuria rhizophila* DC2201 | KRH_ RS02755 | 303 | 32.59 | 41.08% | IPR017813 |
| *Streptomyces griseus* subsp. griseus NBRC 13350 | SGR_ RS19560 | 307 | 33.19 | 44.77% | IPR017813 IPR050832 |
| *Kineococcus radiotolerans* SRS30216 | KRAD_ RS06730 | 303 | 31.42 | 40.08% | IPR017813 IPR050832 |

**Table S16.** Proposed catalytic roles of conserved residues in MST.

| Conserved Residues (RHA1) | Function [44] |
|---|---|
| Asn39 | Carbonyl O accepts H-bond from GlcN-OH; Backbone NH donates a H-bond contact to the GlcN part; two contacts make a bidentate clamp with GlcN hydroxyls and “clip” to maintain a fixed sugar in place, ensuring the thiol is oriented toward the metal |
| Gly87 | Backbone NH donates a H-bond contact to the GlcN part; locks GlcN ring and reinforces Asn 39’s clamp |
| Asp150 | Ligates metal ions; one O forms an H-bond to the CyS amide, and pre-organize MSH for S-transfer |
| Gln157 | Backbone NH donates a H-bond contact to the Ins ring; further stabilizes distal end of MSH |
| His47-Asp150-His154 | Three metal chelators; by immobilizing $Zn^{2+}$ or other metal ions, the His47-Asp150-His154 rigid tripod help position the S and the adjacent carbonyl of Mycothiol; it activates the sulfur and aim it at a nucleophilic angle toward the incoming xenotoxins |
| Tyr86 and Trp134 | Conserved Tyr 86 (97%) / Phe 86 (3%); Upon MSH binding, Tyr86 adopts a rotamer shift of the phenolic ring towards Trp134 ~5 Å closer; Trp134 side chain also swings towards Tyr86 once binding; the flip triggers the opening of the second substrate (xenotoxins) pocket and serves as hydrophobic gate for xenotoxins |

**Table S17.** Proposed catalytic roles of conserved residues in Mca.

| Conserved Residues in Mca (RHA1) | Function |
|---|---|
| His12-Asp15-His142 | Similar function as His47-Asp150-His154 tripod in MST; metal chelators formed His12-Asp15-His142-metal that immobilize $Zn^{2+}$ or other metal ions to form the tripod that 1) fixes the position of metal ions; 2) polarizes the scissile amide carbonyl of the conjugate substrate by activation via metal ion, and 3) helps to position the metal ion-bound water ligand for nucleophilic attack. [22, 46, 47] |
| Asp14 | Act as base in hydrolysis; its carboxylate abstracts a proton from $Zn^{2+}$-water and generates the hydroxide that attacks amide linking AcCyS and GlcN; after generation of AcCyS-R, the His139 helps donate the proton to the leaving amine and forms GlcN-Ins. [22, 46, 47] |
| His139 | It acts as electrophile, with its positively charged Nε form a hydrogen bond to the carbonyl moiety of the AcCyS. This stabilizes the emerging oxyanion in the tetrahedral intermediate (His12-Asp15-His142-metal ion tripod; Asp14 base; His139/Asp141 proton donating pair, Tyr137 oxyanion clamp); after nucleophilic attack by the $Zn^{2+}$-activated hydroxide, His139 transfers its proton to the leaving amine of GlcN-Ins and facilitate C–N bond cleavage [22, 46, 48, 49] |
| Asp141 | It forms a hydrogen bond to His139; this interaction enables His139 to oxyanion stabilization on the cysteine carbonyl moiety via building His-Asp charge-relay to stabilize its protonated state; help lower the histidine's $pK_a$ [22] |
| Tyr137 | The side chain rotates inwards when MS-R docks; the phenolic Oη forms a hydrogen bond that quenches negative charge on the carbonyl when metal ion activated water attacks the carbonyl [49] |
| The metal ion tripod, the Asp14 base, His139/Asp141 proton donating pair, and Tyr137 constitute system that enable amide scission by Mca | |
| Asp87 | Its carboxylate potentially forms hydrogen bonds with axial hydroxyls on the GlcN ring and help clamp the sugar moiety of MS-R [47] |
| Arg65 | Strictly conserved in MshB and Mca; Arg65's Nη1/Nη2 forms H-bond network with axial hydroxyls with the GlcN moiety; the interaction, together with Asp87, fixes the position the glucosamine ring [47, 48] |
| Lys19 | Unique in Mca replacing the Ser/Thr in other deacetylases (MshB); the cationic ammonium side chain can occupy the |

| | |
|---|---|
| | pocket and exclude the GlcNAc ring sterically. In comparison MshB without Lys19 will accommodate GlcNAc-Ins and perform deacetylase activity [48] |

**Table S18.** Positions of conserved catalytic residue in MST homologs from eight representative *Actinomycetota.*

| | AcCyS Binding | GlcN Binding | | Ins Binding | Tripod Forming w/ Metal | | | Flip when Binding | |
|---|---|---|---|---|---|---|---|---|---|
| *Mycobacterium tuberculosis* H37Rv | Asp155 | Asn44 | Gly92 | Gln162 | His52 | Asp155 | His159 | Tyr91 | Trp139 |
| *Rhodococcus jostii* RHA1 | Asp150 | Asn39 | Gly87 | Gln157 | His47 | Asp150 | His154 | Tyr86 | Trp134 |
| *Mycobacterium smegmatis* $MC^2155$ | Asp168 | Asn57 | Gly105 | Gln175 | His65 | Asp168 | His172 | Tyr104 | Trp152 |
| *Mycobacterium dioxanotrophicus* PH-06 | Asp156 | Asn45 | Gly93 | Gln163 | His53 | Asp156 | His160 | Tyr92 | Trp140 |
| *Pseudonocardia dioxanivorans* CB1190 | Asp152 | Asn39 | Gly89 | Gln159 | His47 | Asp152 | His156 | Tyr88 | Trp136 |
| *Kocuria rhizophila* DC2201 | Asp151 | Asn39 | Gly87 | Gln158 | His47 | Asp151 | His155 | Phe86 | Trp135 |
| *Streptomyces griseus* subsp. griseus NBRC13350 | Asp151 | Asn39 | Gly87 | Gln158 | His47 | Asp151 | His155 | Tyr86 | Trp135 |
| *Kineococcus radiotolerans* SRS30216 | Asp150 | Asn39 | Gly87 | Gln157 | His47 | Asp150 | His154 | Tyr86 | Trp134 |

**Table S19.** Positions of conserved catalytic residue in Mca homologs from eight representative *Actinomycetota.*

| | Tetrahedral Intermediate Formation | | | | | | |
|---|---|---|---|---|---|---|---|
| | Tripod forming W/ metal ion | | | Base | Acid | His-Asp "Charge-rely" | Oxyanion stabilize |
| *Mycobacterium tuberculosis* H37Rv (MshB, Rv1170) | His13 | Asp16 | His147 | Asp15 | His144 | Asp146 | Tyr142 |
| *Mycobacterium tuberculosis* H37Rv | His12 | Asp15 | His142 | Asp14 | His139 | Asp141 | Tyr137 |
| *Rhodococcus jostii* RHA1 | His12 | Asp15 | His142 | Asp14 | His139 | Asp141 | Tyr137 |
| *Mycobacterium smegmatis* MC$^2$155 | His12 | Asp15 | His142 | Asp14 | His139 | Asp141 | Tyr137 |
| *Mycobacterium dioxanotrophicus* PH-06 | His12 | Asp15 | His142 | Asp14 | His139 | Asp141 | Tyr137 |
| *Pseudonocardia dioxanivorans* CB1190 | His25 | Asp28 | His155 | Asp27 | His152 | Asp154 | Tyr150 |
| *Kocuria rhizophila* DC2201 | His15 | Asp18 | His146 | Asp17 | His143 | Asp145 | Tyr141 |
| *Streptomyces griseus* subsp. Griseus NBRC13350 | His13 | Asp16 | His144 | Asp15 | His141 | Asp143 | Tyr139 |
| *Kineococcus radiotolerans* SRS30216 | His13 | Asp16 | His144 | Asp15 | His141 | Asp143 | Tyr139 |

| | Sugar ring binding | | Mca specific |
|---|---|---|---|
| *Mycobacterium tuberculosis* H37Rv (MshB, Rv1170) | Asp95 | Arg68 | Lys19 |
| *Mycobacterium tuberculosis* H37Rv | Asp87 | Arg65 | Lys19 |
| *Rhodococcus jostii* RHA1 | Asp87 | Arg65 | Lys19 |
| *Mycobacterium smegmatis* MC$^2$155 | Asp87 | Arg65 | Lys19 |
| *Mycobacterium dioxanotrophicus* PH-06 | Asp87 | Arg65 | Lys19 |
| *Pseudonocardia dioxanivorans* CB1190 | Asp91 | Arg78 | Lys32 |
| *Kocuria rhizophila* DC2201 | Asp100 | Arg69 | Lys22 |
| *Streptomyces griseus* subsp. Griseus NBRC13350 | Asp89 | Arg67 | Lys20 |
| *Kineococcus radiotolerans* SRS30216 | Asp89 | Arg67 | Lys20 |

**Table S20.** Information of the six WWTPs for activate sludge collection.

| WWTP | Service Area | Service Population | Design Capacity (Million Gallons per Day/MGD) | Average Influent (Million Gallons per Day/MGD) |
|---|---|---|---|---|
| WWTP-P | 150 mi² across 48 municipalities | ~1.5 million | 330 | 226 |
| Influent Sources | Municipal (residential + commercial) & industrial wastewater; land-fill leachate and other special wastes | | | |
| WWTP-R | 48.85 mi² across 11 municipalities | ~250,000 residents; ~3000 industrial/ commercial customers | 40 | 25-30 |
| Influent Sources | Mixed residential and pre-treated industrial/commercial wastewater | | | |
| WWTP-L | 13 mi² from two cities of New Jersey | ~60,000 | 20 | 10-15 |
| Influent Sources | Residential; some industrial wastewater | | | |
| WWTP-W | 18.8 mi² from parts of two boroughs | ~1 million | 275 | 275 |
| Influent Sources | Residential, commercial, industrial wastewater; stormwater runoff | | | |
| WWTP-BH | ~6.2 mi² from a township | ~13,500 | 3.1 | 1.5-1.8 |
| Influent Sources | Primarily residential; some commercial | | | |
| WWTP-V | ~2.79 mi² from a township | ~14,500 | 3.0 | 2-3 |
| Influent Sources | Primarily residential; minor commercial | | | |

**Table S21.** Search parameters for molecule docking predictions with AutoDock 4.2.

| Search Method | Lamarckian Genetic Algorithm (LGA) |
|---|---|
| ga_run | 70 |
| ga_pop_size | 300 |
| ga_num_evals | $2.5\times 10^7$ |
| ga_num_generations | 50000 |
| elite | 1 |
| ga_crossover_rate | 0.8 |
| ga_mutation_rate | 0.02 |
| ls_search_rate | 0.06 |
| grid_center | -4.494, -2.605, -7.612 |
| torsdof | 16 |
| sw_max_succ | 4 |
| sw_max_fail | 4 |

**Table S22.** LODs and LOQs for the three calibration standards used for semi-quantification.

| | LOD (*ppb*) | LOQ (*ppb*) |
|---|---|---|
| GS-mBBr | 0.082 | 0.25 |
| 5:3 FTCA | 0.066 | 0.20 |
| PFBA | 0.057 | 0.17 |

**Table S23.** LC instrument conditions and gradient for target analysis by LC/MS/MS.

| 1290 Infinity II LC System (Symmetry C18 column) | | | | |
|---|---|---|---|---|
| Column Temp: 50°C | Inject Volume: 10μL | Max. Pressure: 750 bars | | |
| | Time | Solvent A (%) | Solvent B (%) | Flow Rate (mL/min) |
| Run | 0 | 95 | 5 | 0.30 |
| | 0.5 | 95 | 5 | 0.30 |
| | 3 | 60 | 40 | 0.35 |
| | 8 | 10 | 90 | 0.35 |
| | 9 | 10 | 90 | 0.35 |
| | 9.5 | 95 | 5 | 0.35 |
| Post-run | 11.5 | 95 | 5 | 0.30 |

**Table S24.** Multiple reaction monitoring (MRM) settings for target analysis by LC/MS/MS.

| Analyte | Precursor Ion (*m/z*) | Product Ion (*m/z*) | Collision Energy (V) | Fragmentor (V) | Cell Accelerator Voltage (V) |
|---|---|---|---|---|---|
| M8PFOA | 421.0 | 376.0 | 8 | 64 | 2 |
| M2-6:2 FTCA | 379.0 | 294.0 | 12 | 44 | 2 |
| 6:2 FTCA | 377.0 | 293.0 | 12 | 48 | 2 |
| PFHpA | 363.0 | 318.9 | 8 | 72 | 2 |
| 6:2 FTUCA | 357.0 | 292.9 | 18 | 32 | 2 |
| 5:3 FTCA | 341.0 | 237.0 | 14 | 28 | 2 |
| PFHxA | 313.0 | 269.0 | 24 | 56 | 2 |
| PFPeA | 263.0 | 219.0 | 4 | 76 | 2 |
| PFBA | 213.0 | 168.9 | 4 | 42 | 2 |

### References:


1. Barzen-Hanson, K.A., et al., *Discovery of 40 Classes of Per- and Polyfluoroalkyl Substances in Historical Aqueous Film-Forming Foams (AFFFs) and AFFF-Impacted Groundwater.* Environmental Science & Technology, 2017. **51**(4): p. 2047-2057.
2. Wu, C., et al., *Rapid quantitative analysis and suspect screening of per-and polyfluorinated alkyl substances (PFASs) in aqueous film-forming foams (AFFFs) and municipal wastewater samples by Nano-ESI-HRMS.* Water Research, 2022. **219**: p. 118542.
3. Backe, W.J., T.C. Day, and J.A. Field, *Zwitterionic, Cationic, and Anionic Fluorinated Chemicals in Aqueous Film Forming Foam Formulations and Groundwater from U.S. Military Bases by Nonaqueous Large-Volume Injection HPLC-MS/MS.* Environmental Science & Technology, 2013. **47**(10): p. 5226-5234.
4. Chen, H., et al., *Occurrence and Distribution of Per- and Polyfluoroalkyl Substances in Tianjin, China: The Contribution of Emerging and Unknown Analogues.* Environmental Science & Technology, 2020. **54**(22): p. 14254-14264.
5. Liu, Y., et al., *High-resolution mass spectrometry (HRMS) methods for nontarget discovery and characterization of poly- and per-fluoroalkyl substances (PFASs) in environmental and human samples.* TrAC Trends in Analytical Chemistry, 2019. **121**: p. 115420.
6. Washington, J.W., et al., *Nontargeted mass-spectral detection of chloroperfluoropolyether carboxylates in New Jersey soils.* Science, 2020. **368**(6495): p. 1103-1107.
7. Munoz, G., et al., *Target and Nontarget Screening of PFAS in Biosolids, Composts, and Other Organic Waste Products for Land Application in France.* Environmental Science & Technology, 2022. **56**(10): p. 6056-6068.
8. Schymanski, E.L., et al., *Identifying Small Molecules via High Resolution Mass Spectrometry: Communicating Confidence.* Environmental Science & Technology, 2014. **48**(4): p. 2097-2098.
9. Charbonnet, J.A., et al., *Communicating Confidence of Per- and Polyfluoroalkyl Substance Identification via High-Resolution Mass Spectrometry.* Environmental Science & Technology Letters, 2022. **9**(6): p. 473-481.
10. Allred, B.M., et al., *Orthogonal zirconium diol/C18 liquid chromatography-tandem mass spectrometry analysis of poly and perfluoroalkyl substances in landfill leachate.* J Chromatogr A, 2014. **1359**: p. 202-11.
11. Jacob, P., K.A. Barzen-Hanson, and D.E. Helbling, *Target and Nontarget Analysis of Per- and Polyfluoralkyl Substances in Wastewater from Electronics Fabrication Facilities.* Environmental Science & Technology, 2021. **55**(4): p. 2346-2356.
12. Wu, C., et al., *Distinctive biotransformation and biodefluorination of 6:2 versus 5:3 fluorotelomer carboxylic acids by municipal activated sludge.* Water Research, 2024. **254**: p. 121431.
13. Kruve, A., *Strategies for Drawing Quantitative Conclusions from Nontargeted Liquid Chromatography–High-Resolution Mass Spectrometry Analysis.* Analytical Chemistry, 2020. **92**(7): p. 4691-4699.
14. Wilm, M. and M. Mann, *Analytical Properties of the Nanoelectrospray Ion Source.* Analytical Chemistry, 1996. **68**(1): p. 1-8.
15. Pieke, E.N., et al., *A framework to estimate concentrations of potentially unknown substances by semi-quantification in liquid chromatography electrospray ionization mass spectrometry.* Analytica Chimica Acta, 2017. **975**: p. 30-41.

16. Cioni, L., et al., *Fluorine Mass Balance, including Total Fluorine, Extractable Organic Fluorine, Oxidizable Precursors, and Target Per- and Polyfluoroalkyl Substances, in Pooled Human Serum from the Tromsø Population in 1986, 2007, and 2015.* Environmental Science & Technology, 2023. **57**(40): p. 14849-14860.
17. Carugo, O. and S. Pongor, *A normalized root-mean-square distance for comparing protein three-dimensional structures.* Protein Sci, 2001. **10**(7): p. 1470-3.
18. Young, R.B., et al., *PFAS Analysis with Ultrahigh Resolution 21T FT-ICR MS: Suspect and Nontargeted Screening with Unrivaled Mass Resolving Power and Accuracy.* Environmental Science & Technology, 2022. **56**(4): p. 2455-2465.
19. Fahey, R.C. and G.L. Newton, *Determination of low-molecular-weight thiols using monobromobimane fluorescent labeling and high-performance liquid chromatography*, in *Methods in Enzymology*. 1987, Academic Press. p. 85-96.
20. Holsclaw, C.M., et al., *Mass Spectrometric Analysis of Mycothiol levels in Wild-Type and Mycothiol Disulfide Reductase Mutant Mycobacterium smegmatis.* Int J Mass Spectrom, 2011. **305**(2-3): p. 151-156.
21. Alfaro, C.M., et al., *Investigations of Analyte-Specific Response Saturation and Dynamic Range Limitations in Atmospheric Pressure Ionization Mass Spectrometry.* Analytical Chemistry, 2014. **86**(21): p. 10639-10645.
22. Newton, G.L., N. Buchmeier, and R.C. Fahey, *Biosynthesis and functions of mycothiol, the unique protective thiol of Actinobacteria.* Microbiol Mol Biol Rev, 2008. **72**(3): p. 471-94.
23. Rawat, M. and Y. Av-Gay, *Mycothiol-dependent proteins in actinomycetes.* FEMS Microbiol Rev, 2007. **31**(3): p. 278-92.
24. Newton, G.L. and R.C. Fahey, *Mycothiol biochemistry.* Archives of microbiology, 2002. **178**: p. 388-394.
25. Newton, G.L., et al., *The Structure of U17 Isolated from Streptomyces clavuligerus and its Properties as an Antioxidant Thiol.* European Journal of Biochemistry, 1995. **230**(2): p. 821-825.
26. Newton, G.L., Y. Av-Gay, and R.C. Fahey, *A Novel Mycothiol-Dependent Detoxification Pathway in Mycobacteria Involving Mycothiol S-Conjugate Amidase.* Biochemistry, 2000. **39**(35): p. 10739-10746.
27. Buchmeier, N.A., et al., *Association of mycothiol with protection of Mycobacterium tuberculosis from toxic oxidants and antibiotics.* Molecular Microbiology, 2003. **47**(6): p. 1723-1732.
28. Rawat, M., et al., *Mycothiol-deficient Mycobacterium smegmatis mutants are hypersensitive to alkylating agents, free radicals, and antibiotics.* Antimicrob Agents Chemother, 2002. **46**(11): p. 3348-55.
29. Dosanjh, M., G.L. Newton, and J. Davies, *Characterization of a mycothiol ligase mutant of Rhodococcus jostii RHA1.* Research in Microbiology, 2008. **159**(9): p. 643-650.
30. Masai, E., et al., *Characterization of biphenyl catabolic genes of gram-positive polychlorinated biphenyl degrader Rhodococcus sp. strain RHA1.* Appl Environ Microbiol, 1995. **61**(6): p. 2079-85.
31. Kim, D., et al., *Identification of a novel dioxygenase involved in metabolism of o-xylene, toluene, and ethylbenzene by Rhodococcus sp. strain DK17.* Appl Environ Microbiol, 2004. **70**(12): p. 7086-92.
32. Dávila Costa, J.S., et al., *Proteome analysis reveals differential expression of proteins involved in triacylglycerol accumulation by Rhodococcus jostii RHA1 after addition of methyl viologen.* Microbiology (Reading), 2017. **163**(3): p. 343-354.

33. Bentel, M.J., et al., *Defluorination of Per- and Polyfluoroalkyl Substances (PFASs) with Hydrated Electrons: Structural Dependence and Implications to PFAS Remediation and Management.* Environmental Science & Technology, 2019. **53**(7): p. 3718-3728.
34. Tseng, N., et al., *Biotransformation of 6:2 Fluorotelomer Alcohol (6:2 FTOH) by a Wood-Rotting Fungus.* Environmental Science & Technology, 2014. **48**(7): p. 4012-4020.
35. Zhang, H., et al., *Biotransformation of 6:2 fluorotelomer alcohol by the whole soybean (Glycine max L. Merrill) seedlings.* Environmental Pollution, 2020. **257**: p. 113513.
36. Bhardwaj, S., et al., *Biotransformation of 6:2/4:2 fluorotelomer alcohols by Dietzia aurantiaca J3: Enzymes and proteomics.* Journal of Hazardous Materials, 2024. **478**: p. 135510.
37. Fasano, W.J., et al., *Absorption, Distribution, Metabolism, and Elimination of 8-2 Fluorotelomer Alcohol in the Rat.* Toxicological Sciences, 2006. **91**(2): p. 341-355.
38. Fasano, W.J., et al., *Kinetics of 8-2 fluorotelomer alcohol and its metabolites, and liver glutathione status following daily oral dosing for 45 days in male and female rats.* Chemico-Biological Interactions, 2009. **180**(2): p. 281-295.
39. Martin, J.W., S.A. Mabury, and P.J. O'Brien, *Metabolic products and pathways of fluorotelomer alcohols in isolated rat hepatocytes.* Chemico-Biological Interactions, 2005. **155**(3): p. 165-180.
40. Nabb, D.L., et al., *In Vitro Metabolism of 8-2 Fluorotelomer Alcohol: Interspecies Comparisons and Metabolic Pathway Refinement.* Toxicological Sciences, 2007. **100**(2): p. 333-344.
41. Yu, Y., et al., *Electron bifurcation and fluoride efflux systems implicated in defluorination of perfluorinated unsaturated carboxylic acids by <i>Acetobacterium</i> spp.* Science Advances, 2024. **10**(29): p. eado2957.
42. Mothersole, R.G., et al., *Formation of CoA Adducts of Short-Chain Fluorinated Carboxylates Catalyzed by Acyl-CoA Synthetase from Gordonia sp. Strain NB4-1Y.* ACS Omega, 2023. **8**(42): p. 39437-39446.
43. Mothersole, R.G., et al., *Enzyme Catalyzed Formation of CoA Adducts of Fluorinated Hexanoic Acid Analogues using a Long-Chain acyl-CoA Synthetase from Gordonia sp. Strain NB4-1Y.* Biochemistry, 2024. **63**(17): p. 2153-2165.
44. Jayasinghe, Y.P., et al., *The Mycobacterium tuberculosis mycothiol S-transferase is divalent metal-dependent for mycothiol binding and transfer.* RSC Medicinal Chemistry, 2023. **14**(3): p. 491-500.
45. Newton, G.L., et al., *The glycosyltransferase gene encoding the enzyme catalyzing the first step of mycothiol biosynthesis (mshA).* J Bacteriol, 2003. **185**(11): p. 3476-9.
46. Fan, F., et al., *Structures and mechanisms of the mycothiol biosynthetic enzymes.* Curr Opin Chem Biol, 2009. **13**(4): p. 451-9.
47. Metaferia, B.B., et al., *Synthesis of Natural Product-Inspired Inhibitors of Mycobacterium tuberculosis Mycothiol-Associated Enzymes: The First Inhibitors of GlcNAc-Ins Deacetylase.* Journal of Medicinal Chemistry, 2007. **50**(25): p. 6326-6336.
48. Maynes, J.T., et al., *The crystal structure of 1-D-myo-inosityl 2-acetamido-2-deoxy-alpha-D-glucopyranoside deacetylase (MshB) from Mycobacterium tuberculosis reveals a zinc hydrolase with a lactate dehydrogenase fold.* J Biol Chem, 2003. **278**(47): p. 47166-70.
49. Lamprecht, D.A., et al., *An enzyme-initiated Smiles rearrangement enables the development of an assay of MshB, the GlcNAc-Ins deacetylase of mycothiol biosynthesis.* Org Biomol Chem, 2012. **10**(27): p. 5278-88.
